# A Total-Evidence Dated Phylogeny of Echinoids and the Evolution of Body Size across Adaptive Landscape

**DOI:** 10.1101/2020.02.13.947796

**Authors:** Nicolás Mongiardino Koch, Jeffrey R. Thompson

**Affiliations:** Department of Geology & Geophysics, Yale University. 210 Whitney Ave., New Haven, CT 06511, USA; Research Department of Genetics, Evolution and Environment, University College London, Darwin Building, Gower Street, London WC1E 6BT, UK

**Keywords:** Echinoidea, phylogenomics, total evidence, time calibration, fossils, body size, macroevolution, adaptive landscape

## Abstract

Several unique properties of echinoids (sea urchins) make them useful for exploring macroevolutionary dynamics, including their remarkable fossil record that can be incorporated into explicit phylogenetic hypotheses. However, this potential cannot be exploited without a robust resolution of the echinoid tree of life. We revisit the phylogeny of crown group Echinoidea using both the largest phylogenomic dataset compiled for the clade, as well as a large-scale morphological matrix with a dense fossil sampling. We also gather a new compendium of both tip and node age constraints, allowing us to combine phylogenomic, morphological and stratigraphic data using a total-evidence dating approach. For this, we develop a novel method for subsampling phylogenomic datasets that selects loci with high phylogenetic signal, low systematic biases and enhanced clock-like behavior. Our approach restructure much of the higher-level phylogeny of echinoids, and demonstrates that combining different data sources increases topological accuracy. We are able to resolve multiple alleged conflicts between molecular and morphological datasets, such as the position of Echinothurioida and Echinoneoida, as well as unravelling the relationships between sand dollars and their closest relatives. We then use this topology to trace the evolutionary history of echinoid body size through more than 270 million years, revealing a complex pattern of convergent evolution to stable peaks in macroevolutionary adaptive landscape. Our efforts show how combining phylogenomic and paleontological evidence offers new ways of exploring evolutionary forces operating across deep timescales.

---

Echinoidea, one of the five living classes of echinoderms, constitutes one of the most iconic clades of marine animals. A little over 1,000 species of echinoids live in today’s oceans (Kroh and Mooi 2019), including species commonly known as sea urchins, heart urchins and sand dollars. Echinoids are ubiquitous in benthic marine environments, occurring at all latitudes and depths (Emlet 2002; Schultz 2015). As the most important consumers in many shallow marine habitats, sea urchins have been recognized as keystone species and ecosystem engineers (Lawton and Jones 1995; Lessios et al. 2001; Steneck 2013). Their abundance and trophic interactions have a strong impact on the health and stability of marine communities like kelp forests (Harrold and Pearse 1987; Steneck et al. 2002), coral reefs (Carpenter 1981; Edmunds and Carpenter 2001) and seagrass meadows (Valentine and Heck 1991), being able to trigger ecosystem-wide shifts to low diversity stable states (Hughes 1994; Ling et al. 2015). Likewise, bioturbation associated with the feeding and burrowing activities of a large diversity of infaunal echinoids has a strong impact on the structure and function of marine sedimentary environments (Thayer 1983; Lohrer et al. 2004).

Sea urchins have a long independent evolutionary history, having diverged from their closest relatives—the sea cucumbers (Holothuroidea)—more than 450 million years ago (Smith 1984). Crown echinoids, on the other hand, likely originated during the Permian (Thompson et al. 2015) and diversified through the early Mesozoic in the aftermath of the Permo-Triassic (P-T) mass extinction (Erwin 1994; Twitchett and Oji 2005; Thompson et al. 2018). One of the most characteristic morphological features of echinoids is their robust globular skeleton, or test, composed of numerous calcium carbonate plates. This complex structure provides a wealth of morphological information that can be synthetized into morphological matrices encompassing both extant and extinct diversity (Suter 1994; Smith 2001; Stockley et al. 2005; Kroh and Smith 2010). The resilient echinoid test also underlies the clade’s high preservation potential (Kidwell and Baumiller 1990; Nebelsick 1996) and contributes to its outstanding fossil record, comprising more than 10,000 described species (Smith and Kroh 2013). With such an extraordinary fossil record, and the ability to place it in a rigorous phylogenetic framework, echinoids provide unparalleled opportunities for macroevolutionary research.

Several authors have used sea urchins to investigate evolutionary dynamics in deep time. Smith and Jeffery (1998) explored determinants of extinction selectivity across the Cretaceous-Paleogene mass extinction. Eble (2000) and Boivin et al. (2018) analyzed the interplay of morphological disparity and diversity across different lineages. Hopkins and Smith (2015) showed echinoid evolution to be punctuated by bursts of change linked to ecological innovations. Thompson et al. (2017) and Erkenbrack and Thompson (2019) capitalized on our profound understanding of echinoid development to explore the origin of developmental innovations. However, much of this research did not account for the phylogenetic relationships of taxa, complicating study of the potential drivers behind the observed patterns. Those investigations that incorporated a phylogenetic dimension were limited by topological uncertainty arising both from a lack of phylogenetic signal as well as conflicts between morphological and molecular data. Unlocking the true potential of sea urchins as a model clade for macroevolutionary research is thus contingent on the resolution of a robust time-calibrated phylogeny.

The modern classification of crown group Echinoidea can be traced back to Mortensen’s seminal series of monographs. His strong emphasis on the morphology of the echinoid test provided a means to classify fossil and extant lineages in a common scheme, an approach preserved by subsequent revisions (Durham and Melville 1957; Jensen 1981; Smith 1984; Kroh and Smith 2010; Kroh 2020). Among these, that of Kroh and Smith (2010) is especially relevant as it relied on the compilation of a morphological matrix covering most crown group families, both living and extinct. Although shedding new light on the phylogeny of the clade, this study also found multiple regions of the echinoid tree that attained poor levels of support, resolved differently depending on inference approach, or conflicted with molecular phylogenies.

Early molecular phylogenies of echinoids, based mostly on ribosomal genes (Littlewood and Smith 1995; Smith et al. 1995; Smith et al. 2006), were received as confirming previous morphological results (Smith 1997). In all of these, cidaroids were placed as the sister group to a clade containing all other crown group echinoids (Euechinoidea). Two major lineages, Echinacea (including model species used in developmental research) and Irregularia (containing all bilaterally symmetrical clades) were also recovered within Euechinoidea. However, the resolution of a few key regions of the echinoid tree of life systematically disagreed. Although initially disregarded as artifacts of poor sampling or low phylogenetic signal in molecular datasets, the consistent recovery of nodes not supported by morphology (Littlewood and Smith 1995; Smith et al. 1995; Smith et al. 2006; Smith and Kroh 2013; Thompson et al. 2017; Mongiardino Koch et al. 2018) eventually cast doubt on morphological topologies.

As summarized by Kroh and Smith (2010) and Smith and Kroh (2013), the incongruence between molecular and morphological datasets is largely concentrated in two regions of the echinoid tree of life. The first of these relates to the Echinothurioida, an enigmatic clade of venomous echinoids that inhabit almost exclusively the deep sea, and whose phylogenetic placement has been strongly debated since their discovery (Gregory 1897). More recently, echinothurioids were suggested to be the sister lineage to all other euechinoids (Smith 1981, 1984), a topology later recovered in some phylogenetic analyses (Smith et al. 2006; Kroh and Smith 2010). A key line of evidence supporting this placement is the flexible test of the echinothurioids, a condition also found among stem group echinoids (Kier 1977). Given the supposedly primitive nature of this condition, other unique characteristics of echinothurioids were also interpreted as plesiomorphic. Molecular evidence, on the other hand, supported a derived position of echinothurioids (Smith et al. 2006; Thompson et al. 2017; Mongiardino Koch et al. 2018), nested in a clade that includes lineages characterized by a unique dental morphology (aspidodiadematoids, diadematoids, pedinoids and micropygoids). This clade was also favored by some early systematists (Jackson 1912; Durham and Melville 1957), and has since been named Aulodonta (Kroh 2020). Under this alternative scenario, the evolution of test flexibility among echinothurioids, along with the other unique features of the clade, likely constitute autapomorphies (Mongiardino Koch et al. 2018). Given that the deepest euechinoid lineages likely diverged during the Triassic (Kier 1977; Smith et al. 2006; Thompson et al. 2017), conflicting solutions between morphological and ribosomal datasets are not surprising.

A second region of phylogenetic incongruence, relating to the position of sand dollars and sea biscuits, is much more problematic. Ever since the early 19^th^ century, these two groups have been recognized as sister clades and united within Clypeasteroida (*sensu* Agassiz 1873). Within this framework, clypeasteroids were thought to have originated during the Paleocene from within an assemblage of lineages known as “cassiduloids”, a grade with deep roots in the Mesozoic that includes three extant lineages (Cassiduloida, Echinolampadoida and Apatopygidae) and a large diversity of extinct forms. Numerous synapomorphies unite clypeasteroids to the exclusion of “cassiduloids” (e.g., extreme flattening and internal reinforcement of the test, multiplication of tube feet per plate, presence of an Aristotle’s lantern in adults; Mooi 1990b; Kroh and Smith 2010). Nonetheless, all previous molecular studies have yielded a paraphyletic Clypeasteroida (Littlewood and Smith 1995; Smith et al. 1995; Smith et al. 2006; Nowak et al. 2013; Smith and Kroh 2013; Thompson et al. 2017; Mongiardino Koch et al. 2018), with at least one extant “cassiduloid” nested inside the clade. This resolution strongly conflicts with morphological evidence, and implies substantially longer ghost ranges for clypeasteroids, a clade with a remarkable fossil record (Kroh and Smith 2010). It was therefore deemed too problematic and disregarded in favor of the morphological hypothesis (Kroh and Smith 2010; Smith and Kroh 2013). More recently, the strong support for a sister group relationship between echinolampadoids and sand dollars prompted Mongiardino Koch et al. (2018) to restrict Clypeasteroida to include only the sea biscuits (henceforth, the usage given to this name), and reclassify the sand dollars in a separate order (Scutelloida), a nomenclatural change followed by Kroh (2020). The implications of this topology for the position of the remaining “cassiduloids” remain unclear, however, as well as leaving open questions regarding the evolution of the many morphological features that conflict with it. Resolving these issues is indispensable for understanding the origin and early evolutionary history of the sand dollars, the most morphologically and ecologically specialized lineage of echinoids (Mooi 1990b; Smith 2001; Hopkins and Smith 2015).

Even though conflicts regarding the position of Echinothurioida and the relationships between sand dollars and their closest relatives have been recognized for several decades, few attempts have been made address them specifically. Recent molecular analyses have often failed to sample the taxa necessary to test these alternative hypotheses, while finding support for novel topologies that disagree in other ways with those based on morphology (Bronstein and Kroh 2019; Lin et al. 2019). For example, Lin et al. (2019) found Echinoneoida, a clade thought to represent the sister group to all other crown group irregulars (Smith 1981; Kroh and Smith 2010), as sister to Neognathostomata alone (“cassiduloids”, clypeasteroids and scutelloids), further increasing uncertainty among some of the oldest nodes in the echinoid tree of life.

The recent increase of available genomic data for echinoids provides a unique opportunity to explore the sources of these conflicts. Likewise, the development of tip-dated approaches to divergence time estimation offers novel ways to incorporate the rich paleontological record of echinoids into the process of phylogenetic inference. Although hailed as an early test of the molecular clock (Smith et al. 2006), estimation of echinoid divergence times has been little explored, and there has been no attempt to time-calibrate a phylogeny including extinct lineages. Here, we compile the largest phylogenomic dataset for echinoids and explore their higher-level phylogeny by combining molecular, morphological and stratigraphic data. We infer independent estimates of phylogeny from morphology and molecular data, as well as performing a total-evidence dated (TED) analysis including representatives of most families of crown group echinoids. In order to do so, we compile a new dataset of both tip and node dates from the literature and develop a novel pipeline to subsample loci with high phylogenetic content and low systematic biases. Finally, we explore the evolutionary history of echinoid body size as a proof of concept of the potential of the clade for macroevolutionary research.

## Materials & Methods

### Morphological Analyses

The morphological matrix employed was originally assembled by Kroh and Smith (2010). This comprehensive dataset, on which much of the classification of echinoids is based, was designed to sample at least one representative of all families and subfamilies of crown group echinoids (living and extinct). The characters chosen correspond to those traditionally considered diagnostic at or above the level of families/subfamilies. The version of the dataset used is that of Hopkins and Smith (2015), who excluded five rogue terminals, resulting in a final tally of 164 taxa. This sample includes at least one representative of 89.4% of families recognized in the World Echinoidea Database (WED; Kroh and Mooi 2019). *Archaeocidaris*, an unambiguous late stem group echinoid from the Carboniferous (Kroh and Smith 2010; Thompson et al. 2019), was used as outgroup. The matrix (available as SI File 1) contains 300 characters (after removing some that were rendered invariant by taxon exclusion), of which 16 were considered ordered by the authors and were treated as such here. Further details on taxon sampling can be found in Table S1 of SI File 2. Nomenclature follows that of the WED (Kroh and Mooi 2019), and clades above family level were updated to conform with Kroh (2020).

**Table 1:** Loadings for the first two PC axes of the gene properties dataset. Percentages shown in parentheses correspond to the variance explained by each dimension.

|  | PC 1 (47.7%) | PC 2 (20.4%) |
| --- | --- | --- |
| **Saturation** | 0.364 | -0.391 |
| **Comp. heterogeneity** | 0.339 | -0.221 |
| **Root-to-tip variance** | 0.391 | -0.427 |
| **Av. patristic distance** | 0.490 | -0.143 |
| **Average BS support** | 0.400 | 0.468 |
| Gene tree error | -0.345 | -0.549 |
| % invariant sites | -0.282 | -0.275 |

The original analysis of this dataset was performed under maximum parsimony (MP), with the topology obtained under successive weighting largely determining the proposed classification (Kroh and Smith 2010). Here, we reanalyze this dataset under both equal and implied weights (*k* = 6) parsimony using TNT v. 1.5 (Goloboff 1993; Goloboff and Catalano 2016). We used driven tree searches with ten initial replicates subject to new technology search heuristics. Searches were continued until minimum length was found 100 times, and TBR branch swapping was then performed on the topologies in memory. Support was evaluated using absolute node frequencies in 1,000 jackknifed replicates.

We also used MrBayes 3.2.6 (Ronquist et al. 2012) to analyze the morphological dataset under Bayesian inference (BI). We subdivided the matrix into four partitions depending on the number of states (two, three, four and five or more states). Partitioning by number of states has been found to be superior to unpartitioned approaches (Gavryushkina et al. 2017), but also increases the weights assigned to characters with more states, as these attain lower stationary frequencies (King et al. 2017). We believe that our approach represents a balance between these two phenomena. Rates of evolution were allowed to vary across partitions. Evolution within each partition was modeled using Mkv + Г models (Lewis 2001), which further accommodates differences in evolutionary rates within partitions and corrects for the ascertainment bias introduced by coding only variable characters.

Inference was performed using both uncalibrated and calibrated BI approaches, in the latter case enforcing a morphological clock. Time-calibration was performed using a newly gathered dataset of both tip and node dates (SI File 3). Combining constraints on both tip and node ages has been found to outperform the use of either approach individually (O’Reilly and Donoghue 2016; Kealy and Beck 2017), and is particularly useful in our case as sampling of fossil terminals did not seek to represent the earliest members of lineages (Püschel et al. 2020). We sought to identify the most precise stratigraphic duration for all terminals, when possible down to the biozone, although period or age levels were used when occurrences were insufficiently constrained. Tip dates were enforced using uniform priors between the first and last appearances of 103 fossil terminals. Node dates were only used for monophyletic clades established as such by previous studies, as well as by our uncalibrated BI and MP analyses, and were intentionally avoided for nodes relating to the position of “cassiduloids”, Clypeasteroida, Scutelloida, Echinoneoida and Echinothurioida. Hard minimum and soft maximum bounds were enforced using offset exponential distributions. Mean values for the exponential distributions were set to leave 5% prior probability of the nodes being older than the maxima. Paleontological and stratigraphic justification of all tip and node dates employed can be found in SI File 3.

A uniform prior was used for the distribution of branch lengths, as more complex approaches such as the fossilized birth-death prior (Heath et al. 2014) failed to converge when performing total-evidence inference. An independent gamma rates model was employed using the default prior on the variance of the distribution, and a wide prior on the clock rate (normal distribution, mean = 0.001 and standard deviation = 0.01). Four independent runs of four Metropolis-coupled Markov-Chain Monte Carlo chains were continued for 100 million generations, with sampling of every ten thousand. The initial 25% were discarded as burn-in, and convergence and stationarity were confirmed by examining traces and posterior distributions using Tracer v. 1.7 (Rambaut et al. 2018) and *rwty* (Warren et al. 2017).

### Molecular Analyses

#### Matrix construction

Publicly available genomic and transcriptomic datasets were downloaded from either NCBI or EchinoBase (Kudtarkar and Cameron 2017). Taxonomic sampling corresponds to that of Mongiardino Koch et al. (2018), with the addition of four spatangoid (heart urchins; Romiguier et al. 2014) and two camarodont transcriptomes (Gaitán-Espitia et al. 2016; Clark et al. 2019). Given the presumed contamination between the transcriptomes of *Arbacia punctulata* and *Eucidaris tribuloides* detected by Mongiardino Koch et al. (2018), these datasets were replaced by the transcriptomes of *Arbacia lixula* (Pérez-Portela et al. 2016) and a different transcriptomic dataset from the same cidaroid species (Reich et al. 2015). Three holothuroid transcriptomes, representing highly divergent lineages (Miller et al. 2017), were used as outgroups. Further details, including SRA accession numbers, can be found in Table S2 (SI File 2).

**Table 2:** Macroevolutionary modeling of echinoid body size. Whenever multiple options of a given model were explored, only the best one was incorporated in the process of model comparison, and can be identified by the details shown in brackets (see Materials and methods and Fig. S17 of SI File 4). Asterisks identify non-uniform models. Models are ordered in decreasing order of fit (Diff. = difference in AICc between a given model and the best-fit one). The OUMVAZ model attains an AICc weight of 0.9999.

| Model | AICc | Diff. |
| --- | --- | --- |
| *OUMVAZ (4 regimes) | 260.0 | 0.0 |
| *OUMVA (5 regimes) | 277.9 | 17.9 |
| *OUM (7 regimes) | 289.0 | 29.0 |
| *BMS | 359.4 | 99.4 |
| BBMV (2 parameters) | 371.2 | 111.2 |
| Pulsed (NIG) | 371.6 | 111.6 |
| BBM | 372.4 | 112.4 |
| OU | 375.3 | 115.3 |
| FPK | 376.7 | 116.7 |
| Trend | 381.2 | 121.2 |
| BM | 383.0 | 123.0 |
| EB | 385.1 | 125.1 |
| White noise | 422.1 | 162.1 |

The majority of datasets employed (32 out of 37) are pair-end Illumina transcriptomes, and these were all assembled *de novo*. Raw reads were trimmed or excluded based on quality scores using Trimmomatic v. 0.36 (Bolger et al. 2014) before being further sanitized and assembled with the Agalma 2.0 phylogenomic workflow (Dunn et al. 2013) and Trinity 2.5.1 (Grabherr et al. 2011). The assembled transcriptomes were further processed with alien_index v. 3.0 (Ryan 2014) to remove transcripts with a likely non-metazoan origin, and CroCo v. 1.1 (Simion et al. 2018) to remove cross-contaminants product of multiplexed sequencing. These approaches removed on average 0.28% and 0.99% of transcripts per transcriptome, respectively (Table S2 of SI File 2 and Fig. S1 of SI File 4).

The sanitized transcriptomes were imported back into Agalma for orthology inference, alignment and supermatrix construction following the methods described in Dunn et al. (2013) and Guang et al. (2017). Four assembled single-end Illumina transcriptomes (also previously run through alien_index) and one genomic protein model were also incorporated at this stage (see Table S2 of SI File 2). The resulting supermatrix was reduced to a 70% occupancy value, composed of 2,356 loci (613,269 amino acids). This dataset was further curated from primary sequence errors using HmmCleaner (Di Franco et al. 2019), which relies on profile hidden Markov models to identify sequence segments that display a poor fit to their multiple sequence alignment (MSA). Given that regions with ambiguous alignment had already been removed by Agalma using GBlocks (Talavera and Castresana 2007), segment filtering was performed using the high-specificity parameter setting. Finally, gene trees were inferred as explained below, and scrutinized using TreeShrink v. 1.3.1 (Mai and Mirarab 2018) in order to filter out outlier sequences. This method detects sequences with unexpectedly long branches given species-specific distributions of proportional reduction in gene tree diameter after exclusion. Given the thorough sanitation already performed, we used a reduced tolerance for false positives (-q 0.01), and capped exclusion at 3 terminals per gene, each of which had to impact gene tree diameter by at least 25% (-k 3 -b 25). The successive application of these two filtering procedures increased the percentage of missing data from 40.81% to 41.33% (final occupancy of 69.7%). This final phylogenomic dataset is available as SI File 5.

#### Phylogenetic inference

We employed diverse approaches to phylogenetic inference, including coalescent and concatenation methods, based on both site-homogenous and site-heterogenous models. We first inferred gene trees using the parallel tool ParGenes v. 1.0.1 (Morel et al. 2018), which automated model selection using ModelTest-NG (Darriba et al. 2020) and phylogenetic inference using RAxML-NG (Kozlov et al. 2019) for each MSA. Support values for each gene tree were estimated using 100 replicates of non-parametric bootstrapping (BS). As explained above, these gene trees were then used to remove outlier sequences with TreeShrink. The set of gene trees was then updated, repeating model selection and tree inference for the 207 loci for which outliers were removed. Coalescent-based species tree inference was then performed using ASTRAL-III (Zhang et al. 2018), estimating support using local posterior probabilities (Sayyari and Mirarab 2016).

We also analyzed the supermatrix using two maximum likelihood (ML) concatenation approaches implemented in IQ-TREE 1.6.3 (Nguyen et al. 2014). First, we performed inference under the best-fitting partitioning scheme, found using the fast-relaxed clustering algorithm among the top 10% of schemes (Chernomor et al. 2016; Kalyaanamoorthy et al. 2017; Lanfear et al. 2017). Support was evaluated using 1,000 replicates of ultrafast bootstrap (Hoang et al. 2017). We then used the topology obtained to implement the PMSF method (Wang et al. 2017), a site heterogenous mixture model that accounts for heterogeneity in the process of amino acid substitution between sites. Support values were derived from 100 replicates of BS.

### Combined Analyses

#### Dataset merging

A TED analysis was performed by combining the morphological, molecular and stratigraphic datasets. Given the computational burden of this approach, the molecular dataset had to be drastically reduced. Even though subsampling of loci has become standard in phylogenomic research, this is generally performed using a single gene property such as the rate of molecular evolution (e.g., Sharma et al. 2014), a given proxy for phylogenetic signal (e.g., Salichos and Rokas 2013) or a potential source of systematic bias (e.g., Nesnidal et al. 2010). When multiple properties have been considered, these have usually been optimized sequentially (e.g., Whelan et al. 2015; Smith et al. 2018). Here, we quantified seven gene properties routinely employed for phylogenomic subsampling, including: 1) Level of saturation, estimated as one minus the slope of the regression of patristic distances versus *p*-distances (Nosenko et al. 2013); 2) Average pair-wise patristic distance; 3) Compositional heterogeneity, measured by the relative composition frequency variability (RCFV; Nesnidal et al. 2010); 4) Proportion of invariant sites; 5) Average BS support; 6) Gene tree error, using the Robinson-Foulds distance (Robinson and Foulds 1981) to the species tree; and 7) Clock-likeness, estimated as the variance of root-to-tip distances. With the exception of RCFV, which was estimated using BaCoCa v. 1.103 (Kück and Struck 2014), all other indices were calculated in the R environment (R Core Team 2019) using data files and code available in SI File 6. This relied on functions from packages *adephylo* (Jombart and Dray 2010), *ape* (Paradis and Schliep 2018), *phangorn* (Schliep 2011) and *phytools* (Revell 2012).

We explored the pattern of correlation among these metrics using Spearman rank correlations and principal component analysis (PCA). To better understand what determines the structure of correlation in this dataset, we also estimated the rate of evolution of each locus using both tree-based and tree-independent methods. The evolutionary rate of each site was estimated using the empirical Bayes method implemented in Rate4Site (Mayrose et al. 2004). This was achieved by first time-calibrating the partitioned ML topology using penalized likelihood (Sanderson 2002), as implemented in the function ‘chronos’ of R package *ape*. We employed the correlated model of substitution rate variation among branches and six of the node calibrations defined in SI File 3. The resulting chronogram can be found as Figure S2 (SI File 4). As an alternative proxy we used the tree-independent method TIGER (Cummins and McInerney 2011), which relies exclusively on congruence between characters. This approach is based on the observation that partitions defined by fast-evolving characters will likely show low repeatability and high disagreement, being mostly determined by noise rather than phylogenetic signal, whereas slow-evolving positions are expected to partition terminals into more similar subgroups. For both approaches, we excluded outgroups and eliminated estimates for positions with 5 or less non-missing entries. We then averaged estimates across sites to obtain a single value that describes the evolutionary rate of each locus.

Subsampling of loci was then performed retaining those that exhibited the highest score along PC 2 of the gene properties dataset. Multivariate statistical analyses revealed that this dimension provides a means of selecting subsets of genes with increased phylogenetic signal and clock-like behavior, reduced evidence of systematic biases and intermediate rates of evolution (see Results). The top scoring 300 genes were selected, and the level of occupancy of all terminals in this reduced matrix was calculated (Fig. S3 of SI File 4). Given that our molecular dataset includes 34 echinoid terminals, which represent only 21 lineages in the morphological partition, we sampled one terminal per main lineage in order to assemble the combined dataset. When occupancy was markedly uneven between different representatives of the same clade, we chose the one with the highest occupancy, otherwise we selected the one considered to be most closely related to the one sampled in the morphological dataset (Fig. S3 of SI File 4). Given that the morphological dataset was designed to include only characters variable between families/subfamilies, the generation of composite terminals is not expected to be problematic. We then retained 50 loci with the least amount of missing data across the selected taxa (21 echinoids and 3 holothuroid outgroups), resulting in a final dataset of 13,933 amino acid positions and an occupancy of 87.2%.

#### Phylogenetic inference

The 50 loci selected were combined with the morphological dataset to build the final combined matrix, available as SI File 7. Terminals bear the name of the least inclusive clade that contains the species sampled in the morphological, molecular, and stratigraphic datasets (see Table S1 of SI File 2). A TED analysis of this combined dataset was performed in MrBayes. The best-fit partitioning scheme for the molecular data was determined using the greedy algorithm in PartitionFinder2 (Lanfear et al. 2017). In order to enforce correct rooting with molecular or morphological outgroups, the nodes corresponding to Echinozoa (Echinoidea + Holothuroidea), all sampled Echinoidea, and all echinoids except for *Archaeocidaris*, were constrained and given node age priors. Further details on time calibration can be found in SI File 3. The analysis was parameterized as for the morphological analyses and run for 70 million generations, and convergence was evaluated as previously described.

The phylogenetic position of a few terminals was explored using RoguePlots (Klopfstein and Spasojevic 2019), and ancestral state reconstruction of several morphological features was performed using 1,000 replicates of stochastic character mapping under equal rates (Bollback 2006).

### Macroevolution of Echinoid Body Size

#### Body size estimation

We gathered a comprehensive dataset of length, width and height of echinoid tests from both the primary literature and specimens in museum collections. In total we obtained information from 5,317 specimens, representing an average of 33.7 observations per species. Further details on this dataset can be found in SI File 8. We transformed these measurements into an estimate of body size using the formula for the volume of a regular ellipsoid (*V* = ^π^⁄_6_ *lw*ℎ). This biovolume approximation has been widely used to estimate body size in similarly shaped organisms (e.g., Finnegan and Droser 2008; Church et al. 2019), including echinoids at the intra-specific level (Uthicke et al. 2016). Biovolume was then averaged by species and log_10_ transformed (the dataset is available as SI File 9).

#### Macroevolutionary modeling

We used the maximum clade credibility (mcc) tree and 20 randomly selected topologies from the posterior distribution of the TED analysis to explore the macroevolution of echinoid body size. The three holothuroid outgroups and six echinoids for which no reliable body size data could be gathered were pruned (see SI File 8). We analyzed the phylogenetic signal of the trait using Pagel’s λ (Pagel 1999) and the temporal dynamics of disparity using disparity-through-time (DTT) plots (Harmon et al. 2003). The empirical DTT trajectory was compared with 10,000 replicates of character evolution under Brownian motion (BM), and deviations were evaluated using the rank envelope test (Murrell 2018).

To further explore the process underlying body size evolution, we fit a suite of models of continuous trait macroevolution. We included a BM model, the most common null model of evolution in comparative studies, which represents a process of random walk with constant variance (Felsenstein 1985; Freckleton et al. 2002). The dynamics of this model are governed by a single parameter (σ^2^) that determines the rate at which independent lineages diffuse away from an ancestral condition. BM is a special case of the more general Ornstein-Uhlenbeck (OU) model, which includes an additional tendency (α) to evolve towards an optimum value (θ). This added deterministic component emulates a process of constrained morphological evolution within adaptive zones (Hansen 1997; Butler and King 2004; Beaulieu et al. 2012). We also incorporated several modifications of the BM process that capture other conceivable ways in which traits might change through time, including a directionality to the process of diffusion (trend model; Hunt and Carrano 2010), and rates that vary exponentially with time (EB model; Blomberg et al. 2003; Harmon et al. 2010). All of these models account for phylogenetic relatedness, assuming a covariance structure between species that is proportional to their shared evolutionary history. This assumption was further relaxed using a white noise model.

For many years, these relatively simple and uniform (i.e., using a single process for the entire tree) models were the only ones available for exploring the dynamics of morphological change between species. However, the repertoire of models has been recently expanded, and we focused specifically on including and comparing the behavior of this expanded toolkit. We incorporated the bounded Brownian motion model (BBM; Boucher and Démery 2016), which restricts the variance that traits can attain by enforcing reflective boundaries; as well as the family of Lévy process models (Landis and Schraiber 2017) specifically designed to capture the pattern of stasis punctuated by bursts of change that abounds in the fossil record. Furthermore, we explored several alternative ways of characterizing a macroevolutionary adaptive landscape (Simpson 1944; Hansen 2012). First, we employed the Fokker-Planck-Kolmogorov (FPK) model (Boucher et al. 2017), which can approximate adaptive landscapes with more complex shapes than those assumed by OU models, thereby accommodating scenarios of directional and disruptive selection. This is achieved by inferring an evolutionary potential, which biases the underlying random walk of all lineages towards different regions of morphospace. This model, as well as the previously described BBM can be thought of as special cases of a general model (BBMV; Boucher 2019) that combines the attractive force of the evolutionary potential with maximum and minimum trait boundaries.

As an alternative, we also explored the fit of a series of multi-OU (OUM) models, which constitute convenient representations of evolution towards adaptive peaks (Mahler et al. 2013). In these models, in contrast to BBMV, different peaks can only be discovered by certain lineages (which are inferred to be under the same regime), and do not affect the evolutionary dynamics of other regions of the tree. For the same reason, OUM models differ from all other models considered in being non-uniform, i.e. distinct sets of parameters are applied to different branches of the phylogeny. Although the number and location of regime shifts in OUM models are often defined *a priori*, some implementations can infer the location of regime shifts (as well as the parameters that characterize regimes) without any previous specification. One such method, SURFACE (Ingram and Mahler 2013), is employed here. SURFACE operates by iteratively locating shifts in the value of trait optima (θ) along branches of the tree (forward phase), selecting at each step the model that minimizes the Akaike’s information criterion for finite sample sizes (AICc). Once the AICc cannot be improved, optimization is attempted by merging independent regimes, which are thus considered to have been convergently encountered by different lineages (backwards phase).

Regimes identified by SURFACE vary exclusively in their value of θ; both α and σ^2^ are kept constant across the phylogeny. To explore more complex models, we employed OUwie (Beaulieu et al. 2012) which requires *a priori* identification of selective regimes but can estimate independent sets of parameters for each of them. The regimes identified by SURFACE were therefore used as input in OUwie, allowing the estimation of all three parameters for each regime (model OUMVA), and a further model for which the ancestral state at the root is not constrained to be the optimal trait value for the ancestral regime, but is free to vary as an additional parameter (model OUMVAZ). As these models can often be difficult to fit, the reliability of parameter estimates was evaluated by checking that all eigenvalues of the Hessian matrix were positive (Beaulieu et al., 2012). The temporal dynamics of OUM models (and their extensions) were explored using phylogenetic half-lives (*t*_1/2_), which measure the amount of time needed for a lineage to move half-way from its current trait value to a new optimum (Hansen 1997). Finally, to test if the heterogeneity captured by SURFACE as different regimes can be simply explained by differences in evolutionary rate (rather than jumps in adaptive landscape), a variable-rate BM (BMS) model was also fit, inferring different σ^2^ for each regime.

Model fitting relied on packages *geiger* (Harmon et al. 2008), *BBMV* (Boucher 2019), *pulsR* (Landis and Schraiber 2017), *SURFACE* (Ingram and Mahler 2013) and *OUwie* (Beaulieu and O’Meara 2019). We compared the fit of all of the models described above through the use of AICc weights. For BBMV, the shape of the evolutionary potential was estimated using one, two and three free parameters (Boucher 2019), and only the best option was included in the final model comparison. Similarly, for the pulsed models, the fit of the JN, NIG, BMJN and BMNIG models was first compared and only the best was used for model selection (Landis and Schraiber 2017). Measurement errors were incorporated in the estimation process, calculated as squared standard errors of the mean values (see SI File 8 and Fig. S4 of SI File 4 for further details). Data files and R code necessary to fit and compare these models are available as SI File 10.

## Results

### Molecular Phylogeny

All methods of phylogenetic inference recovered the same topology, with all nodes attaining maximum support values (Fig. 1). This topology is identical to that of Mongiardino Koch et al. (2018), confirming their higher-level phylogenetic structure with a better-curated matrix including more than twice the number of loci. Euechinoids are subdivided into Aulodonta—including echinothurioids, diadematoids and pedinoids—and Carinacea (*sensu* Kroh 2020). The monophyly of Clypeasteroida + Scutelloida is again not supported, with the only sampled “cassiduloid” confidently resolving as the sister group to scutelloids. The improved sampling and data curation also provide a robust resolution of the topology within Echinacea, the clade containing stomopneustoids, arbacioids and camarodonts. Early molecular phylogenies supported a close relationship between stomopneustoids and arbacioids (Smith et al. 2006), whereas morphological data resolved arbacioids and camarodonts as sister clades (Kroh and Smith 2010), a result utilized in the latest classification (Kroh 2020). Previous phylogenomic work tentatively rejected both topologies, supporting instead a clade containing Stomopneustoida + Camarodonta (Mongiardino Koch et al. 2018), a result that is here confirmed. The topologies within spatangoids, camarodonts and scutelloids agree with previous molecular and morphological studies (Mooi 1990b; Stockley et al. 2005; Smith et al. 2006; Kroh and Smith 2010; Thompson et al. 2017;Mongiardino Koch et al. 2018; Bronstein and Kroh 2019).

**Figure 1:**
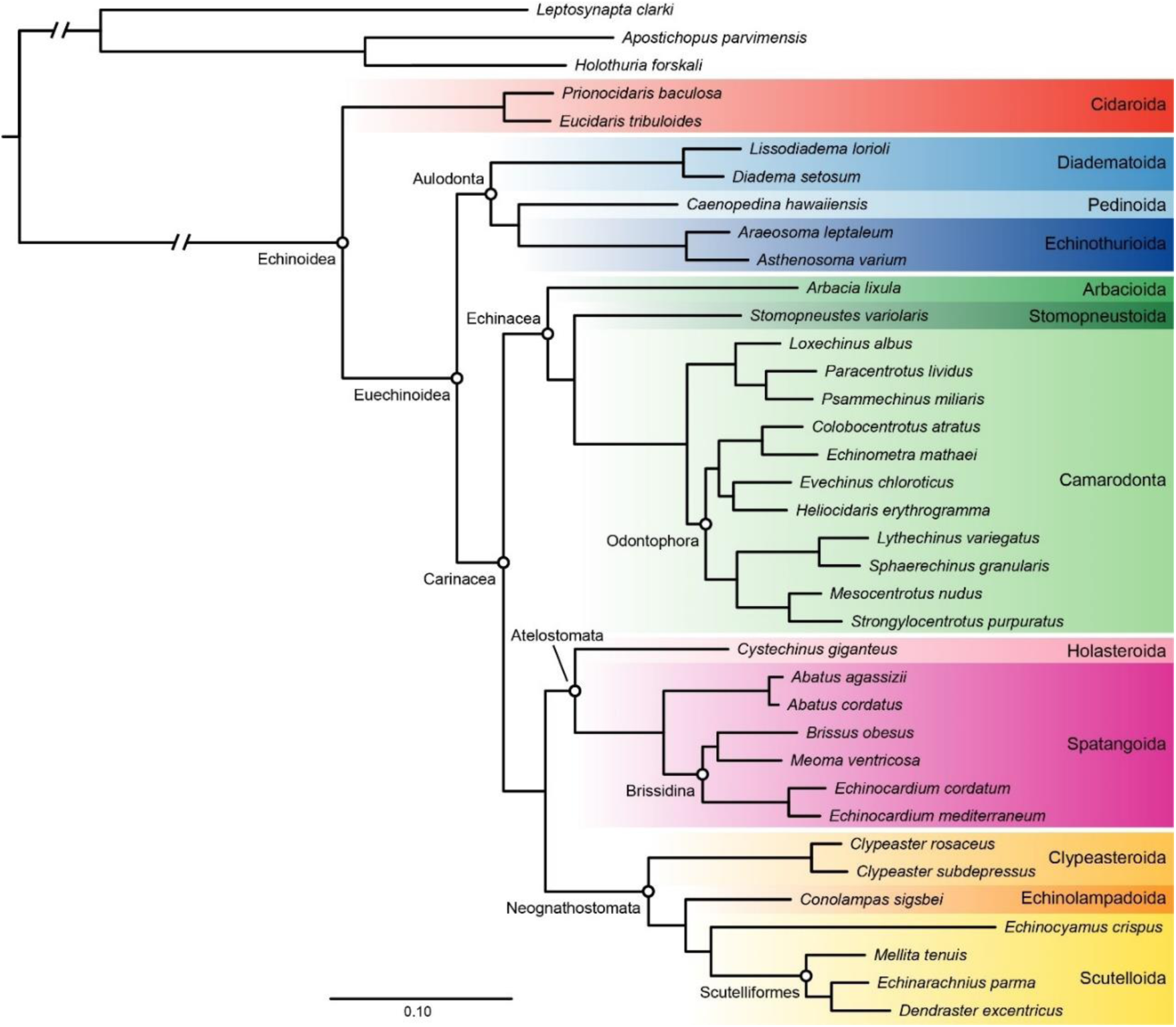
Phylogenomic tree obtained through all three methods of inference (ASTRAL, site-homogenous and site-heterogenous ML). All nodes attained maximum support across inference approaches. Branch lengths are those of the site-homogenous ML analysis.

### Morphological Phylogeny

As expected, both MP analyses (equal and implied weights, Figs. S5-6 of SI File 4) returned broadly similar topologies to those previously found for this dataset (Kroh and Smith 2010; Hopkins and Smith 2015). In these, echinothurioids are the sister clade to all other euechinoids, and the other four aulodont lineages (aspidodiadematoids, diadematoids, pedinoids and micropygoids) are spread across the euechinoid backbone along a succession of poorly supported nodes. The topology within Neognathostomata is highly uncertain and differs markedly depending on method of inference, especially regarding the position of stem neognathostomates, stem clypeasteroids and “cassiduloids”. Echinoneoids are strongly supported as the sister clade of all other crown group irregular echinoids.

Bayesian approaches, on the other hand, produced a number of novel and strongly supported topological rearrangements, the most significant of which relates to the position of echinothurioids and the other aulodonts. Neither uncalibrated (Fig. S7 of Si File 4) nor calibrated BI (Fig. 2) analyses support a sister-group relationship between echinothurioids and the remaining euechinoids. Instead, uncalibrated BI strongly supports the monophyly of a clade consisting of echinothurioids, diadematoids and micropygoids, plus the extinct pelanechinids. The implementation of a morphological clock further results in the addition of aspidodiadematoids and pedinoids to this clade (Fig. 2), recovering a monophyletic Aulodonta in full agreement with molecular data.

**Figure 2:**
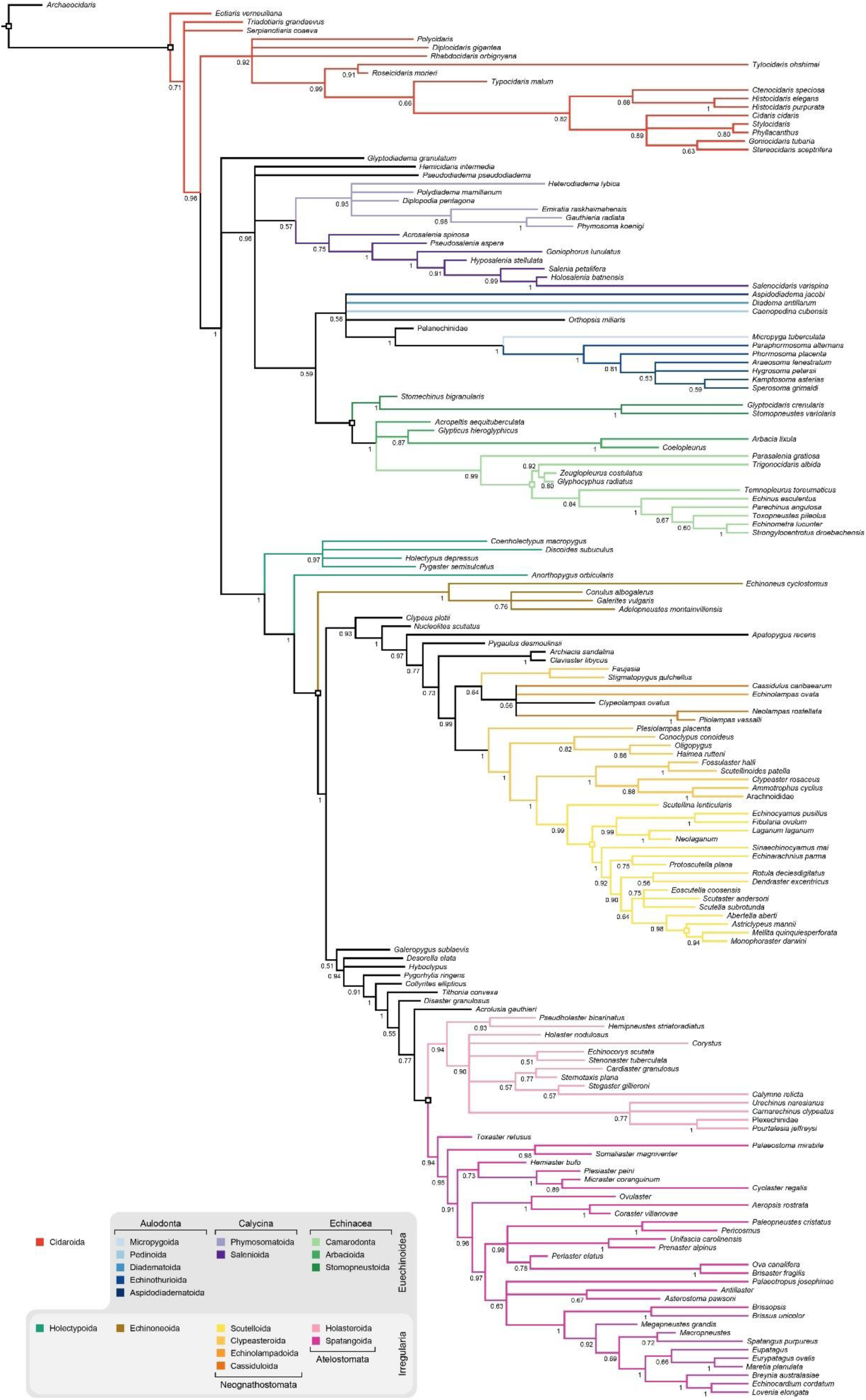
Majority-rule consensus tree of the calibrated BI analysis of morphology. Branches are colored according to the classification of terminals in the World Echinoidea Database (Kroh and Mooi 2019), updated according to Kroh (2020). Numbers on branches represent posterior probabilities. Squares denote nodes that were constrained and assigned age priors.

BI also provided some stability to the relationships among neognathostomates. Both calibrated and uncalibrated approaches show strong support for a clade containing all lineages currently classified as clypeasteroids, scutelloids, cassiduloids and echinolampadoids, as well as the extinct clypeolampadids (Figs. 2 and S7 of SI File 4). However, the monophyly of Clypeasteroida as currently defined is not recovered. With the addition of stratigraphic information this clade splits into two groups (Fig. 2). The first contains faujasiids, clypeolampadids, echinolampadoids and cassiduloids, all of which were at some point included within Cassiduloida, but many of them were removed by Kroh and Smith (2010) in an attempt to find natural subdivisions of the “cassiduloids”. Our results suggest that all of these lineages are closely related to each other, as suggested by Souto et al. (2019). The other subdivision contains a clade of Scutelloida + crown group Clypeasteroida which is subtended by plesiolampadids, conoclypids and oligopygids (Fig. 2).

Finally, both BI approaches resolve *Eotiaris*, *Serpianotiaris* and *Triadotiaris*, traditionally considered as the earliest members of crown group Echinoidea, along the stem of the clade. Their novel position renders Cidaroidea (one of the two subclasses of crown group echinoids, including Cidaroida; Kroh 2020) paraphyletic with respect to Euechinoidea. This topology has strong consequences for the timing of origin of crown echinoids, the dynamics of stem and crown group echinoids across the P-T mass extinction, and patterns of morphological evolution among the earliest members of the clade (see Discussion).

### Subsampling of Loci

The exploration of patterns of correlation between gene properties revealed several interesting patterns. As expected, all potential sources of systematic bias (level of saturation, compositional heterogeneity, average patristic distance and root-to-tip variance) were positively correlated with each other (Fig. 3a). However, these were also found to be positively correlated with the average BS of gene trees, and negatively correlated with gene tree error. Subsampling based on these commonly-used proxies for phylogenetic signal will enrich the dataset in problematic loci, whereas the opposite strategy, reducing the degree of systematic biases present in the dataset, will enrich it in uninformative loci (Fig. 3b, left panel). Especially strong are the relationships between phylogenetic signal and average patristic distance (ρ = 0.58 and 0.44 against average BS and gene tree error, respectively; both *P* < 10^-16^) and root-to-tip variance (ρ = 0.30 for average support; *P* < 10^-16^), properties that can induce topological artifacts and bias divergence times.

**Figure 3:**
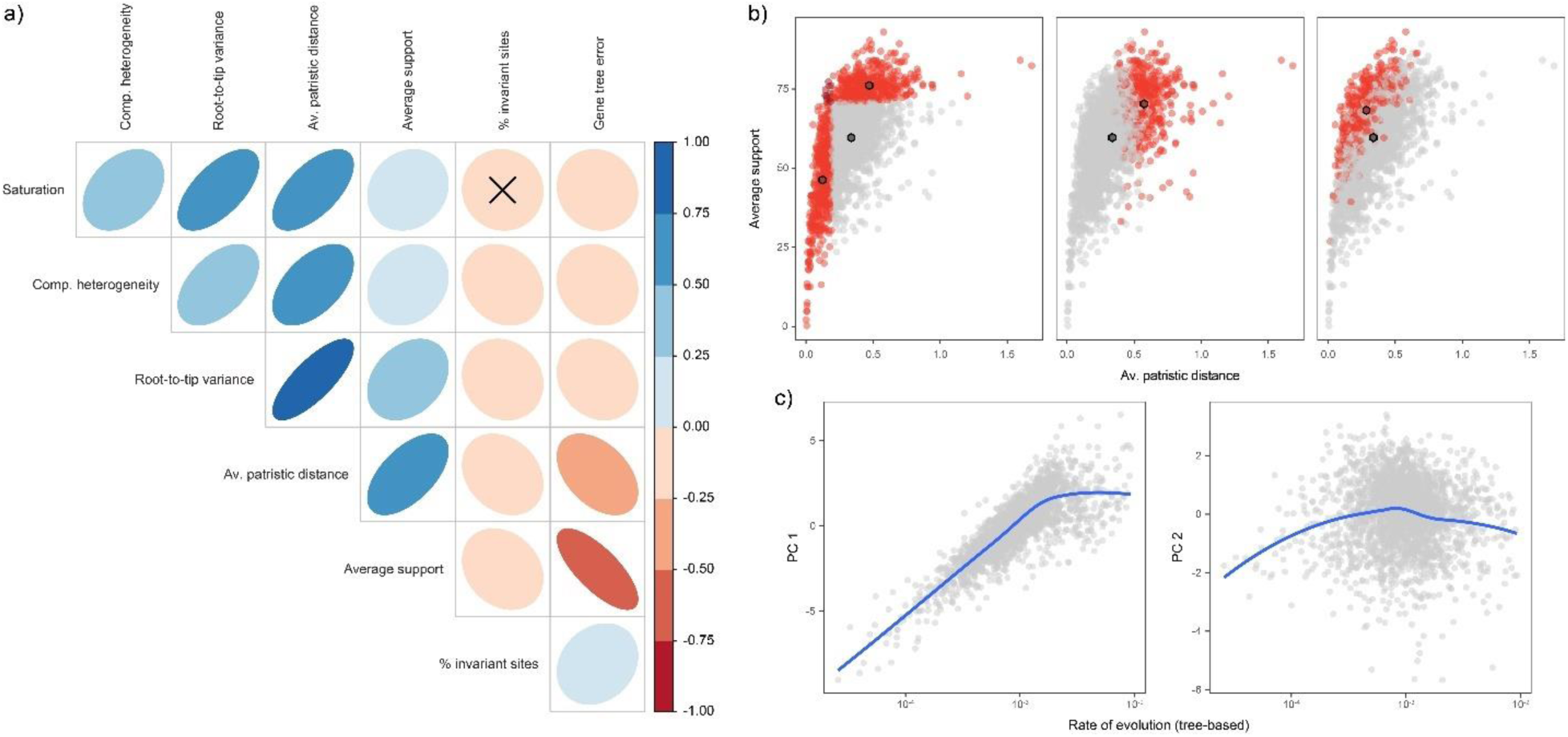
Exploration of different approaches to loci subsampling. **a)** Correlation structure between seven gene properties representing either sources of systematic bias or proxies for phylogenetic signal. **b)** Alternative methods to subsample molecular datasets. Each point represents a locus, with those in red represent the top scoring 20% under a given subsampling scheme. Hexagons show the centroid of the distributions. Left: Subsampling genes with high phylogenetic signal also increases sources of bias in the dataset, while subsampling those unaffected by bias decreases average signal. These two approaches also produce almost entirely non-overlapping subsets of loci. Center: Subsampling based on PC 1 selects loci with high phylogenetic signal but also enriched in potential sources of bias. Right: Subsampling based on PC 2 selects loci with high phylogenetic signal while decreasing sources of error. c) PC 1 strongly correlates with rate of evolution (left), while PC 2 favors loci with intermediate rates. Blue lines are loess regressions.

As an alternative, loci subsampling can be performed while directly accounting for this structure of correlation. To do this, we explored a novel approach relying on PCA. Interestingly, the first PC axis seems to select loci with both high levels of phylogenetic signal and systematic biases, while the second axis maximizes signal while decreasing possible sources of bias (Table 1). We present a visual representation of how subsampling along these two dimensions affects loci selection in Figure 3b (center and right panels). Estimation of the average evolutionary rate of loci reveals the underlying factor generating this pattern. The first PC axis largely orders loci according to their evolutionary rate (Fig. 3c, left panel; linear regression: R^2^ = 0.65, *P* < 10^-16^), selecting loci that are highly saturated but generate gene trees with above-average levels of support. The second PC axis, in contrast, seems to select loci with intermediate rates of evolution (Fig. 3c, right panel). These rates are enough to produce highly supported and accurate gene trees, but not high enough to accumulate noise (Table 1). This result holds true if a tree-independent method is used to estimate evolutionary rate (Fig. S8 of SI File 4). Even though tree-independent methods have been criticized (Simmons and Gatesy 2016), TIGER produces rate estimates that have a strong log-linear relationship with those of a tree-based approach (Fig. S9 of SI File 4; see Mongiardino Koch and Gauthier 2018).

In order to further explore the effect of subsampling loci along PC 2, the properties of the 300 top-scoring loci were compared to those of 10,000 randomly selected subsets of equal size (Fig. 4). Our subsampling approach simultaneously increases phylogenetic signal, decreases potential sources of systematic bias, and increases the clock-like behavior of the data. Furthermore, it selects loci that are longer and exhibit reduced evolutionary rates (Fig. 4), even though these properties were not included in the PCA. Not only were the average values of all of these properties significantly different from those obtained through random subsampling (all *P* < 10^-4^), the variances of all properties were also significantly smaller than expected by chance (all *P* < 0.025). Thus, our method significantly decreases heterogeneity in the dataset, likely reducing model misspecification.

**Figure 4:**
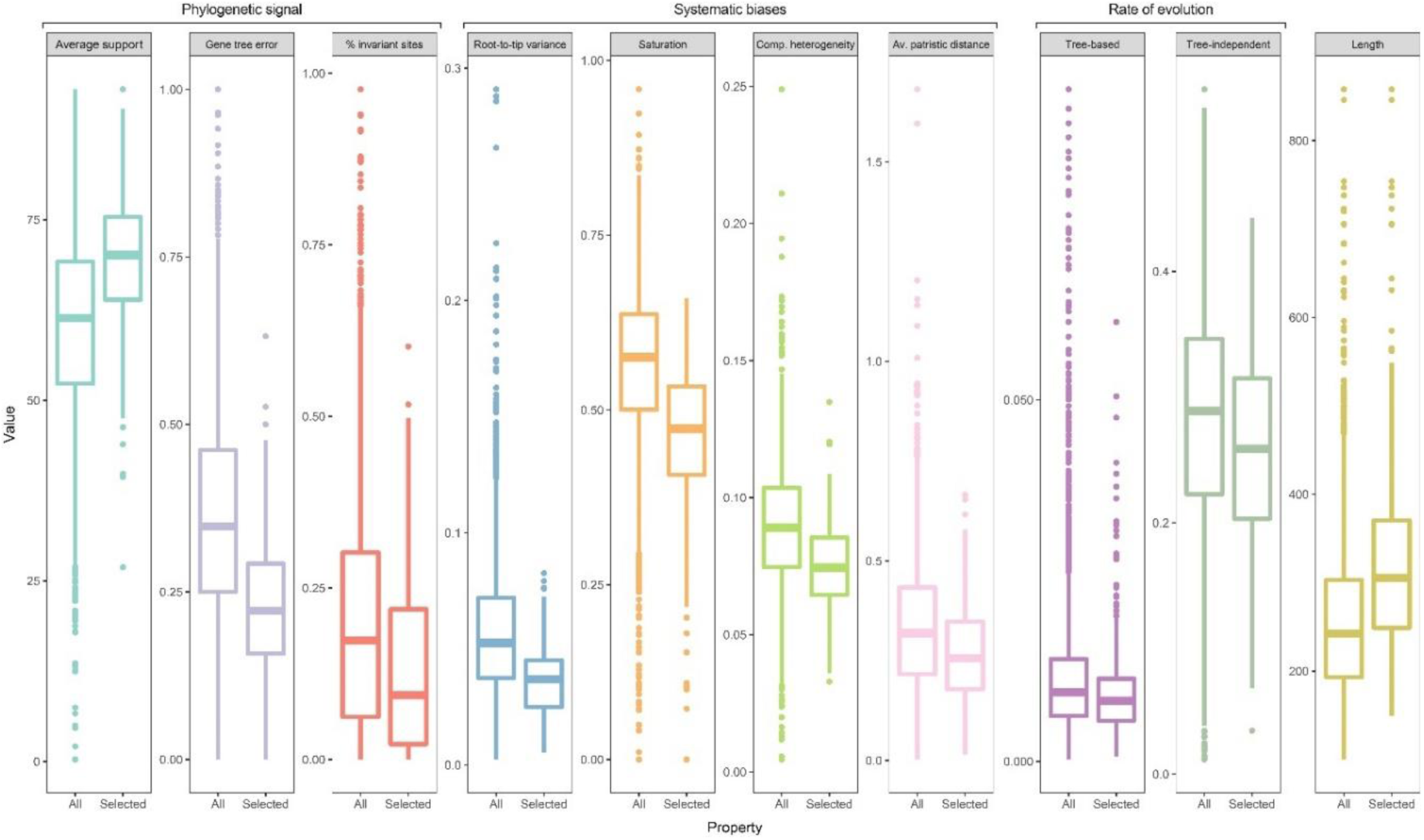
Effect of subsampling loci along PC 2 of the gene properties dataset. For each variable, the boxplot on the left represents the distribution across all 2,356 loci, while the one on the right shows that of the top-scoring 300 loci along PC 2. Subsampling based on PC 2 produces means that significantly differ from those expected under random subsampling (as well as significantly lower variances) for all variables.

Partitioned ML inference on the subsampled 300-gene dataset produced a topology identical to that of Figure 1 (Fig. S10 of SI File 4), with all nodes attaining more than 95% BS support, further demonstrating the utility of the approach. After further reducing the dataset to 21 echinoids and three holothuroids (as explained above, see Fig. S3 of SI File 4), and retaining the 50 loci with the lowest proportions of missing data, the molecular matrix was combined with the stratigraphic and morphological datasets to perform a TED analysis. Neither of these additional steps of matrix reduction modified the inferred topology (Figs. S11-S12 of SI File 4).

### Total-Evidence Phylogeny

The majority-rule consensus tree (Fig. 5) is the first to rely on morphological, molecular and stratigraphic data to resolve the relationships among the main lineages of extant and extinct echinoids (see also the mcc tree in Fig. S13 of SI File 4 and the inferred divergence times of major clades in Table S3 of SI File 2). The resulting topology strongly supports the position of *Eotiaris*, *Serpianotiaris* and *Triadotiaris* (the earliest fossil clades assigned to the crown group) along the echinoid stem. Despite the different placement of these early lineages, the age of crown group Echinoidea is inferred to be 266.9 Ma (245.1-287.3 Ma 95% highest posterior density, HPD), similar to that of previous studies (Smith et al. 2006; Nowak et al. 2013; Thompson et al. 2017) and assigning a high probability to an origin prior to the P-T mass extinction. Even though morphological approaches yielded a diversity of arrangements for the main lineages of euechinoids (Figs. 2 and S5-S7 of SI File 4), the TED tree agrees with molecular data in splitting this clade into Aulodonta and Carinacea, the latter composed of a monophyletic clade of Echinacea + Calycina (also including Pseudodiadematidae and Hemicidaridae) which forms the sister group to Irregularia. The aulodont-carinacean split also likely predated the end Permian mass extinction (257.8 Ma, 95% HPD: 237.9-283.3 Ma), implying multiple lineages of crown group echinoids survived this event.

**Figure 5:**
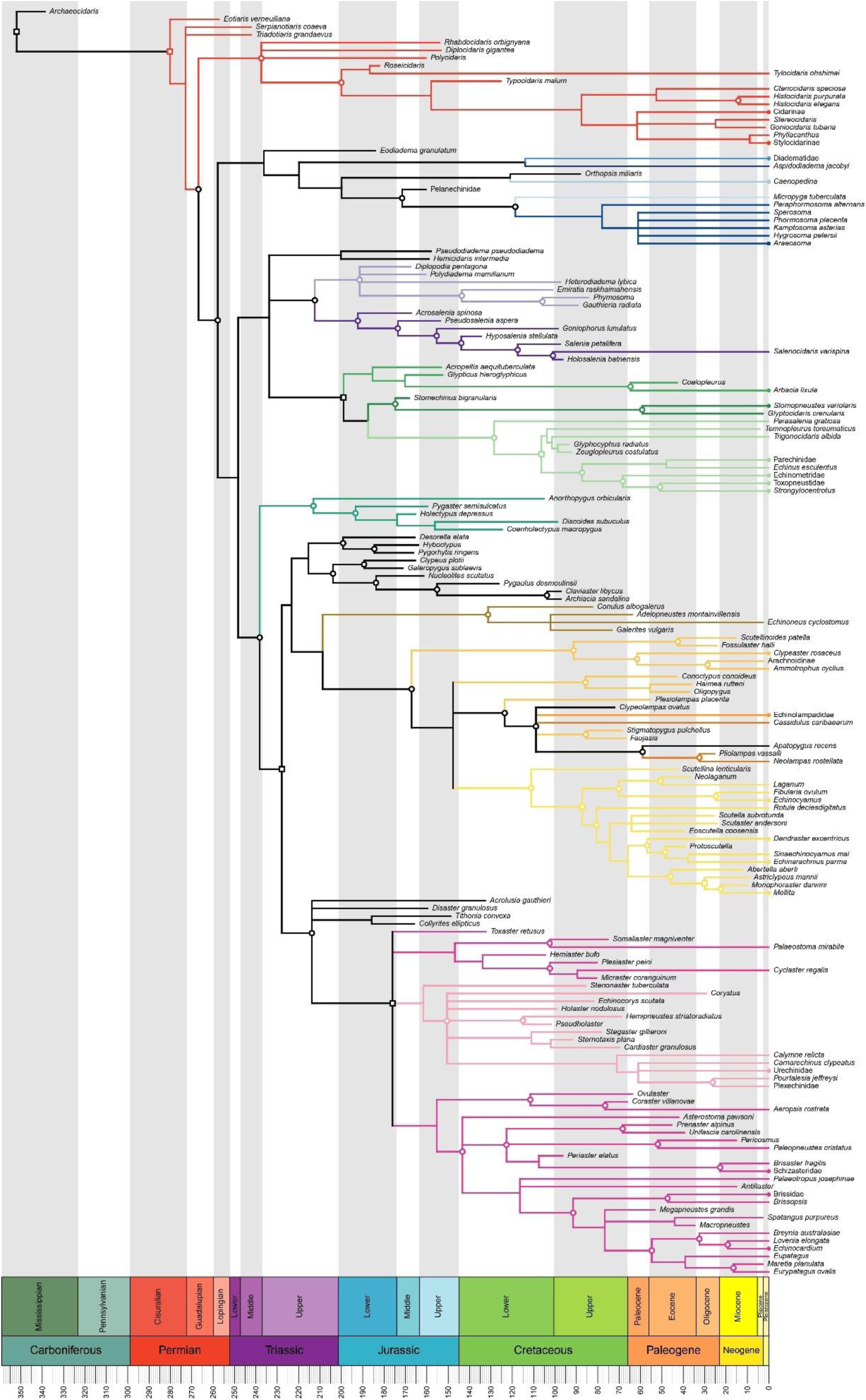
Majority-rule consensus tree of the total-evidence dated analysis. Clades are colored as in Fig. 2. Circles highlight nodes with posterior probability > 0.9, as well as tips with molecular data. Squares denote nodes that were constrained and assigned age priors. Holothuroid outgroups were pruned. The maximum clade credibility tree can be found in Fig. S13 of SI File 4; divergence times for higher-level clades are shown in Table S3 of SI File 3.

**Table 3:**
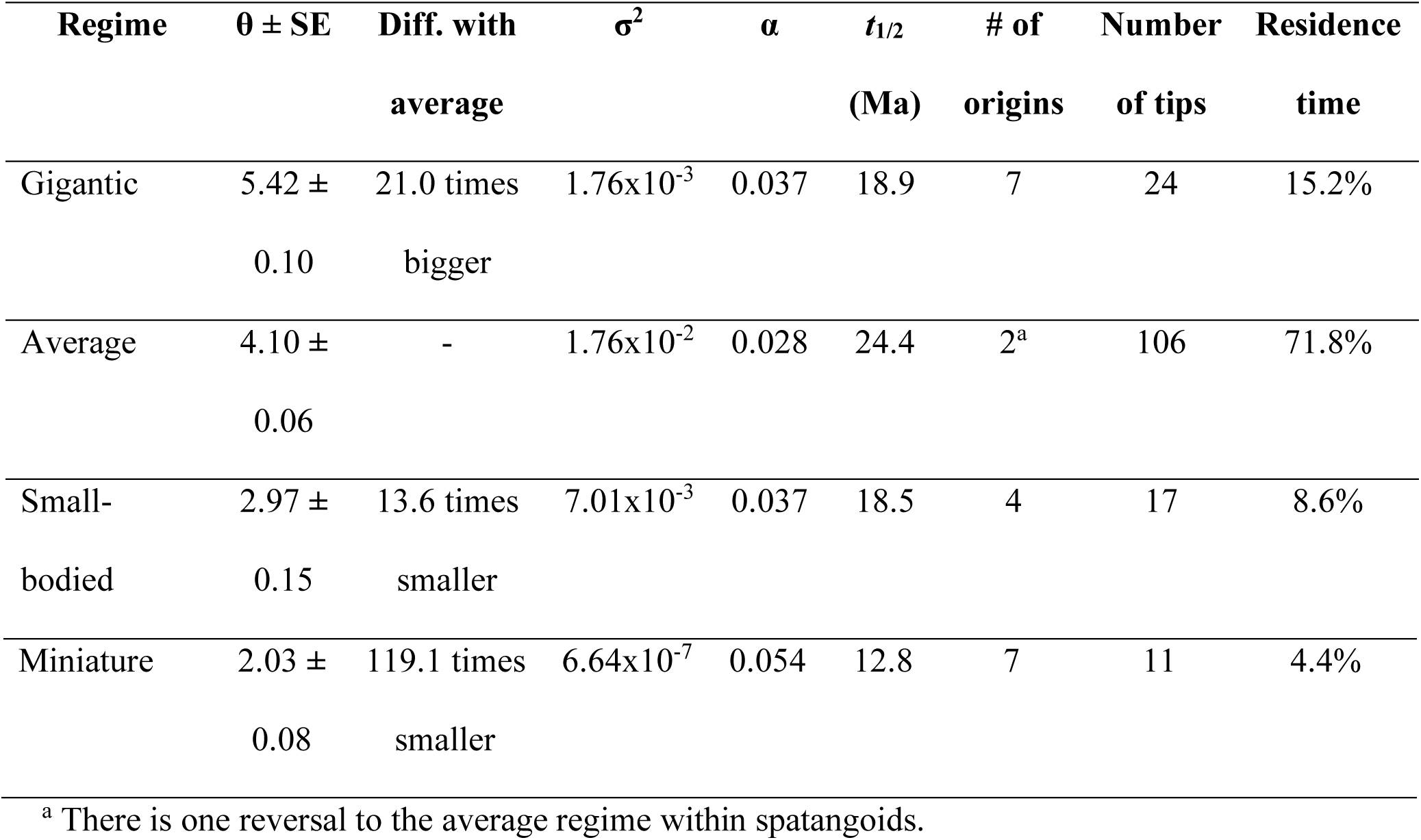
Parameters of the four-peak OUMVAZ model. “Diff. with average” is the difference in optimum body sizes for regimes against the ancestral (average) one. Residence time was calculated using stochastic character mapping under equal rates, and expressed as percentage of total tree length. SE = standard error. Z_0_ (state at the root) = 4.22 ± 0.87.

Much of the remaining tree agrees with the morphological and molecular phylogenies, reflecting relatively high levels of congruence between data sources (at least when morphology is analyzed under calibrated BI, see above). However, the topology within Neognathostomata, and the relationships between this clade and other members of Irregularia, undergoes a dramatic reorganization in our TED analysis. The incorporation of molecular data disrupts the monophyly of Scutelloida + crown group Clypeasteroida found in morphological analyses, suggesting that multiple lineages of “cassiduloids” and fossil neognathostomates are nested within this clade. These are organized into two strongly supported groups. The first includes the clade of faujasiids, clypeolampadids, echinolampadoids and cassiduloids already found in the calibrated BI morphological analysis, to which apatopygids and plesiolampadids are added. This clade is found to be closely related to the sand dollars (Fig. S13 of SI File 4), as supported by molecular data (Fig. 1). The inclusion of *Apatopygus recens* in this clade is unexpected, given that all morphological analyses resolve this species as distantly related to the other lineages of extant “cassiduloids” (Figs. 2 and S5-S7 of SI File 4). The second group is composed of conoclypids and oligopygids, a clade previously recognized by multiple authors (Durham and Melville 1957; Philip 1965; Wagner and Durham 1966). Its exact position is left unresolved, with almost equal probability of being the sister group to the clypeasteroids or to the other clade of “cassiduloids” previously mentioned (Fig. S14 of SI File 4). The divergence between Clypeasteroida and Scutelloida (representing the last common ancestor of crown group Neognathostomata in this topology) is estimated to have occurred 167.1 Ma (95% HPD: 145.0-190.7 Ma) during the Middle Jurassic, consistent with the deep fossil record of some of the lineages it includes.

The repositioning of the extant “cassiduloids” and stem clypeasteroids also strongly modifies the backbone phylogeny of Irregularia. Echinoneoids, traditionally considered to be the sister group to all other crown group irregular echinoids (Mortensen 1948a; Durham and Melville 1957; Smith 1981; Kroh and Smith 2010), resolve instead as the sister group of neognathostomates, in agreement with molecular data (Lin et al. 2019). This larger clade (Neognathostomata + Echinoneoida) is in turn found to be sister to a large monophyletic group of extinct irregular echinoids with previously unclear affinities. Although many of these families (Archiaciidae, Clypeidae, Nucleolitidae and Pygaulidae; Kroh and Mooi 2019) are currently classified as stem group neognathostomates, others have often been recovered along the stem (Kroh and Smith 2010) or even the crown (Barras 2007) of Atelostomata (as in our morphological analyses, Figs. 2 and S5-S7 of SI File 4). Our topology therefore suggests that atelostomates represent the sister clade to all other crown group irregular echinoids, which are here dated to 227.9 Ma (95% HPD: 210.4-242.5 Ma).

The stability provided by molecular data to the echinoid backbone thus provides a robust resolution of the position of many lineages for which only morphological data are available. Beyond the examples of “cassiduloids”, stem group neognathostomates and echinoneoids already mentioned, other unstable taxa also resolve in positions more consistent with their presumed affinities. *Anorthopygus orbicularis*, for example, is classified as a member of Holectypoida, but only resolves as such in the TED analysis, which alone recovers the monophyly of this order. Similarly, *Orthopsis miliaris*, noteworthy for displaying a mix of aulodont and echinacean attributes (Mortensen 1943; Durham and Melville 1957; Durham 1966a), resolves as an aulodont with a posterior probability (pp) of 0.79 (Fig. S15 of SI File 4). The most likely position for orthopsids is sharing a common ancestry with pedinoids, as previously suggested (Fell and Pawson 1966). The addition of molecular data also rearranges the topology within scutelloids, resulting in rotulids and taiwanasterids shifting away from the problematic positions recovered by Kroh and Smith (2010), and resolving instead in positions more concordant with previous treatments (Mortensen 1948b; Durham 1966b; Seilacher 1979; Mooi 1990c; Mooi 1990b; Ziegler et al. 2016).

Other aspects of our combined topology are less easy to reconcile with previous findings. Even though none of our analyses confirms the monophyly of the two main subdivisions of crown group cidaroids currently recognized (i.e., with Histocidaridae + Psychocidaridae sister to the rest; Kroh and Mooi 2019), both of our tip-dated analyses rearrange the clade is ways that are discordant with molecular and morphological studies (Kroh and Smith 2010; Brosseau et al. 2012). The relative stem-ward movement of psychocidarids (*Roseicidaris* and *Tylocidaris ohshimai*) and typocidarids in both tip-dated analyses might indicate that this approach favored topologies with better stratigraphic fit at the expense of morphological evidence (King and Beck 2019). The resolution within Aulodonta also disagrees with previous hypotheses by allying *Micropyga* with echinothurioids instead of diadematoids (Mortensen 1940; Durham and Melville 1957). Finally, our TED phylogeny lacks the level of resolution within Atelostomata found in the calibrated BI analysis, to the extent that the consensus tree does not even resolve a monophyletic Spatangoida. This might be a consequence of support for much older dates for many nodes: crown group atelostomates shift to become more than 22 Myr older, and total group atelostomates more than 30 Myr older, with the addition of molecular data. Increased molecular sampling within Aulodonta, Atelostomata and Cidaroida should therefore be a future priority.

### Body Size Macroevolution

The biovolumes represented in our phylogeny span more than five orders of magnitude, from the minute fibulariids to the gigantic echinothurioids, attesting to the versatility of the echinoid body plan. Despite this huge variability, body size also exhibits a strong phylogenetic signal (Pagel’s λ = 0.90, *P* = 4.65 x 10^-11^). The analysis of the trajectory of body size disparity through time however reveals a complex picture (Fig. 6). Even though relative disparity follows the expectations of a BM process during the earliest part of the echinoid evolutionary history, relative disparity is consistently higher than expected from the Jurassic onwards, driving a significant deviation from null expectations (Rank envelope test, *P* = 0.016). This deviation is not consistent with an “early burst” of disparity that often characterizes paleontological datasets (Foote 1995; Cooper and Purvis 2010; Hughes et al. 2013), as this would result in deviations towards lower values.

**Figure 6:**
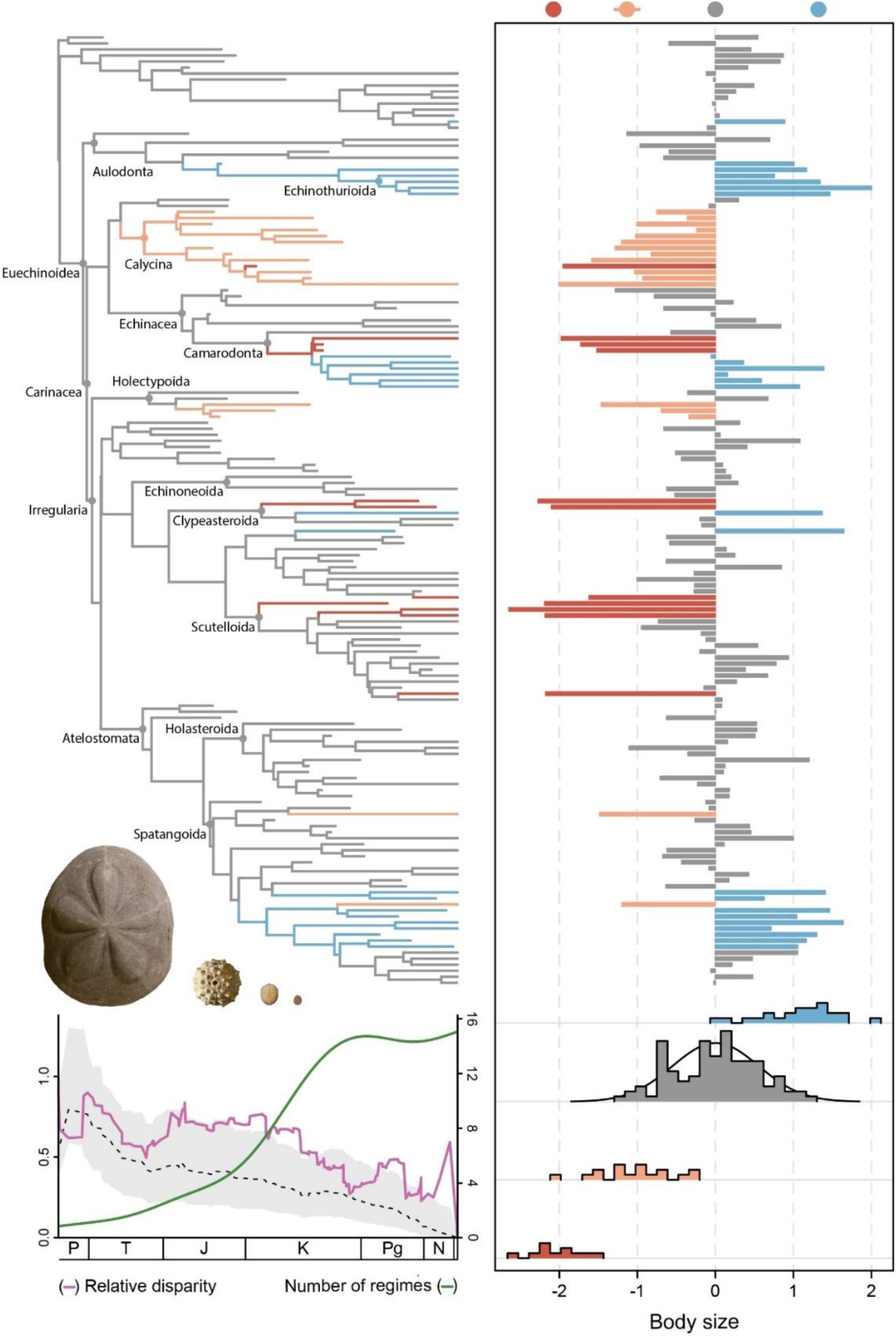
Macroevolutionary history of echinoid body size. The four selective regimes supported by the OUMVAZ model are painted on the mcc tree (left; terminal names are shown in Fig. S18 of SI File 4), body sizes of terminals are shown on the right. Color dots represent the inferred location of peaks (θ). Body size is log_10_-scaled and shown relative to the value of the average peak (4.10, see Table 3). The distribution of body sizes in each regime are shown using histograms, along with the expected distribution for the average regime (grey) at equilibrium. The DTT plot on the bottom left corner shows a significant deviation of the observed disparity (pink) compared to the expectations under BM (the dashed line is the median trajectory of simulations; grey areas denote the 95% confidence interval). The green line shows the number of active regimes operating in the phylogeny through time, smoothed using a generalized additive model. The echinoids shown include a member of each of the four adaptive peaks. From left to right (photo credit): *Clypeaster subdepressus* (J. Utrup), *Hemicidaris intermedia* (J. Utrup), *Palaeostoma mirabile* (R. Mooi) and *Echinocyamus crispus* (T. Grun).

Positive deviations in DTT plots indicate that subclades contain a larger share of the total morphological disparity than expected. This pattern has been shown to arise from either increases in the rate of evolution with time or an exploration of trait space affected by constraints and/or convergences, such as is generated by OU processes (Colombo et al. 2015). To explore the potential drivers of this pattern we compared the fit of multiple models of macroevolution. We found no evidence of changes in the overall rate of evolution with time: the EB model performed worse than the time-homogenous BM (Table 2). All non-uniform models fitted the data better than those assuming a single process, revealing an intricate history of morphological evolution. From the set of non-uniform processes, the SURFACE algorithm returned a highly complex OUM model, describing an adaptive landscape with seven distinct peaks. However, this model contained unrealistic selective regimes, characterized by peaks well outside the range of body sizes explored by echinoids (e.g., suggesting optimum test diameters > 2.5 m; Fig. S16 of SI File 4). This is likely a consequence of SURFACE fixing common σ^2^ and α parameters across the entire tree, leaving rate heterogeneity to be accommodated exclusively by θ values. However, relaxing this assumption is not straightforward; more realistic models with multiple selective regimes are generally too complex to be fit (Fig. S17 of SI File 4), a common problem of OUM models with multiple σ^2^ and/or α parameters (Beaulieu et al. 2012; Benson et al. 2018).

To resolve this problem, we used a novel approach that relies on continuing the backwards phase of the SURFACE algorithm, forcing it to merge independent selective regimes even when this results in suboptimal OUM models (i.e., with higher AICc values). We refer to this as the extended phase of SURFACE (Fig. S17 of SI File 4), and code to implement it can be found in SI File 10. The simpler models obtained this way were used as starting points for the optimization of regime-specific σ^2^ and α parameters using OUwie. The best OUMVA and OUMVAZ models obtained were incorporated into the final model comparison shown in Table 2, and found to fit the data much better than all other options explored. In particular, an OUMVAZ model with four selective regimes (Figs. 6 and S18 of SI File 4) outperforms other models to the point that it attains virtually maximum AICc weight (Table 2; similar results were obtained for randomly sampled posterior trees, Fig. S19 of SI File 4). This model consists of an ancestral regime that became established even before the origin of the crown group. This regime can be considered to constitute the evolution of echinoids with an average body size: it not only permeates much of the clade’s evolutionary history (67% of tips and 72% of total tree length; Table 3), but it also describes attraction to an optimum value of 4.10 (approx. 30 mm in diameter x 14 mm in height), which lies very close to the average body size of 4.01 found across all sampled species (Table 3).

The evolutionary dynamics of 18 lineages deviate from the behavior expected of this regime (Fig. 6). Their trajectories are better explained by invoking three other selective regimes, describing evolution towards adaptive peaks that differ from the ancestral one by more than an order of magnitude (Table 3). Each of these three regimes are encountered convergently by multiple lineages (between 4 and 7 independent times), representing body sizes accessible by most clades of regular and irregular echinoids. These regimes are also characterized by relatively stronger strengths of attraction than the average regime (Table 3), although all phylogenetic half-lives are on the order of 10-25 Ma. This indicates that radical innovations in body size were not produced by sudden bursts of morphological change but by relatively slow—yet directional— macroevolutionary processes. Despite these long time-scales, most lineages have had enough time to adapt and reach their optimal sizes. These regimes have contributed significantly to the overall morphological disparity of echinoids (Fig. 6 and S20 of SI File 4), allowing entire lineages of both gigantic and miniaturized forms to diversify well outside the range of sizes achievable within the average regime. In fact, the variance of body sizes across species (V_total_ = 0.83) almost triples the variance observed among species in the average regime (V_average_ = 0.29), which is very close to the expected variance of this regime at equilibrium (*σ*^2^⁄2*α* = 0.31; Hansen 1997). This shows that much of the observed disparity can be attributed to clades that have managed to escape the constraints of a regime that has exhausted its ability to produce morphological novelty.

Given the strong relationship between body size and multiple aspects of ecology, physiology and life history (Peters 1986; Calder 1996; Smith and Lyons 2013), the origin of new regimes is likely associated with significant ecological innovations. The pattern of accumulation of regimes through time can therefore provide a proxy for the breadth of occupied ecospace. From this perspective, echinoids seem to have undergone relatively little ecological change early on, followed by a drastic increase in the number of active regimes throughout the Jurassic and Cretaceous (Fig. 6). Ecological innovation decelerates once again in the Cenozoic, with the rate of origin of regimes decaying to eventually match that of their extinction (Fig. S21 of SI File 4). The evolutionary history of echinoid body size is therefore characterized by a late increase in disparity (possibly linked to a late expansion of ecospace), achieved through the repeated evolution of multiple lineages towards extreme morphologies.

## Discussion

### Phylogenomics and Total-Evidence Dating

A comprehensive approach to phylogenetic reconstruction requires integration across the independent lines of evidence left behind by the process of evolution (Lee and Palci 2015; Pyron 2015). Methodological advances such as TED inference provide new ways to leverage information from distinct data sources. The recent increase in genomic resources for echinoids, coupled with their extensive fossil record and morphological complexity, makes them an ideal study system for such synthesizing studies. We have here undertaken such an approach, resulting in an updated estimate of the phylogenetic relationships and divergence times for the main lineages of crown group Echinoidea.

Despite the evident benefits of genome-scale data for phylogenetic reconstruction, the computational burden imposed by many models of divergence-time estimation requires reduced datasets. This means that most TED analyses that explicitly incorporate morphological and stratigraphic information have been unable to tap into the vast molecular resources available in the genomic era. Loci subsampling (or “gene shopping”) is not only a necessity for TED analyses, but has also become a standard phylogenomic procedure given systematic biases in complex and heterogenous molecular datasets (Molloy and Warnow 2018; Smith et al. 2018; Mongiardino Koch 2019). Loci are generally selected for either high phylogenetic signal or low systematic bias, a practice that is relatively straightforward if there is evidence that the phylogenetic question addressed might suffer from issues related to either one of these. However, it remains unclear how different these two approaches are and which one is preferable in the absence of readily diagnosable issues. Even when subsampling is performed in a way that accounts for multiple gene properties, this is normally performed in a stepwise fashion whose precise order is hard to justify (e.g., Whelan et al. 2015; Smith et al. 2018; see Kocot et al. 2017 for a noteworthy exception).

We developed a novel pipeline for the selection of loci that is based on a multivariate analysis of gene properties (Fig. 3). We show how this approach selects loci in a way that simultaneously maximizes phylogenetic signal and minimizes sources of systematic bias, as well as favoring increased clock-like behavior (Fig. 4). This is achieved by explicitly accounting for the correlation structure between different gene properties, which outperforms results obtained by optimizing these variables individually (Fig. 3). Our approach results in new dimensions that capture variability in both rate of evolution and overall phylogenetic usefulness across loci (Figs. 3 and S8 of SI File 4). The set of selected genes supports a topology identical to that obtained with the complete dataset, and all nodes attain high support even when the molecular dataset is reduced to 2% of its original size (Figs. S10-S12 of SI File 4). Our method is an efficient way to obtain informative subsamples of loci from genome-scale datasets, and especially useful for combining phylogenomic resources with methods of inference that incorporate fossil data.

The most striking result of our TED analysis is the extent to which different sources of evidence complement each other to reduce topological conflict. A clear example of this is the position of Echinothurioida, a lineage of traditionally uncertain affinities given conflicts between morphological and molecular evidence (Kroh and Smith 2010; Mongiardino Koch et al. 2018). However, our reanalysis of morphology in combination with stratigraphic information resolves this clade in the same position as molecular data (Fig. 2). The implementation of a Bayesian clock thus seems to increase the topological accuracy of inference from morphological evidence. Another striking example of the benefits of combined approaches is the shift in position of many lineages that lack molecular data. Even though our TED analysis includes molecular data for only 21 lineages, the resulting topological reorganization impacts the position of many other lineages. Previous studies have shown that molecular data can modify the position of fossils (e.g., Wiens et al. 2010; Arcila et al. 2015), but these topological changes cannot be corroborated with independent data sources, leading some to doubt whether they represent improvements in phylogenetic accuracy (McMahan et al. 2015). In our morphological analyses (Figs. 2 and S5-7 of SI File 4), echinoneoids are strongly supported as the sister group to the remaining crown group irregulars, in line with previous morphological studies (Smith 1981; Kroh and Smith 2010). However, the incorporation of molecular data for other terminals resolves this clade as sister to crown group neognathostomates (Fig. 5). This topological change can be corroborated against independent sources of evidence, and is found to agree with a recent molecular study (Lin et al. 2019). Thus, combining different data sources serves to resolve two of the most prominent cases of uncertainty in the higher-level phylogeny of echinoids.

### Phylogeny of Echinoidea

One of the novel results of our analyses is the placement of a number of Permian and Triassic taxa (*Eotiaris*, *Serpianotiaris* and *Triadotiaris*), as stem group echinoids. These taxa had previously been regarded as amongst the earliest crown group echinoids (Smith 1990; Smith and Hollingworth 1990; Smith 2007; Kroh and Smith 2010), with recent treatments classifying them as stem cidaroids. *Eotiaris* in particular includes Permian species considered to be the earliest crown group echinoids, and used to constrain the age of the clade (Smith and Hollingworth 1990; Smith et al. 2006; Thompson et al. 2015; Thompson et al. 2017). Our analyses thus imply paraphyly of Cidaroidea (as well as Cidaroida) with respect to euechinoids. Even though this topology excludes all well-known clades of Permian to Middle Triassic fossils from the crown group, the inferred time of origin of the clade remains in the Late Paleozoic, in line with previous estimates (Smith et al. 2006; Nowak et al. 2013; Thompson et al. 2017).

Traditionally, the Permo-Triassic extinction event was considered to have played a major role in shaping the macroevolutionary history of echinoids, with only one or two lineages crossing the boundary (Kier 1977; Smith 1984; Smith and Hollingworth 1990; Erwin 1994; Twitchett and Oji 2005). However, recent work on echinoids and other echinoderm groups, suggests that the end-Permian extinction had a lower impact on echinoderm macroevolution (Thuy et al. 2017; Reich et al. 2018; Thompson et al. 2018). Our topology and divergence times are in line with these findings, indicating that multiple lineages of echinoids survived the end-Permian mass extinction, including members of Miocidaridae, Serpianotiaridae, Triadotiaridae and the three main clades of crown group echinoids (cidaroids, aulodonts and carinaceans).

The novel placement of these Permian and Triassic taxa also has important implications for reconstructing the morphological evolution associated with the origin of the crown. The sustained lack of morphological innovation along the cidaroid evolutionary lineage (Hopkins and Smith 2015) has historically complicated a natural delimitation of this clade. Many early systematists allied extant cidaroids with most or even all of the Paleozoic taxa now recognized to be stem group echinoids (Mortensen 1928; Philip 1965; Durham 1966a and references therein). Later phylogenetic work was crucial to disentangling true cidaroids from stem group echinoids (Jensen 1981; Smith 1981, 1984), and identifying the morphological changes associated with the origin of the crown. However, many of these changes no longer represent synapomorphies of crown group Echinoidea in our topology, but are rather innovations that predate its origin. These include the presence of a perignathic girdle (the skeletal protrusions that provide attachment sites for the support muscles of the Aristotle’s lantern), the reduction in the number of plate columns to 20 (two per ambulacral and interambulacral region), and the suturing of interambulacral plating (Jackson 1912; Smith 1980, 1984; Smith and Hollingworth 1990; Thompson and Ausich 2016; Thompson et al. 2019). The loss of imbrication along the ambulacral-interambulacral suture seems to represent the most conspicuous synapomorphy of crown group Echinoidea as redefined here (a trait that secondarily reverses among echinothurioids and other aulodonts, see Fig. S22 of SI File S4).

As previously mentioned, the position of Echinothurioida (and thus the composition of the major euechinoid clades) is one of the most contested issues in echinoid phylogenetics. For long, the unique morphological features of echinothurioids have been interpreted as either plesiomorphic traits indicative of an origin early, or autapomorphic conditions of a highly derived clade (Woodward 1863; Gregory 1897; Mortensen 1935, 1940; Durham and Melville 1957; Fell 1966; Smith 1981; Kroh and Smith 2010; Mongiardino Koch et al. 2018; Kroh 2020). Even though both hypotheses were originally formulated on the basis of morphological evidence, the debate has persisted as an example of conflict between morphological and molecular evidence (Kroh and Smith 2010; Mongiardino Koch et al. 2018). Our analyses are the first to place echinothurioids in a consistent position regardless of data choice (Figs. 1, 2, 5), ending over 150 years of controversy, and confirming a basal split of euechinoids into Aulodonta and Carinacea. The echinothurioid condition of imbricate plating (shared to some extent with other aulodonts; Durham and Melville 1957; Fell 1966) has evolved secondarily and is not homologous with that of Paleozoic stem group echinoids (see Fig. S22 of SI File S4). This is in line with evidence from interambulacral plate microstructure (Smith 1984), as well as the morphology of pelanechinids (such as *Pelanechinus corallinus*) where imbricate plating is only present adapically (Smith 2015), and could represent a transitional morphology between a rigidly plated ancestral euechinoid and the flexible test of echinothurioids.

Echinoneoida also changes position in our TED (Fig. 5, a topology congruent with recent molecular estimates; Lin et al. 2019), leaving Atelostomata as the sister clade to all other crown group irregulars. This shift is likely a consequence of the radical reorganization undergone by “cassiduloids”, one of the most problematic groups of echinoids from a phylogenetic perspective. The high levels of homoplasy and character exhaustion that characterize their morphological evolution has rendered the clade a taxonomic wastebasket (Suter 1994; Souto et al. 2019). The exact delineation of the group has varied through time, but traditionally included the diverse paraphyletic assemblage of neognathosomates subtending the clade comprised of extant Clypeasteroida + Scutelloida. In an attempt to establish a more natural classification of this lineage, Kroh and Smith (2010) reclassified oligopygids (including *Oligopygus* and *Haimea*), faujasiids (here represented by *Faujasia* and *Stigmatopygus*) and plesiolampadids within Clypeasteroida (*sensu* Agassiz 1873, now reclassified within Clypeasteroida *sensu* Mongiardino Koch et al. 2018; Kroh 2020), and placed echinolampadids in their own order. Some features of this classification have since been contested (Souto et al. 2019), and are not supported by our time-calibrated reanalysis of morphology (Fig. 2).

The classification of “cassiduloids” was further complicated by molecular results which placed cassidulids and echinolampadids (either individually or as a clade) as the sister group to the sand dollars (Littlewood and Smith 1995; Smith et al. 1995; Smith et al. 2006; Nowak et al. 2013; Smith and Kroh 2013; Thompson et al. 2017; Mongiardino Koch et al. 2018). Although the phylogenomic analysis of Mongiardino Koch et al. (2018) revealed a strong phylogenetic signal allying echinolampadids with sand dollars, this left the affinities of other “cassiduloids” uncertain. The results of our TED analysis (Fig. 5) show that identifying the close relationships of some “cassiduloids” to sand dollars was necessary to establish a natural subdivision of the clade. Two lineages of “cassiduloids” (whose monophyly attain pp > 0.99) are nested within the clade that contains sand dollars and sea biscuits. One of these includes oligopygids and conoclypids, lineages that share the possession of a lantern in the adults with both scutelloids and clypeasteroids. This clade was recognized in some classifications under the names Conoclypina (Durham and Melville 1957) or Oligopygoida (Rose 1982), and considered either a member or the sister group to the “cassiduloids” (Mortensen 1948a; Philip 1963, 1965; Kier 1967, 1974; Smith 1981, 2001). The second clade includes all extant “cassiduloids” as well as faujasiids, clypeolampadids, pliolampadids and plesiolampadids. This clade lacks the Aristotle’s lantern in adult forms (although it is present in juveniles of all extant lineages; Gladfelter 1978; Ziegler et al. 2012), and agrees with the composition of Cassiduloida by Souto et al. (2019), which is expanded by the addition of *Apatopygus* and taxa not included in their analysis. We suggest these two clades should bear the names Oligopygoida Kier, 1967 and Cassiduloida Agassiz & Desor, 1847, respectively (with the latter subsuming Echinolampadoida Kroh & Smith, 2010). Alongside Clypeasteroida and Scutelloida, these clades constitute the four lineages of a restructured crown group Neognathostomata, all of which originated during the Cretaceous.

The third clade of “cassiduloids” is distantly related to the rest, forming the extinct sister group to echinoneoids + crown group neognathostomates (Fig. 5). This third clade comprises the bulk of Mesozoic “cassiduloid” diversity. It includes most of the taxa originally assigned to Nucleolitoida Hawkins, 1920 (clypeids, galeropygids, nucleolitids and pygaulids; Hawkins 1920) and we suggest resurrecting this name. A clade of archiaciids and claviasterids, as well as other lineages variously classified as among the earliest fossil neognathostomates or atelostomates (Durham and Melville 1957; Kier 1962, 1966; Barras 2007; Kroh and Smith 2010) also appear to belong to this clade. Surprisingly, the extant *Apatopygus recens*—regarded as a close relative of the Mesozoic *Nucleolites* (Hawkins 1920; Mortensen 1948a; Kroh and Smith 2010; Souto et al. 2019)—does not resolve within this clade. Although the relationship between apatopygids and nucleolitids has been questioned previously based on morphological differences and their long separation in time (Kier 1962, 1966; see also Smith 2001), this result requires testing with molecular data. Regardless of the position of *Apatopygus*, our splitting of the “cassiduloids” implies that previous discussion of their evolution and diversification (Kier 1966, 1974; Suter 1988; Mooi 1990a; Souto et al. 2019) conflated processes operating on unrelated clades. It also reveals that some of the synapomorphies thought to support a monophyletic clade composed exclusively of sand dollars and sea biscuits, such as the presence of internal test reinforcements, are the result of convergent evolution (Fig. S22 of SI File 4). Recent studies show that these structures are necessary for the stability of flat tests (Grun et al. 2018), and they likely evolved independently in different flattened lineages. Other traits, however, appear to have originated in the last common ancestor of sand dollars and sea biscuits, and were subsequently lost by some of its descendants (e.g., the presence of Aristotle’s lantern in adults; Fig. S23 of SI File 4).

### Approaches to Macroevolutionary Inference

Research on the evolution of quantitative traits across deep timescales is constrained by the set of models available. Historically, the macroevolutionary toolkit was centered around BM and some extensions capable of accounting for rate variation through time, active trends and attraction to optimum values. However, there has been a recent expansion of the types of evolutionary dynamics that can be modeled, including bounded explorations of traitspace (Boucher and Démery 2016), trajectories punctuated by rapid bursts of change (Landis and Schraiber 2017) and macroevolutionary landscapes that can represent directional and disruptive selection (Boucher et al. 2017). Complex OUM models capable of detecting heterogeneities in evolutionary processes across clades have also been developed (Beaulieu et al. 2012; Ingram and Mahler 2013; Khabbazian et al. 2016; Bastide et al. 2018). Nonetheless, there have been few attempts at comparing the fit of non-uniform OUM processes to the expanded repertoire of uniform models now available.

Our results show that, even though this new generation of models can explain evolutionary patterns better than its predecessors (Table 2; see also Landis and Schraiber 2017), the history of echinoid body size seems to be too complex to be explained by a uniform process. This echoes the results of other explorations of body size evolution in comparative datasets (Price and Hopkins 2015; Benson et al. 2018; Godoy et al. 2019). This complexity is better accommodated by non-uniform macroevolutionary models, all of which attained higher AICc weights than those that assume that a single process governed evolution across the entire phylogeny (Table 2). Our results show that even relatively simple non-uniform models that only allow for rate heterogeneity across lineages (BMS) can fit the data substantially better than uniform models. However, we also demonstrate that the evolution of echinoid body size is better characterized by incorporating attractive forces operating on a macroevolutionary adaptive landscape. This landscape seems to have been relatively stable throughout the history of the clade, with multiple lineages independently evolving similarly extreme morphologies. OUM models are designed to accommodate such recurrent evolutionary outcomes, another factor that likely contributes to the success with which they model echinoid body size evolution.

A common concern with OUM processes (especially the SURFACE algorithm) is their tendency to favor overly complex models (Ho and Ané 2014; Khabbazian et al. 2016; Davis and Betancur-R 2017). Our results show that this might be a consequence of the unrealistic assumption of a common rate of evolution and common force of attraction operating across the entire tree. Relaxing this assumption by optimizing regime-specific σ^2^ and α parameters on suboptimal OUM models confirms that simpler models (i.e., those with fewer adaptive peaks than the one supported by SURFACE) can fit the data better (Fig. S19 of SI File 4). This method allowed us to identify an OUMVAZ model describing a four-peak adaptive landscape which attained almost absolute AICc weight. This approach is a promising way to mitigate the overfitting tendencies of SURFACE while generating more biologically realistic models, especially as other methods to fit OUM models without specifying the location of regime shifts cannot be applied to non-ultrametric trees (e.g., Khabbazian et al. 2016; Bastide et al. 2018).

### Evolution of Echinoid Body Size

The macroevolutionary history of body size across the echinoid crown group was previously synthetized by Kier (1974). He suggested that early post-Paleozoic echinoids were relatively small and, from the later part of the Jurassic onward, different groups of regular and irregular echinoids independently evolved towards larger sizes, culminating in gigantism among living echinothurioids and some clades of spatangoids. Our analysis confirms these observations, while providing a more nuanced picture of the macroevolutionary dynamics of this trait.

Much of the evolutionary history of echinoid body size can be described by attraction to a single adaptive peak. This regime predates the origin of the crown group, and permeates much of the earliest history of the clade, including the origin of most of the major lineages (Figs. 6 and S18 of SI File 4). Some groups with limited body size disparity, such as holasteroids, echinoneoids and nucleolitoids, appear to have been unable to escape the constraints imposed by this regime. At equilibrium (a condition likely to be met today as more than 10 phylogenetic half-lives have elapsed), this regime would produce body sizes spanning a little more than two orders of magnitude (95% confidence interval = 3.01 – 5.19).

However, the observed range of echinoid body size spans more than five orders of magnitude, 295 times larger than expected under the average regime. This remarkable disparity was generated through multiple pulses of directional evolution within specific lineages bridging entire orders of magnitude of body size. The dynamics of these clades are better explained by the presence of three additional peaks in macroevolutionary adaptive landscape, resulting in attraction to gigantic, small-bodied and miniature sizes (Table 3). Occupation of these peaks likely entailed significant innovations, allowing clades to traverse apparently unstable regions of traitspace. Even though these peaks have been encountered repeatedly, we do not consider shifts to the same peak to represent true morphological convergences, as is normally interpreted when employing multivariate characterizations of morphotypes (e.g., Mahler et al. 2013; Davis and Betancur-R 2017; Ceccarelli et al. 2019). In our case, shifts to the same peak denote events of innovation that share similar evolutionary tempo and outcome, but the clades involved are not expected to share ecological or morphological similarities. Nonetheless, these peaks appear to represent stable macroevolutionary outcomes, accessible by multiple lineages over hundreds of millions of years. Although Kier (1974) only mentioned body size increase, innovations towards smaller sizes appear to have been relatively more common (63% of regime shifts).

Even though there is a coarse correspondence between body size and ecological niche (see Benson et al. 2018), lineages that have evolved directionally across orders of magnitude of body size likely innovated substantially from an ecological perspective. This is evident for many of the lineages that have undergone regime shifts: from the miniaturized scutelloids that feed on the organic matter surrounding individual substrate particles (Telford et al. 1983) to the gigantic spatangoids that rework sedimentary environments through their burrowing and feeding activities (Hollertz and Duchêne 2001), these clades have forged entirely new interactions with their environment. Thus, the temporal pattern of accumulation of regime shifts can provide some indication of the breadth of the ecospace occupied through time. There are of course some aspects of ecological history that this approach will miss, as many changes in niche will not impact overall body size (many lineages of ecologically specialized echinoids still occupy the average regime), but there is also no evidence that detectable innovations are temporally biased.

From this perspective, the macroevolution of echinoid body size involved a relatively late onset of innovation, leading up to a dramatic expansion of ecospace during the Jurassic and Cretaceous (Figs. 6 and S21 of SI File 4). An increase in morphological and ecological innovation among Jurassic echinoids was also previously recognized using both discrete characters and geometric morphometrics (Hopkins and Smith 2015; Boivin et al. 2018), data types that can support contradictory macroevolutionary patterns (Mongiardino Koch et al. 2017). These previous studies recognized evolutionary innovations mainly among irregular echinoids, whereas our results show that multiple clades of regular echinoids (echinothurioids, calycineans and camarodonts) also contributed to this expansion of ecospace. The rate of origin of new regimes declined by the late Cretaceous, and has since dwindled to values comparable to those of the Triassic (Fig S21 of SI File 4).

### Conclusions

The development of phylogenomic datasets and methods, and improvements in the way the fossil record is incorporated in the inference procedure, are two of the most important recent advances in the field of systematics. However, these two areas of research have developed largely in isolation. Even though trees of extant taxa can now be inferred from thousands of loci, this represents only the first step in building a phylogenetic framework for clades with a good fossil record. Furthermore, the stratigraphic and morphological information provided by fossils can strongly influence the reconstruction of ancestral states, modes of macroevolution, diversification dynamics, divergence times and morphological topologies (Quental and Marshall 2010; Slater and Harmon 2013; Lee and Palci 2015; Pyron 2015; Mongiardino Koch and Parry 2019 and references therein). Integrating phylogenomics with the fossil record thus captures the evolutionary history of of clades to a degree unattainable by either individually.

Here we combine phylogenomic, morphological, stratigraphic and morphometric data for echinoids, and develop novel methods for both loci subsampling and macroevolutionary model fitting. This approach has yielded a revised phylogeny and timing of origin of the main lineages of crown group echinoids, resolved multiple phylogenetic uncertainties, and revealed compelling evidence that amalgamating distinct data sources improves topological accuracy. We have also been able to perform a detailed study on the evolutionary history of a trait of key ecological significance. We thus believe that our research showcases the unique opportunities that echinoids provide for integrative macroevolutionary research.

## Supplementary Material

Data available from the Dryad Digital Repository: http://dx.doi.org/10.5061/dryad.[NNNN].

## ACKNOWLEDGEMENTS

This project was enriched by discussions with Derek EG Briggs, Casey W Dunn, Andreas Kroh, Jesus Lozano-Fernandez, Rich Mooi and Luke A Parry. We would like to thank Kaylea Nelson and Benjamin Evans for computational assistance, Paul Simion for discussions regarding the software CroCo, and Nicolas Galtier and Jonathan Romingier for sharing transcriptome assemblies. We would also like to thank the managers and curators who gave us access to museum collections: Rich Mooi (CAS); Mark Renczkowski, Jessica Cundiff and Adam Baldinger (MCZ); Timothy Ewin (NHM); Kathy Hollis, William Moser and Mark Florence (NMNH); Charlotte Seid and Greg Rouse (SIO); Jessica Bazeley Utrup, Susan Butts, Lourdes Rojas and Eric Lazo-Wasem (YPM).

## Author Contributions

NMK and JRT designed the study and gathered the morphometric dataset. NMK performed all bioinformatic tasks and generated the phylogenomic dataset, JRT gathered the stratigraphic dataset. NMK ran all phylogenetic and macroevolutionary analyses, wrote all code and analyzed the data. NMK wrote the manuscript with contributions from JRT.

## SI File 2: Supplementary Tables

**Table 1:**
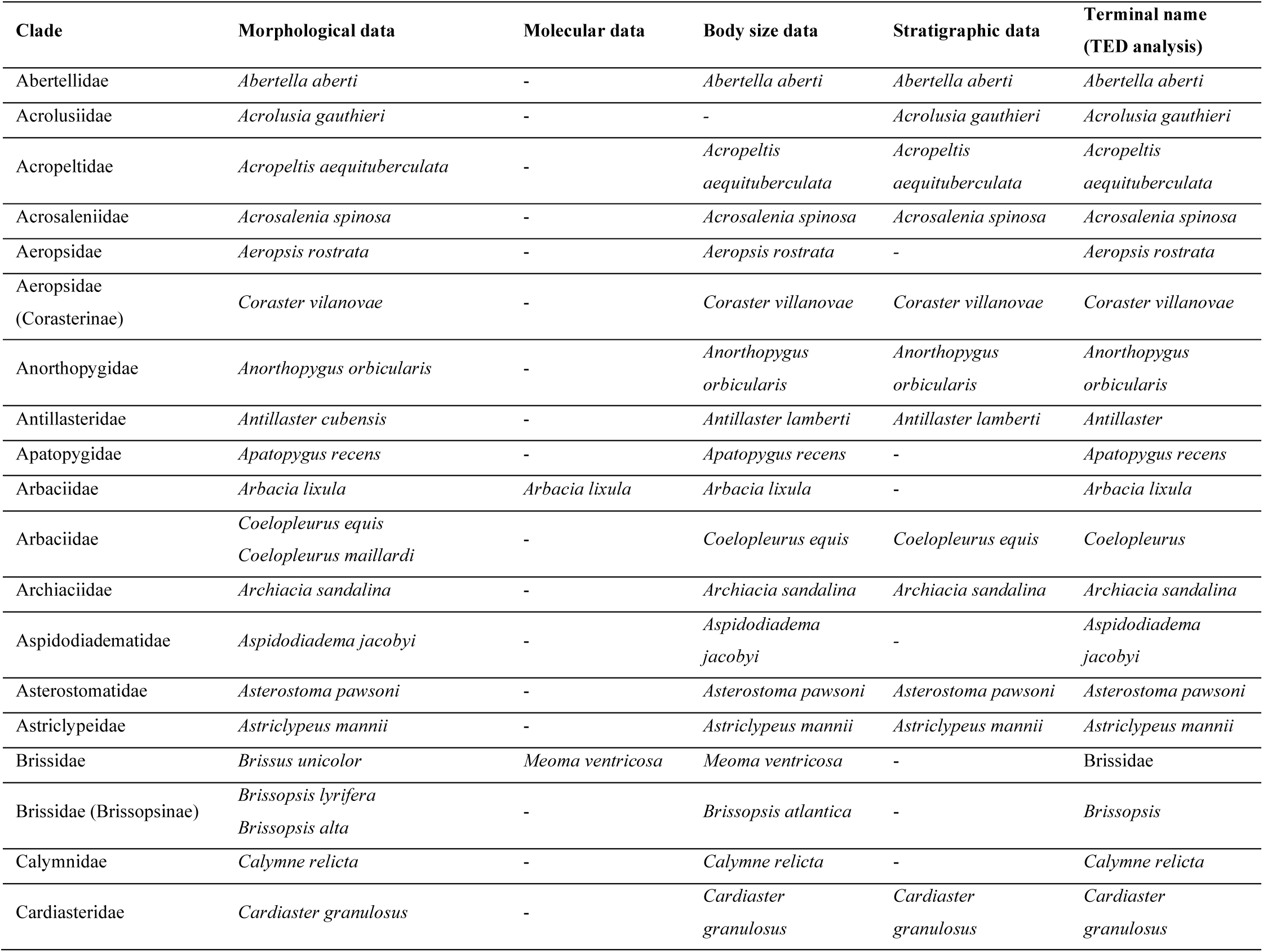
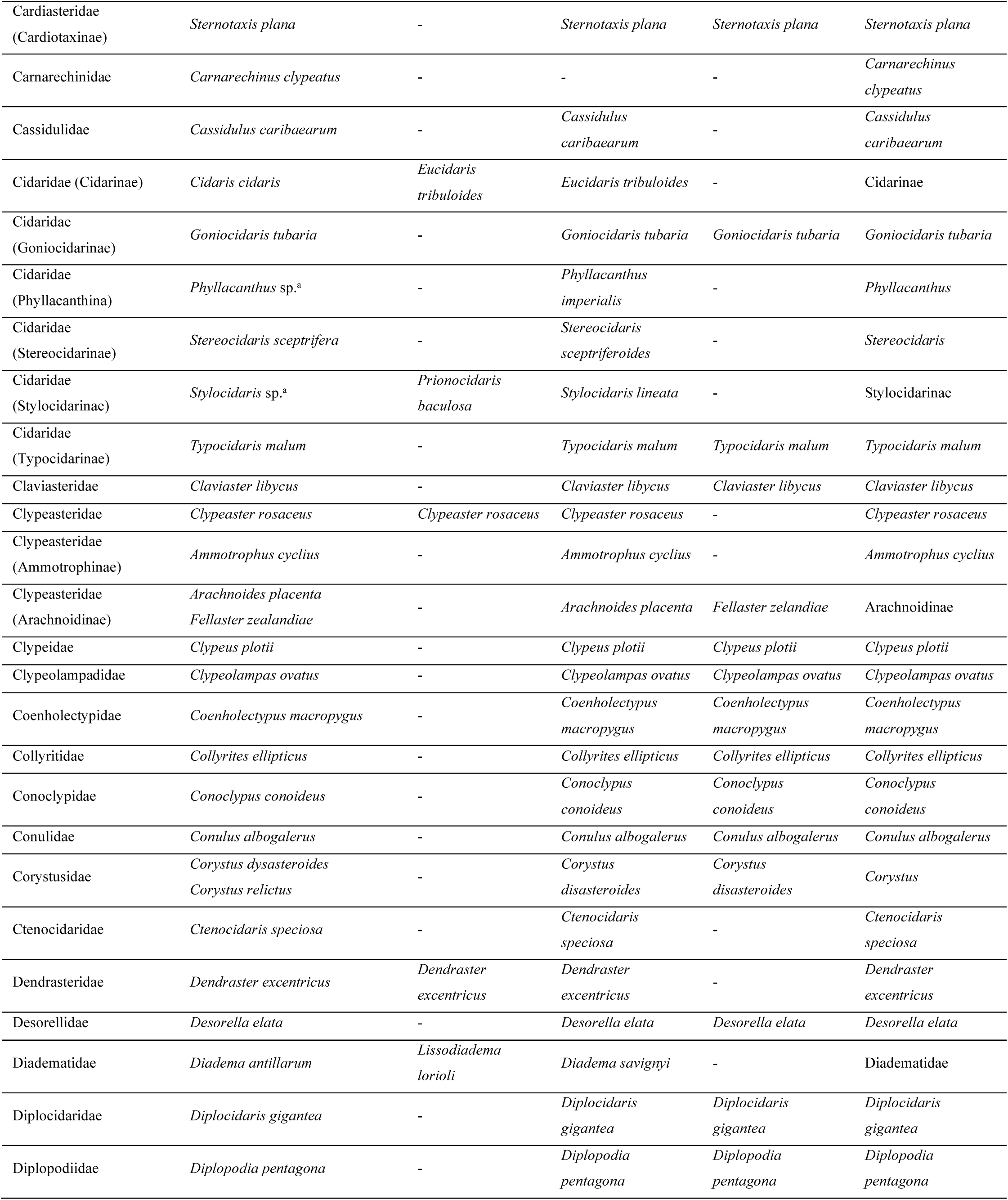
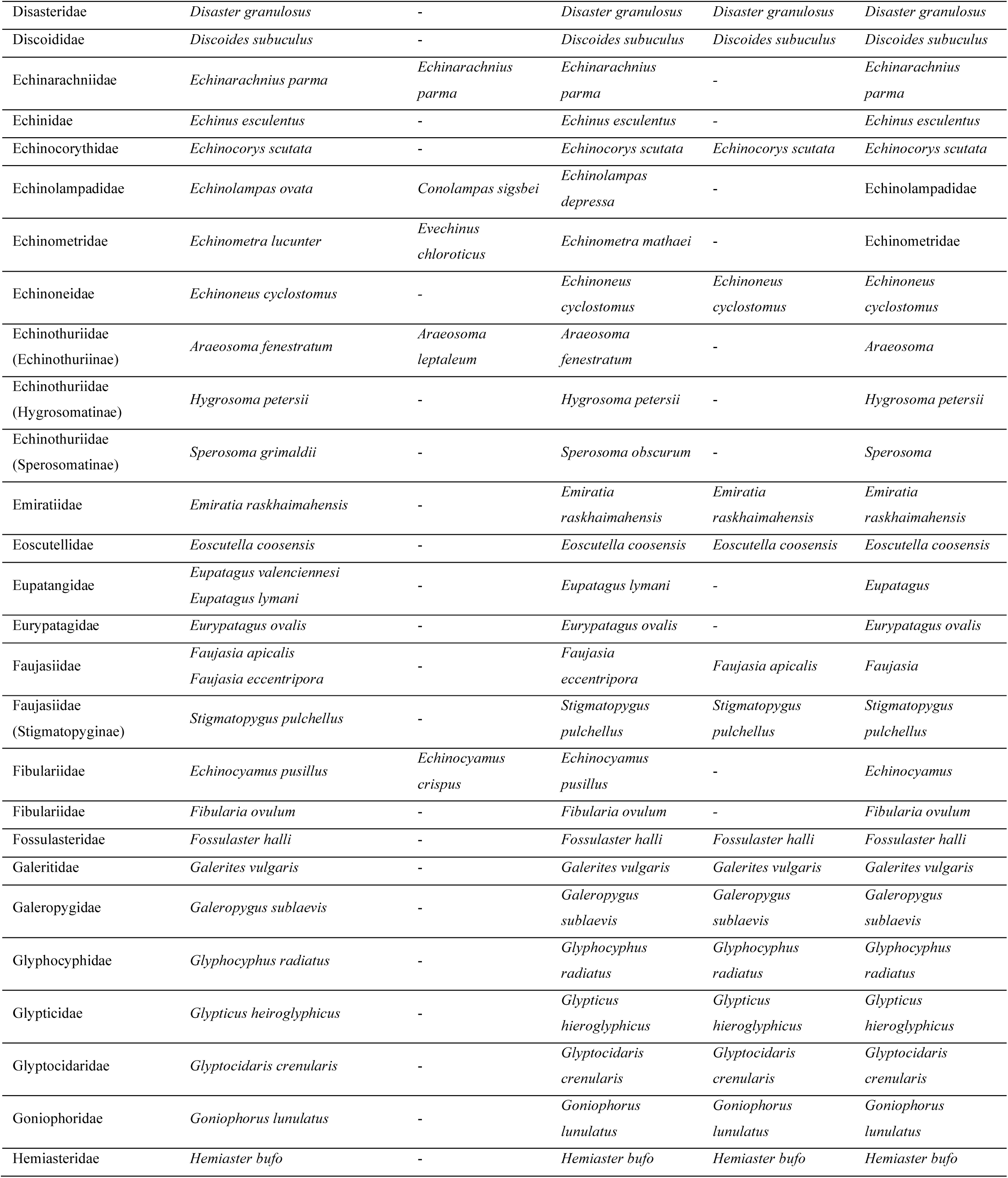
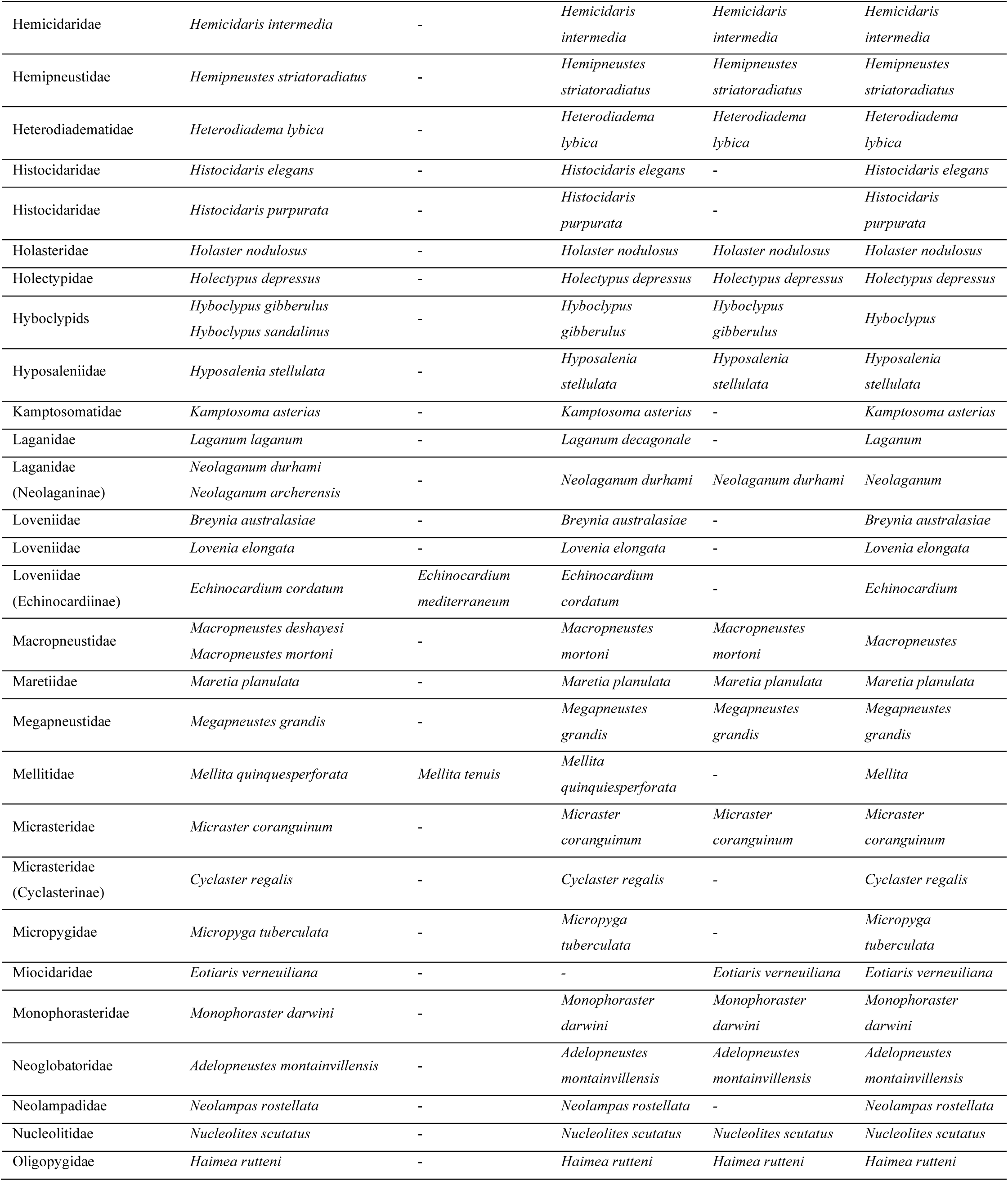
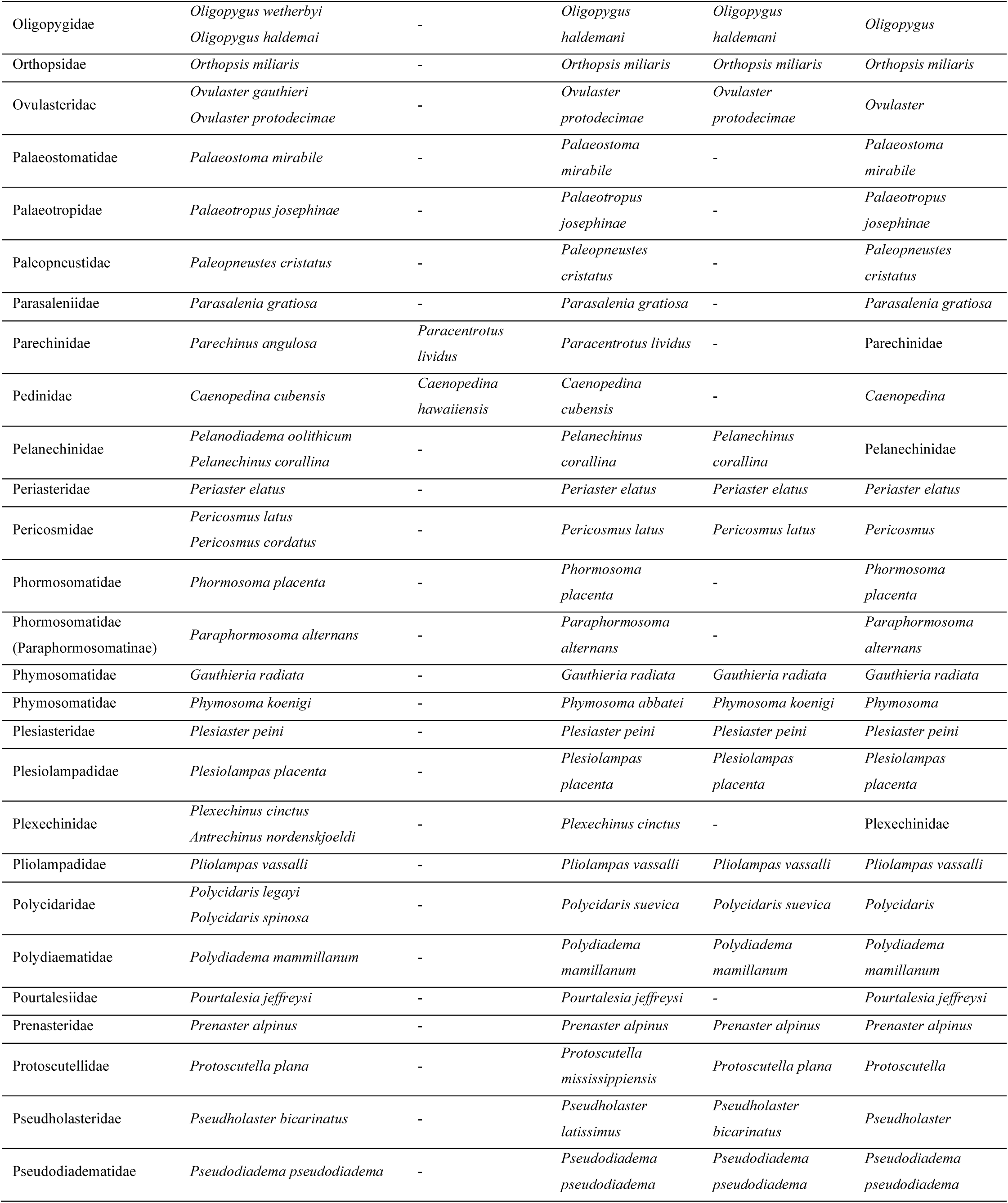
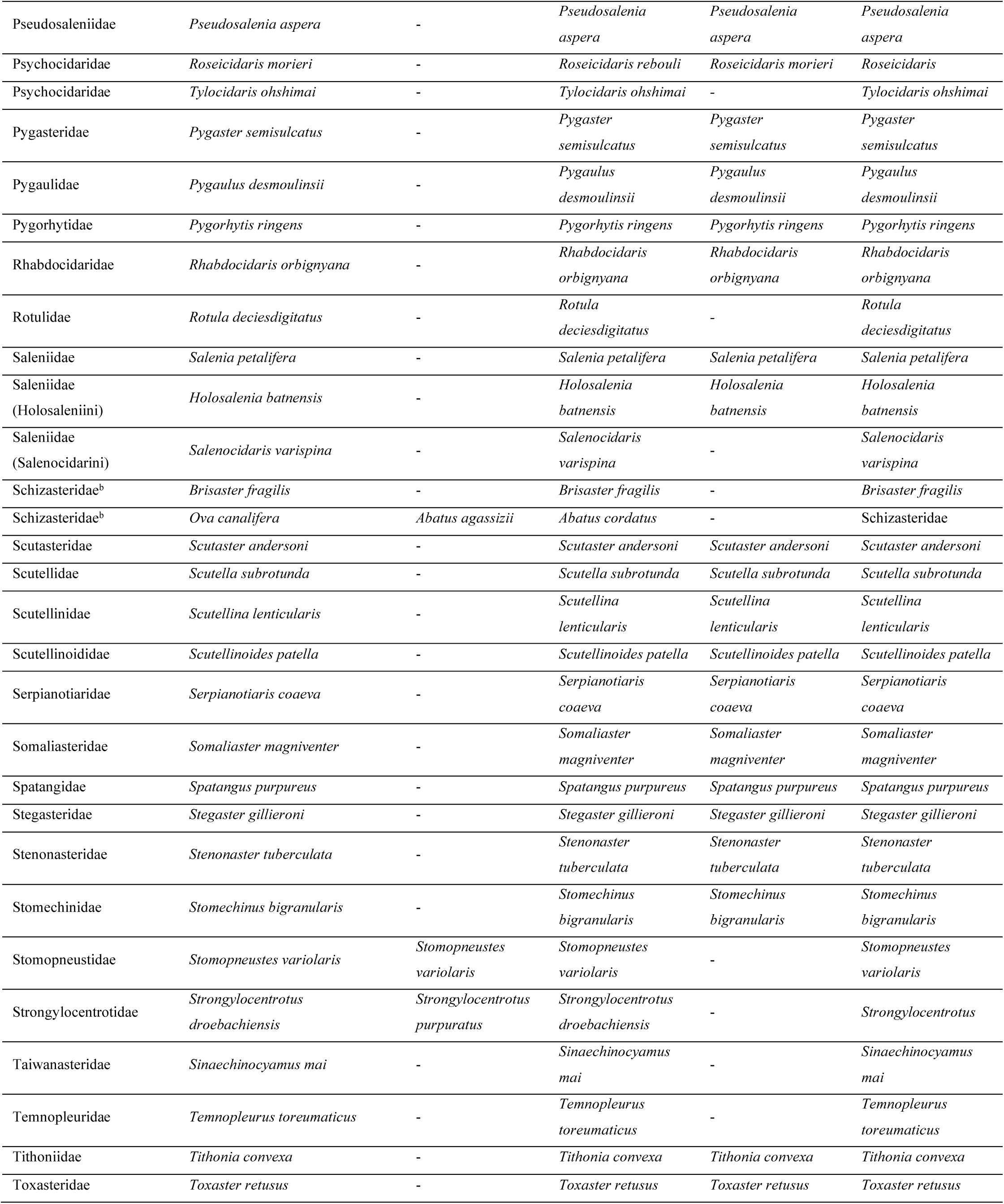
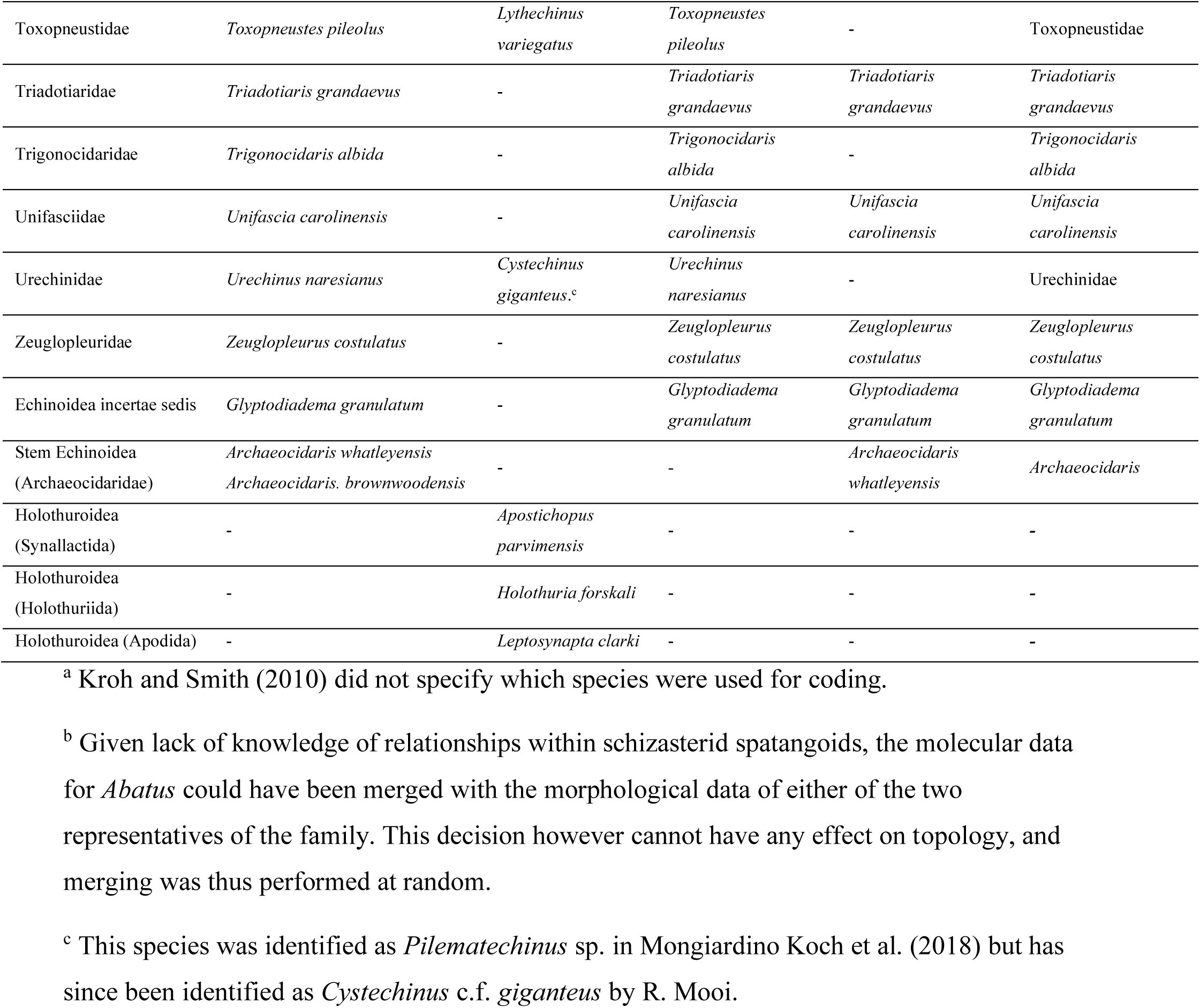
Species sampled in the molecular, morphological, morphometric and stratigraphic datasets. Here and throughout, the nomenclature used follows that of the World Echinoidea Database (Kroh and Mooi 2019), where citations to authorities and dates for scientific names can be found. The names of terminals for the total-evidence dated (TED) analysis are those of the least inclusive clade containing the species sampled across all datasets.

| Clade | Morphological data | Molecular data | Body size data | Stratigraphic data | Terminal name<br>(TED analysis) |
| --- | --- | --- | --- | --- | --- |
| Abertellidae | *Abertella aberti* | - | *Abertella aberti* | *Abertella aberti* | *Abertella aberti* |
| Acrolusiidae | *Acrolusia gauthieri* | - | - | *Acrolusia gauthieri* | *Acrolusia gauthieri* |
| Acropeltidae | *Acropeltis aequituberculata* | - | *Acropeltis*<br>*aequituberculata* | *Acropeltis*<br>*aequituberculata* | *Acropeltis*<br>*aequituberculata* |
| Acrosalenidae | *Acrosalenia spinosa* | - | *Acrosalenia spinosa* | *Acrosalenia spinosa* | *Acrosalenia spinosa* |
| Aeropsidae | *Aeropsis rostrata* | - | *Aeropsis rostrata* | - | *Aeropsis rostrata* |
| Aeropsidae<br>(Corasterinae) | *Coraster villanovae* | - | *Coraster villanovae* | *Coraster villanovae* | *Coraster villanovae* |
| Anorthopygidae | *Anorthopygus orbicularis* | - | *Anorthopygus*<br>*orbicularis* | *Anorthopygus*<br>*orbicularis* | *Anorthopygus*<br>*orbicularis* |
| Antillasteridae | *Antillaster cubensis* | - | *Antillaster lamberti* | *Antillaster lamberti* | *Antillaster* |
| Apatopygidae | *Apatopygus recens* | - | *Apatopygus recens* | - | *Apatopygus recens* |
| Arbaciidae | *Arbacia lixula* | *Arbacia lixula* | *Arbacia lixula* | - | *Arbacia lixula* |
| Arbaciidae | *Coelopleurus equis*<br>*Coelopleurus maillardi* | - | *Coelopleurus equis* | *Coelopleurus equis* | *Coelopleurus* |
| Archiaciidae | *Archiacia sandalina* | - | *Archiacia sandalina* | *Archiacia sandalina* | *Archiacia sandalina* |
| Aspidodiadematidae | *Aspidodiadema jacobyi* | - | *Aspidodiadema*<br>*jacobyi* | - | *Aspidodiadema*<br>*jacobyi* |
| Asterostomatidae | *Asterostoma pawsoni* | - | *Asterostoma pawsoni* | *Asterostoma pawsoni* | *Asterostoma pawsoni* |
| Astriclypeidae | *Astriclypeus mannii* | - | *Astriclypeus mannii* | *Astriclypeus mannii* | *Astriclypeus mannii* |
| Brissidae | *Brissus unicolor* | *Meoma ventricosa* | *Meoma ventricosa* | - | Brissidae |
| Brissidae (Brissopsinae) | *Brissopsis lyrifera*<br>*Brissopsis alta* | - | *Brissopsis atlantica* | - | *Brissopsis* |
| Calymnidae | *Calymne relictia* | - | *Calymne relictia* | - | *Calymne relictia* |
| Cardiasteridae | *Cardiaster granulosus* | - | *Cardiaster*<br>*granulosus* | *Cardiaster*<br>*granulosus* | *Cardiaster*<br>*granulosus* |
| MONGIARDINO KOCH & THOMPSON |  |  |  |  |  |
| Cardiasteridae<br>(Cardiotaxinae) | *Sternotaxis plana* | - | *Sternotaxis plana* | *Sternotaxis plana* | *Sternotaxis plana* |
| Carnarechinidae | *Carnarechinus clypeatus* | - | - | - | *Carnarechinus clypeatus* |
| Cassidulidae | *Cassidulus caribaeorum* | - | *Cassidulus caribaeorum* | - | *Cassidulus caribaeorum* |
| Cidaridae (Cidarinae) | *Cidaris cidaris* | *Eucidaris tribuloides* | *Eucidaris tribuloides* | - | Cidarinae |
| Cidaridae<br>(Goniocidarinae) | *Goniocidaris tubaria* | - | *Goniocidaris tubaria* | *Goniocidaris tubaria* | *Goniocidaris tubaria* |
| Cidaridae<br>(Phyllacanthina) | *Phyllacanthus* sp.<sup>a</sup> | - | *Phyllacanthus imperialis* | - | *Phyllacanthus* |
| Cidaridae<br>(Stereocidarinae) | *Stereocidaris sceptrifera* | - | *Stereocidaris sceptriferoides* | - | *Stereocidaris* |
| Cidaridae<br>(Stylocidarinae) | *Stylocidaris* sp.<sup>a</sup> | *Prionocidaris baculosa* | *Stylocidaris lineata* | - | Stylocidarinae |
| Cidaridae<br>(Typocidarinae) | *Typocidaris malum* | - | *Typocidaris malum* | *Typocidaris malum* | *Typocidaris malum* |
| Claviasteridae | *Claviaster libycus* | - | *Claviaster libycus* | *Claviaster libycus* | *Claviaster libycus* |
| Clypeasteridae | *Clypeaster rosaceus* | *Clypeaster rosaceus* | *Clypeaster rosaceus* | - | *Clypeaster rosaceus* |
| Clypeasteridae<br>(Ammotrophinae) | *Ammotrophus cyclius* | - | *Ammotrophus cyclius* | - | *Ammotrophus cyclius* |
| Clypeasteridae<br>(Arachnoidinae) | *Arachnoides placenta*<br>*Fellaster zelandiae* | - | *Arachnoides placenta* | *Fellaster zelandiae* | Arachnoidinae |
| Clypeidae | *Clypeus plotii* | - | *Clypeus plotii* | *Clypeus plotii* | *Clypeus plotii* |
| Clypeolampadidae | *Clypeolampas ovatus* | - | *Clypeolampas ovatus* | *Clypeolampas ovatus* | *Clypeolampas ovatus* |
| Coenholectypidae | *Coenholectypus macropygus* | - | *Coenholectypus macropygus* | *Coenholectypus macropygus* | *Coenholectypus macropygus* |
| Collyritidae | *Collyrites ellipticus* | - | *Collyrites ellipticus* | *Collyrites ellipticus* | *Collyrites ellipticus* |
| Conoclypidae | *Conoclypus conoideus* | - | *Conoclypus conoideus* | *Conoclypus conoideus* | *Conoclypus conoideus* |
| Conulidae | *Conulus albogalerus* | - | *Conulus albogalerus* | *Conulus albogalerus* | *Conulus albogalerus* |
| Corystusidae | *Corystus dysasteroides*<br>*Corystus relictus* | - | *Corystus dysasteroides* | *Corystus dysasteroides* | *Corystus* |
| Ctenocidaridae | *Ctenocidaris speciosa* | - | *Ctenocidaris speciosa* | - | *Ctenocidaris speciosa* |
| Dendrasteridae | *Dendraster excentricus* | *Dendraster excentricus* | *Dendraster excentricus* | - | *Dendraster excentricus* |
| Desorellidae | *Desorella elata* | - | *Desorella elata* | *Desorella elata* | *Desorella elata* |
| Diadematidae | *Diadema antillarum* | *Lissodiadema lorioli* | *Diadema savignyi* | - | Diadematidae |
| Diplocidaridae | *Diplocidaris gigantea* | - | *Diplocidaris gigantea* | *Diplocidaris gigantea* | *Diplocidaris gigantea* |
| Diplopodiidae | *Diplopodia pentagona* | - | *Diplopodia pentagona* | *Diplopodia pentagona* | *Diplopodia pentagona* |
| Disasteridae | *Disaster granulosus* | - | *Disaster granulosus* | *Disaster granulosus* | *Disaster granulosus* |
| Discoididae | *Discoides subuculus* | - | *Discoides subuculus* | *Discoides subuculus* | *Discoides subuculus* |
| Echinarachniidae | *Echinarachnius parma* | *Echinarachnius parma* | *Echinarachnius parma* | - | *Echinarachnius parma* |
| Echinidae | *Echinus esculentus* | - | *Echinus esculentus* | - | *Echinus esculentus* |
| Echinocorythidae | *Echinocorys scutata* | - | *Echinocorys scutata* | *Echinocorys scutata* | *Echinocorys scutata* |
| Echinolampadidae | *Echinolampas ovata* | *Conolampas sigsbei* | *Echinolampas depressa* | - | Echinolampadidae |
| Echinometridae | *Echinometra lucunter* | *Evechinus chloroticus* | *Echinometra mathaei* | - | Echinometridae |
| Echinoneidae | *Echinoneus cyclostomus* | - | *Echinoneus cyclostomus* | *Echinoneus cyclostomus* | *Echinoneus cyclostomus* |
| Echinothuriidae<br>(Echinothuriinae) | *Araeosoma fenestratum* | *Araeosoma leptaleum* | *Araeosoma fenestratum* | - | *Araeosoma* |
| Echinothuriidae<br>(Hygrosomatinae) | *Hygrosoma petersii* | - | *Hygrosoma petersii* | - | *Hygrosoma petersii* |
| Echinothuriidae<br>(Sperosomatinae) | *Sperosoma grimaldii* | - | *Sperosoma obscurum* | - | *Sperosoma* |
| Emiratiidae | *Emiratia raskhaimahensis* | - | *Emiratia raskhaimahensis* | *Emiratia raskhaimahensis* | *Emiratia raskhaimahensis* |
| Eoscutellidae | *Eoscutella coosensis* | - | *Eoscutella coosensis* | *Eoscutella coosensis* | *Eoscutella coosensis* |
| Eupatagidae | *Eupatagus valenciennesi*<br>*Eupatagus lymani* | - | *Eupatagus lymani* | - | *Eupatagus* |
| Eurypatagidae | *Eurypatagus ovalis* | - | *Eurypatagus ovalis* | - | *Eurypatagus ovalis* |
| Faujasidae | *Faujasia apicalis*<br>*Faujasia eccentricipora* | - | *Faujasia eccentricipora* | *Faujasia apicalis* | *Faujasia* |
| Faujasidae<br>(Stigmatopyginae) | *Stigmatopygus pulchellus* | - | *Stigmatopygus pulchellus* | *Stigmatopygus pulchellus* | *Stigmatopygus pulchellus* |
| Fibulariidae | *Echinocyamus pusillus* | *Echinocyamus crispus* | *Echinocyamus pusillus* | - | *Echinocyamus* |
| Fibulariidae | *Fibularia ovulum* | - | *Fibularia ovulum* | - | *Fibularia ovulum* |
| Fossulasteridae | *Fossulaster halli* | - | *Fossulaster halli* | *Fossulaster halli* | *Fossulaster halli* |
| Galeritidae | *Galerites vulgaris* | - | *Galerites vulgaris* | *Galerites vulgaris* | *Galerites vulgaris* |
| Galeropygidae | *Galeropygus sublaevis* | - | *Galeropygus sublaevis* | *Galeropygus sublaevis* | *Galeropygus sublaevis* |
| Glyphocyphidae | *Glyphocyphus radiatus* | - | *Glyphocyphus radiatus* | *Glyphocyphus radiatus* | *Glyphocyphus radiatus* |
| Glypticidae | *Glypticus hieroglyphicus* | - | *Glypticus hieroglyphicus* | *Glypticus hieroglyphicus* | *Glypticus hieroglyphicus* |
| Glyptocidaridae | *Glyptocidaris crenularis* | - | *Glyptocidaris crenularis* | *Glyptocidaris crenularis* | *Glyptocidaris crenularis* |
| Goniophoridae | *Goniophorus lunulatus* | - | *Goniophorus lunulatus* | *Goniophorus lunulatus* | *Goniophorus lunulatus* |
| Hemiasteridae | *Hemiaster bufo* | - | *Hemiaster bufo* | *Hemiaster bufo* | *Hemiaster bufo* |
| MONGIARDINO KOCH & THOMPSON |  |  |  |  |  |
| Hemicidaridae | *Hemicidaris intermedia* | - | *Hemicidaris intermedia* | *Hemicidaris intermedia* | *Hemicidaris intermedia* |
| Hemipneustidae | *Hemipneustes striatoradiatus* | - | *Hemipneustes striatoradiatus* | *Hemipneustes striatoradiatus* | *Hemipneustes striatoradiatus* |
| Heterodiademataidae | *Heterodiadema lybica* | - | *Heterodiadema lybica* | *Heterodiadema lybica* | *Heterodiadema lybica* |
| Histocidaridae | *Histocidaris elegans* | - | *Histocidaris elegans* | - | *Histocidaris elegans* |
| Histocidaridae | *Histocidaris purpurata* | - | *Histocidaris purpurata* | - | *Histocidaris purpurata* |
| Holasteridae | *Holaster nodulosus* | - | *Holaster nodulosus* | *Holaster nodulosus* | *Holaster nodulosus* |
| Holactypidae | *Holactypus depressus* | - | *Holactypus depressus* | *Holactypus depressus* | *Holactypus depressus* |
| Hyboclypids | *Hyboclypus gibberulus*<br>*Hyboclypus sandalinus* | - | *Hyboclypus gibberulus* | *Hyboclypus gibberulus* | *Hyboclypus* |
| Hyposalenidae | *Hyposalenia stellulata* | - | *Hyposalenia stellulata* | *Hyposalenia stellulata* | *Hyposalenia stellulata* |
| Kamptosomatidae | *Kamptosoma asterias* | - | *Kamptosoma asterias* | - | *Kamptosoma asterias* |
| Laganidae | *Laganum laganum* | - | *Laganum decagonale* | - | *Laganum* |
| Laganidae<br>(Neolaganinae) | *Neolaganum durhami*<br>*Neolaganum archerensis* | - | *Neolaganum durhami* | *Neolaganum durhami* | *Neolaganum* |
| Loveniidae | *Breynia australasiae* | - | *Breynia australasiae* | - | *Breynia australasiae* |
| Loveniidae | *Lovenia elongata* | - | *Lovenia elongata* | - | *Lovenia elongata* |
| Loveniidae<br>(Echinocardiinae) | *Echinocardium cordatum* | *Echinocardium mediterraneum* | *Echinocardium cordatum* | - | *Echinocardium* |
| Macropneustidae | *Macropneustes deshayesi*<br>*Macropneustes mortoni* | - | *Macropneustes mortoni* | *Macropneustes mortoni* | *Macropneustes* |
| Maretiidae | *Maretia planulata* | - | *Maretia planulata* | *Maretia planulata* | *Maretia planulata* |
| Megapneustidae | *Megapneustes grandis* | - | *Megapneustes grandis* | *Megapneustes grandis* | *Megapneustes grandis* |
| Mellitidae | *Mellita quinquesperforata* | *Mellita tenuis* | *Mellita quinquesperforata* | - | *Mellita* |
| Micrasteridae | *Micraster coranguinum* | - | *Micraster coranguinum* | *Micraster coranguinum* | *Micraster coranguinum* |
| Micrasteridae<br>(Cyclasterinae) | *Cyclaster regalis* | - | *Cyclaster regalis* | - | *Cyclaster regalis* |
| Micropygidae | *Micropyga tuberculata* | - | *Micropyga tuberculata* | - | *Micropyga tuberculata* |
| Miocidaridae | *Eotiaris verneuilliana* | - | - | *Eotiaris verneuilliana* | *Eotiaris verneuilliana* |
| Monophorasteridae | *Monophoraster darwini* | - | *Monophoraster darwini* | *Monophoraster darwini* | *Monophoraster darwini* |
| Neoglobatoridae | *Adelopneustes montainvillensis* | - | *Adelopneustes montainvillensis* | *Adelopneustes montainvillensis* | *Adelopneustes montainvillensis* |
| Neolampadidae | *Neolampas rostellata* | - | *Neolampas rostellata* | - | *Neolampas rostellata* |
| Nucleolitidae | *Nucleolites scutatus* | - | *Nucleolites scutatus* | *Nucleolites scutatus* | *Nucleolites scutatus* |
| Oligopygidae | *Haimea ruttini* | - | *Haimea ruttini* | *Haimea ruttini* | *Haimea ruttini* |
| Oligopygidae | *Oligopygus wetherbyi*<br>*Oligopygus haldemai* | - | *Oligopygus haldemani* | *Oligopygus haldemani* | *Oligopygus* |
| Orthopsidae | *Orthopsis miliaris* | - | *Orthopsis miliaris* | *Orthopsis miliaris* | *Orthopsis miliaris* |
| Ovulasteridae | *Ovulaster gauthieri*<br>*Ovulaster protodecimae* | - | *Ovulaster protodecimae* | *Ovulaster protodecimae* | *Ovulaster* |
| Palaeostomatidae | *Palaeostoma mirabile* | - | *Palaeostoma mirabile* | - | *Palaeostoma mirabile* |
| Palaeotropidae | *Palaeotropus josephinae* | - | *Palaeotropus josephinae* | - | *Palaeotropus josephinae* |
| Paleopneustidae | *Paleopneustes cristatus* | - | *Paleopneustes cristatus* | - | *Paleopneustes cristatus* |
| Parasalenidae | *Parasalenia gratiosa* | - | *Parasalenia gratiosa* | - | *Parasalenia gratiosa* |
| Parechinidae | *Parechinus angulosa* | *Paracentrotus lividus* | *Paracentrotus lividus* | - | Parechinidae |
| Pedinidae | *Caenopedina cubensis* | *Caenopedina hawaiiensis* | *Caenopedina cubensis* | - | *Caenopedina* |
| Pelanechinidae | *Pelanodiadema oolithicum*<br>*Pelanechinus corallina* | - | *Pelanechinus corallina* | *Pelanechinus corallina* | Pelanechinidae |
| Periasteridae | *Periaster elatus* | - | *Periaster elatus* | *Periaster elatus* | *Periaster elatus* |
| Pericosmidae | *Pericosmus latus*<br>*Pericosmus cordatus* | - | *Pericosmus latus* | *Pericosmus latus* | *Pericosmus* |
| Phormosomatidae | *Phormosoma placenta* | - | *Phormosoma placenta* | - | *Phormosoma placenta* |
| Phormosomatidae<br>(Paraphormosomatinae) | *Paraphormosoma alternans* | - | *Paraphormosoma alternans* | - | *Paraphormosoma alternans* |
| Phymosomatidae | *Gauthieria radiata* | - | *Gauthieria radiata* | *Gauthieria radiata* | *Gauthieria radiata* |
| Phymosomatidae | *Phymosoma koenigi* | - | *Phymosoma abbatei* | *Phymosoma koenigi* | *Phymosoma* |
| Plesiasteridae | *Plesiaster peini* | - | *Plesiaster peini* | *Plesiaster peini* | *Plesiaster peini* |
| Plesiolampadidae | *Plesiolampas placenta* | - | *Plesiolampas placenta* | *Plesiolampas placenta* | *Plesiolampas placenta* |
| Plexechinidae | *Plexechinus cinctus*<br>*Antrechinus nordenskjoldi* | - | *Plexechinus cinctus* | - | Plexechinidae |
| Pliolampadidae | *Pliolampas vassalli* | - | *Pliolampas vassalli* | *Pliolampas vassalli* | *Pliolampas vassalli* |
| Polycidaridae | *Polycidaris legayi*<br>*Polycidaris spinosa* | - | *Polycidaris suevica* | *Polycidaris suevica* | *Polycidaris* |
| Polydiaematidae | *Polydiadema mamillanum* | - | *Polydiadema mamillanum* | *Polydiadema mamillanum* | *Polydiadema mamillanum* |
| Pourtalesidae | *Pourtalesia jeffreysi* | - | *Pourtalesia jeffreysi* | - | *Pourtalesia jeffreysi* |
| Prenasteridae | *Prenaster alpinus* | - | *Prenaster alpinus* | *Prenaster alpinus* | *Prenaster alpinus* |
| Protoscutellidae | *Protoscutella plana* | - | *Protoscutella mississippiensis* | *Protoscutella plana* | *Protoscutella* |
| Pseudholasteridae | *Pseudholaster bicarinatus* | - | *Pseudholaster latissimus* | *Pseudholaster bicarinatus* | *Pseudholaster* |
| Pseudodiademmatidae | *Pseudodiadema pseudodiadema* | - | *Pseudodiadema pseudodiadema* | *Pseudodiadema pseudodiadema* | *Pseudodiadema pseudodiadema* |
| MONGIARDINO KOCH & THOMPSON |  |  |  |  |  |
| Pseudosaleniidae | *Pseudosalenia aspera* | - | *Pseudosalenia aspera* | *Pseudosalenia aspera* | *Pseudosalenia aspera* |
| Psychocidaridae | *Roseicidaris morieri* | - | *Roseicidaris rebouli* | *Roseicidaris morieri* | *Roseicidaris* |
| Psychocidaridae | *Tylocidaris ohshimai* | - | *Tylocidaris ohshimai* | - | *Tylocidaris ohshimai* |
| Pygasteridae | *Pygaster semisulcatus* | - | *Pygaster semisulcatus* | *Pygaster semisulcatus* | *Pygaster semisulcatus* |
| Pygaulidae | *Pygaulus desmoulinsii* | - | *Pygaulus desmoulinsii* | *Pygaulus desmoulinsii* | *Pygaulus desmoulinsii* |
| Pygorhytidae | *Pygorhytis ringens* | - | *Pygorhytis ringens* | *Pygorhytis ringens* | *Pygorhytis ringens* |
| Rhabdocidaridae | *Rhabdocidaris orbignyana* | - | *Rhabdocidaris orbignyana* | *Rhabdocidaris orbignyana* | *Rhabdocidaris orbignyana* |
| Rotulidae | *Rotula deciesdigitatus* | - | *Rotula deciesdigitatus* | - | *Rotula deciesdigitatus* |
| Saleniidae | *Salenia petalifera* | - | *Salenia petalifera* | *Salenia petalifera* | *Salenia petalifera* |
| Saleniidae<br>(Holosaleniini) | *Holosalenia batnensis* | - | *Holosalenia batnensis* | *Holosalenia batnensis* | *Holosalenia batnensis* |
| Saleniidae<br>(Salenocidarini) | *Salenocidaris varispina* | - | *Salenocidaris varispina* | - | *Salenocidaris varispina* |
| Schizasteridae<sup>b</sup> | *Brisaster fragilis* | - | *Brisaster fragilis* | - | *Brisaster fragilis* |
| Schizasteridae<sup>b</sup> | *Ova canalifera* | *Abatus agassizii* | *Abatus cordatus* | - | Schizasteridae |
| Scutasteridae | *Scutaster andersoni* | - | *Scutaster andersoni* | *Scutaster andersoni* | *Scutaster andersoni* |
| Scutellidae | *Scutella subrotunda* | - | *Scutella subrotunda* | *Scutella subrotunda* | *Scutella subrotunda* |
| Scutellinidae | *Scutellina lenticularis* | - | *Scutellina lenticularis* | *Scutellina lenticularis* | *Scutellina lenticularis* |
| Scutellinoididae | *Scutellinoides patella* | - | *Scutellinoides patella* | *Scutellinoides patella* | *Scutellinoides patella* |
| Serpianotiaridae | *Serpianotiaris coeava* | - | *Serpianotiaris coeava* | *Serpianotiaris coeava* | *Serpianotiaris coeava* |
| Somaliasteridae | *Somaliaster magniventer* | - | *Somaliaster magniventer* | *Somaliaster magniventer* | *Somaliaster magniventer* |
| Spatangidae | *Spatangus purpureus* | - | *Spatangus purpureus* | *Spatangus purpureus* | *Spatangus purpureus* |
| Stegasteridae | *Stegaster gillieronii* | - | *Stegaster gillieronii* | *Stegaster gillieronii* | *Stegaster gillieronii* |
| Stenonasteridae | *Stenonaster tuberculata* | - | *Stenonaster tuberculata* | *Stenonaster tuberculata* | *Stenonaster tuberculata* |
| Stomechinidae | *Stomechinus bigranularis* | - | *Stomechinus bigranularis* | *Stomechinus bigranularis* | *Stomechinus bigranularis* |
| Stomopneustidae | *Stomopneustes variolaris* | *Stomopneustes variolaris* | *Stomopneustes variolaris* | - | *Stomopneustes variolaris* |
| Strongylocentrotidae | *Strongylocentrotus droebachiensis* | *Strongylocentrotus purpuratus* | *Strongylocentrotus droebachiensis* | - | *Strongylocentrotus* |
| Taiwanasteridae | *Sinaechinocyamus mai* | - | *Sinaechinocyamus mai* | - | *Sinaechinocyamus mai* |
| Temnopleuridae | *Temnopleurus toreumaticus* | - | *Temnopleurus toreumaticus* | - | *Temnopleurus toreumaticus* |
| Tithoniidae | *Tithonia convexa* | - | *Tithonia convexa* | *Tithonia convexa* | *Tithonia convexa* |
| Toxasteridae | *Toxaster retusus* | - | *Toxaster retusus* | *Toxaster retusus* | *Toxaster retusus* |
| Toxopneustidae | *Toxopneustes pileolus* | *Lythechinus variegatus* | *Toxopneustes pileolus* | - | Toxopneustidae |
| Triadotiaridae | *Triadotiaris grandaevus* | - | *Triadotiaris grandaevus* | *Triadotiaris grandaevus* | *Triadotiaris grandaevus* |
| Trigonocidaridae | *Trigonocidaris albida* | - | *Trigonocidaris albida* | - | *Trigonocidaris albida* |
| Unifasciidae | *Unifascia carolinensis* | - | *Unifascia carolinensis* | *Unifascia carolinensis* | *Unifascia carolinensis* |
| Urechinidae | *Urechinus naresianus* | *Cystechinus giganteus*.<sup>c</sup> | *Urechinus naresianus* | - | Urechinidae |
| Zeuglopleuridae | *Zeuglopleurus costulatus* | - | *Zeuglopleurus costulatus* | *Zeuglopleurus costulatus* | *Zeuglopleurus costulatus* |
| Echinoidea incertae sedis | *Glyptodidema granulatatum* | - | *Glyptodidema granulatatum* | *Glyptodidema granulatatum* | *Glyptodidema granulatatum* |
| Stem Echinoidea<br>(Archaeocidaridae) | *Archaeocidaris whatleyensis*<br>*Archaeocidaris. brownwoodensis* | - | - | *Archaeocidaris whatleyensis* | *Archaeocidaris* |
| Holothuroidea<br>(Synallactida) | - | *Apostichopus parvimensis* | - | - | - |
| Holothuroidea<br>(Holothuriida) | - | *Holothuria forskali* | - | - | - |
| Holothuroidea (Apodida) | - | *Leptosynapta clarki* | - | - | - |

<sup>a</sup> Kroh and Smith (2010) did not specify which species were used for coding.
<sup>b</sup> Given lack of knowledge of relationships within schizasterid spatangoids, the molecular data for *Abatus* could have been merged with the morphological data of either of the two representatives of the family. This decision however cannot have any effect on topology, and merging was thus performed at random.
<sup>c</sup> This species was identified as *Pilematechinus* sp. in Mongiardino Koch et al. (2018) but has since been identified as *Cystechinus* c.f. *giganteus* by R. Mooi.

**Table S2:**
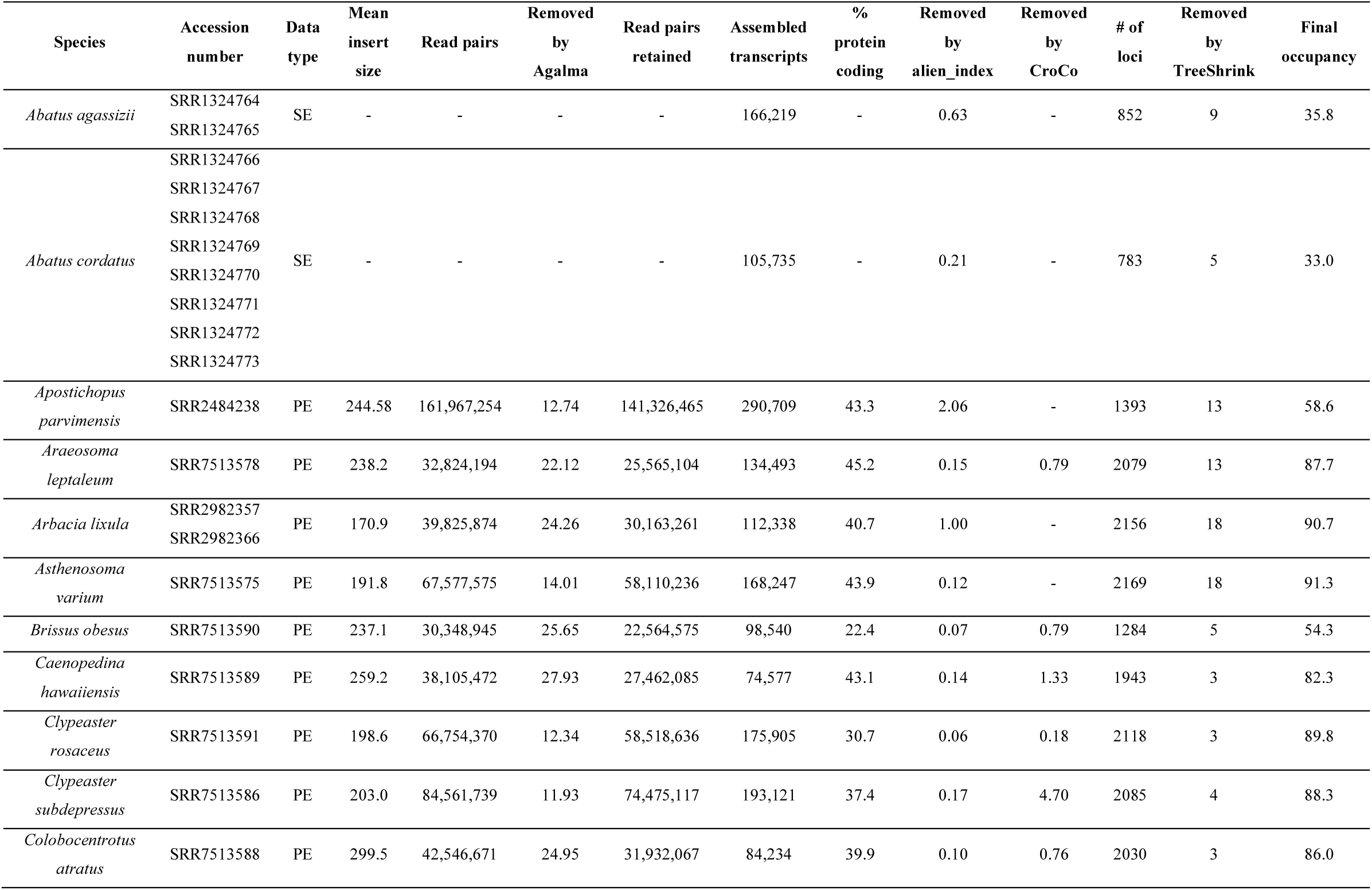
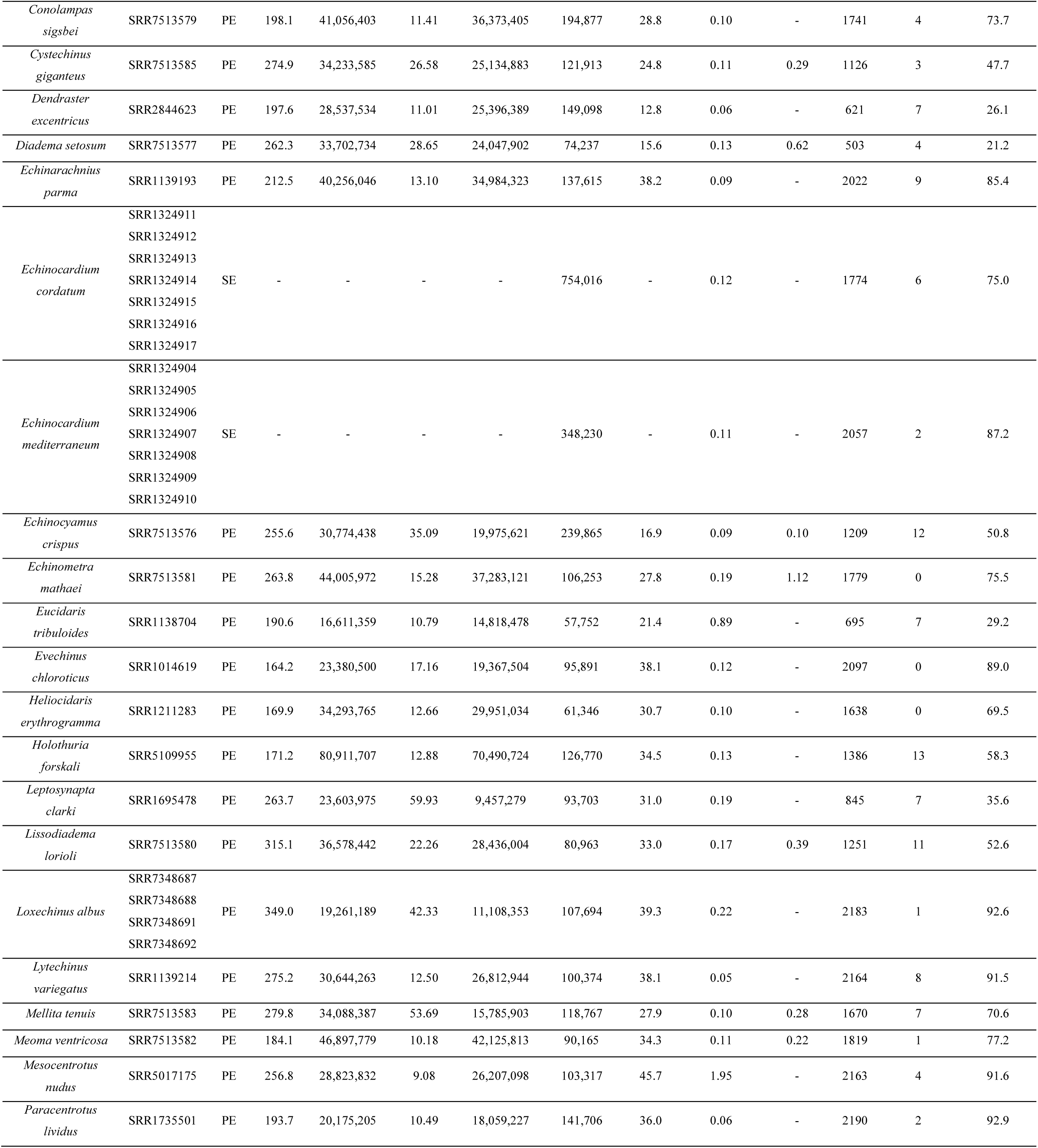
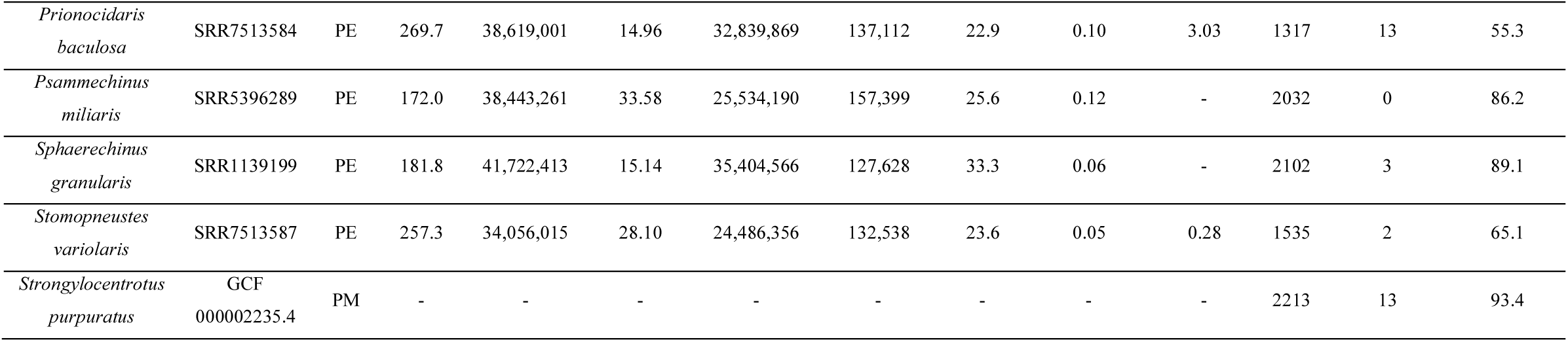
Genomic resources used to build the phylogenomic dataset and statistics of the bioinformatic pipeline used for assembly, curation and orthology inference. Data types are single-end transcriptomes (SE), pair-end transcriptomes (PE) and protein models derived from genomes (PM). Mean insert size is expressed in number of base pairs. Read pairs shows the number of read pairs in each dataset after processing with Trimmomatic. Those further removed by the curation steps taken by Agalma are shown as percentages, resulting in the final number of read pairs retained. These were then used for transcriptome assembly. The assemblies were further sanitized with alien_index and CroCo, and the transcripts removed by both of these are shown as a percentage of the original transcriptome size (see also Fig. S1 of SI File 4). Number of loci shows the occupancy of terminals in the supermatrix generated by Agalma (2,356 loci at 70% occupancy), after which loci were further removed by TreeShrink resulting in the final occupancy (also shown in Fig S3 of Si File 4).

**Table S3:**
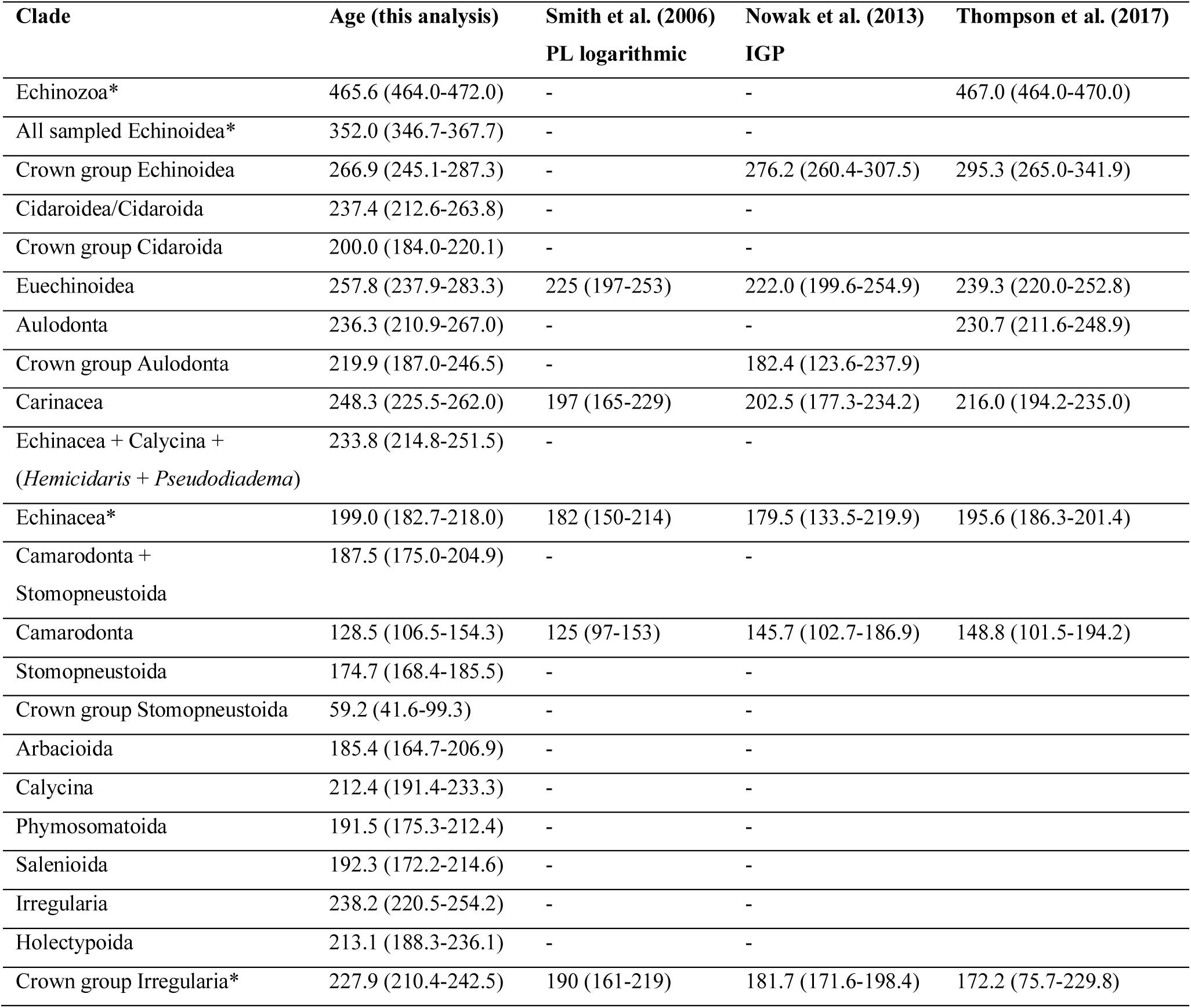
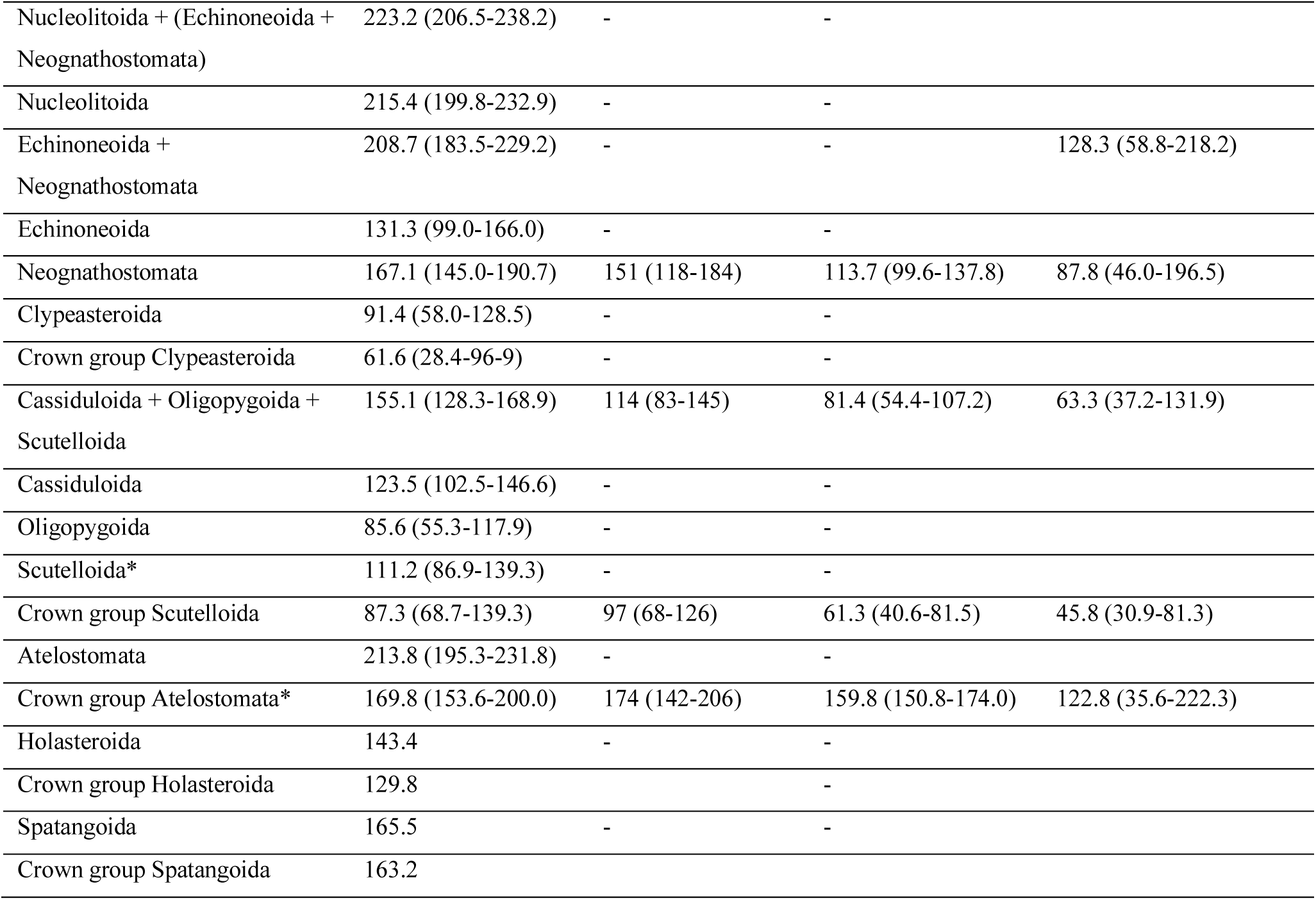
Inferred dates of divergence of major clades in our total-evidence dated analysis. Only clades ranked at or above the level of order are shown. Clades are defined based on the topologies of Figure 5 and Figure S13 (SI File 4), as well as the taxonomic changes proposed in the main text. If a clade is missing from the majority-rule consensus, the age of the mcc tree is shown. Age is expressed in Ma and includes the 95% highest posterior density. These dates should be taken with caution as many are constrained exclusively based on a morphological clock. For comparison, we have included the results of three other time-calibration studies. If those authors used multiple calibration approaches, their preferred method is reported. PL logarithmic = penalized likelihood method with logarithmic-penalty function. Clades with an asterisk were constrained for node dating. IGP = Informative gamma priors.

## SI File 4: Supplementary Figures

**Figure S1:**
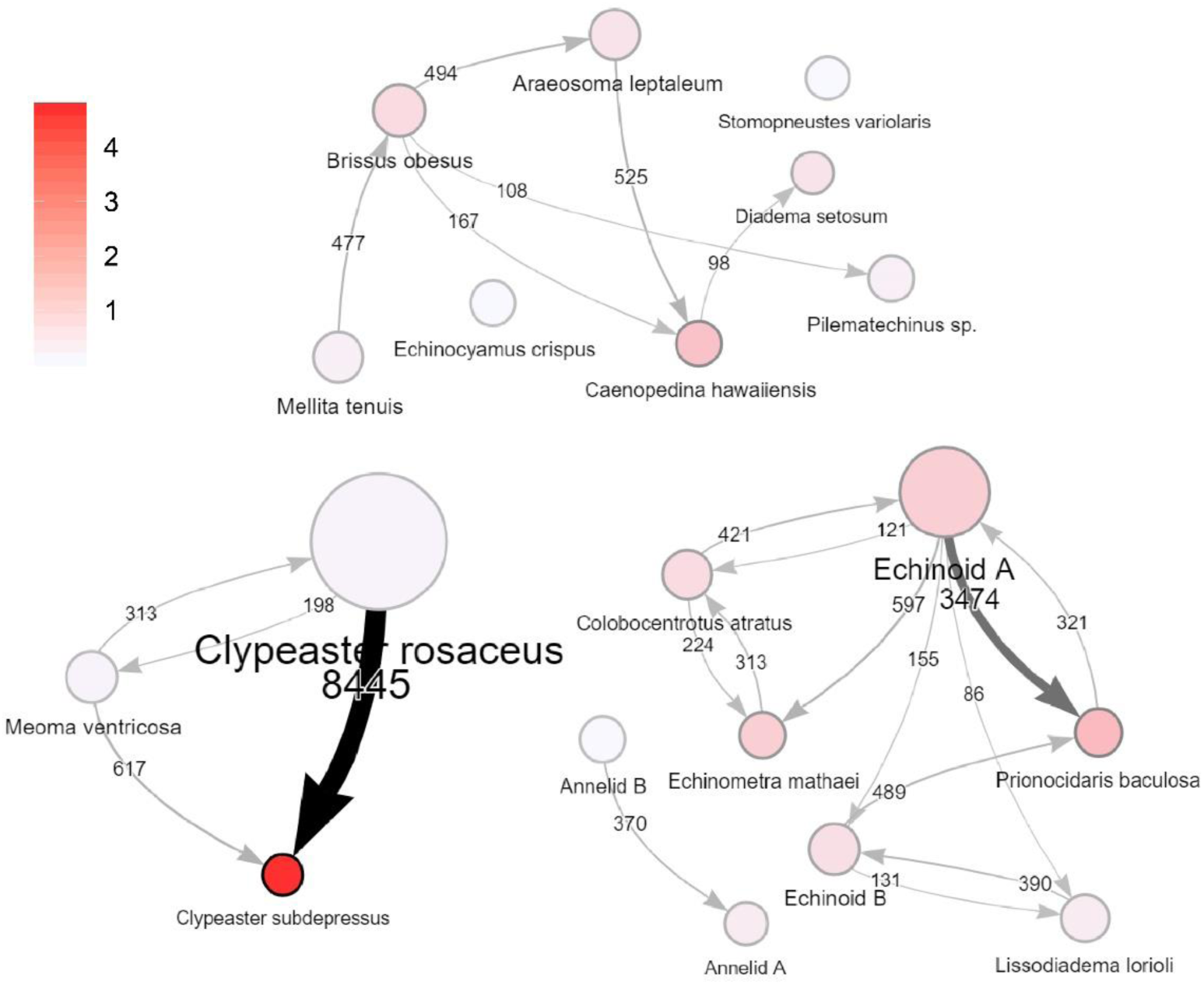
Results of cross-contamination detected by Croco. The three networks constitute three independent multiplexed Illumina runs for which we had access to all taxa sequenced simultaneously, nodes and links represent transcriptomes and cross-contaminations. Node size is proportional to the number of times the transcriptome was found to be the source of contamination, and node color the percentage of contaminated transcripts (see scale). Links shown represent event of contamination leading to at least 1/1000 contaminated transcripts in the target transcriptome. Evidence for contamination is relatively minimal, with an average of 0.99% of transcripts identified as cross-contaminants (see Table S1 of SI File 2). The largest event detected is between two closely related species of *Clypeaster* and should be taken with caution.

**Figure S2:**
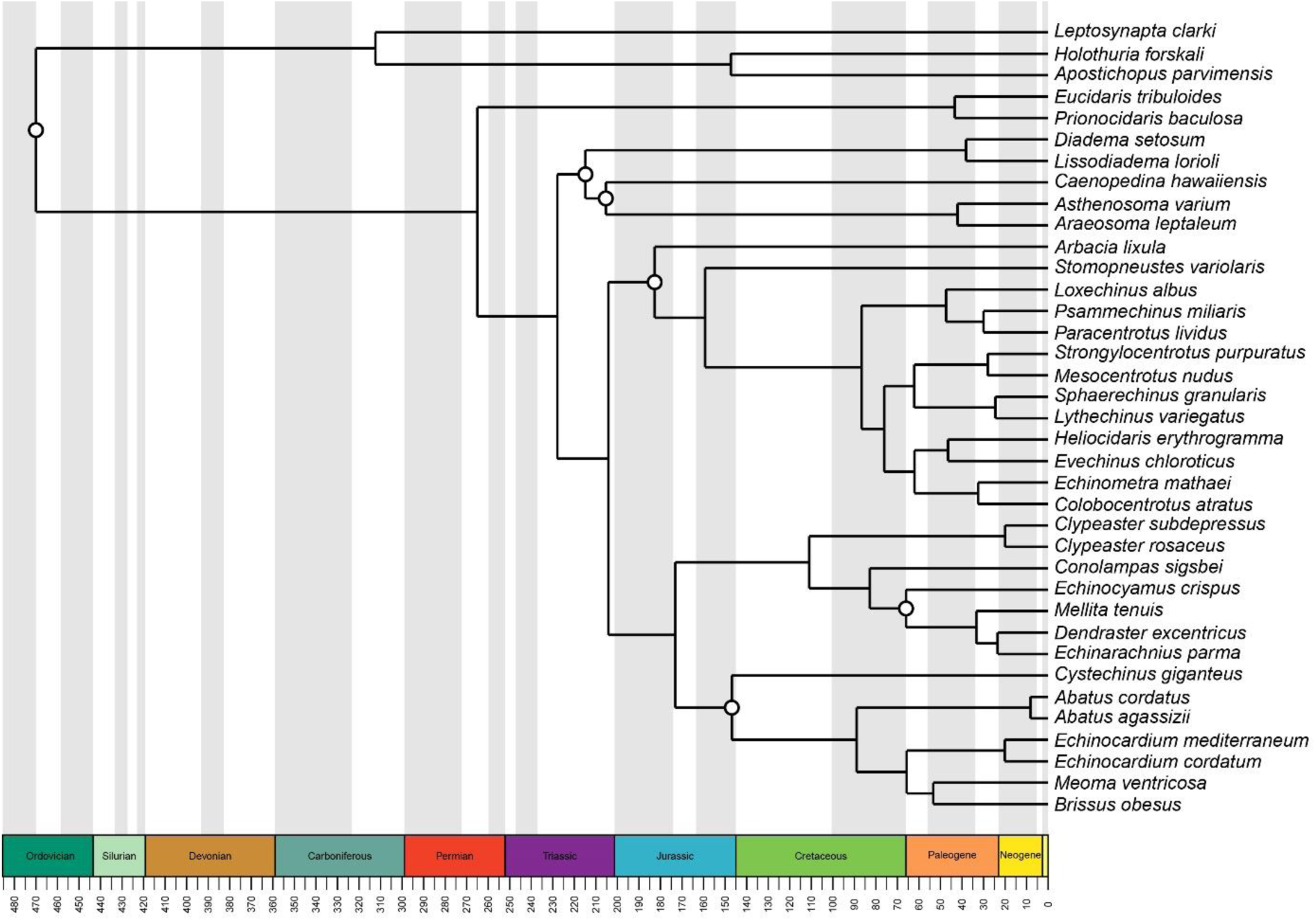
Time-calibrated molecular phylogeny obtained using penalized likelihood. The chronogram was obtained by calibrating the partitioned maximum likelihood topology with the correlated model of substitution rate variation. The six node calibrations employed are represented with circles, and described in detail in SI File 3. The tree was used exclusively to estimate the rate of evolution of positions in the supermatrix.

**Figure S3:**
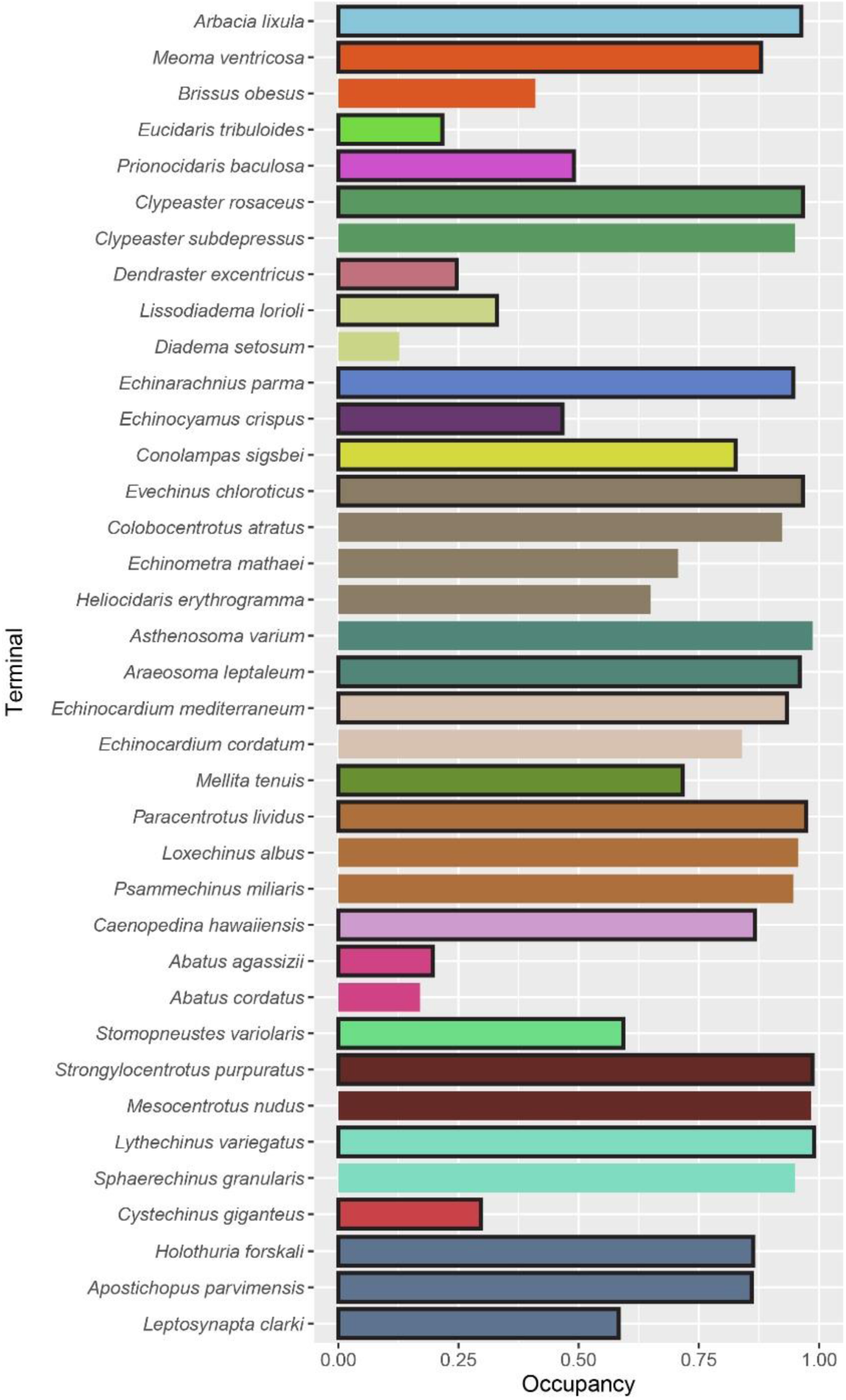
Level of occupancy of terminals in the 300-loci matrix obtained after subsampling. Colors show the different lineages represented in the morphological dataset. Terminals highlighted with a black border were chosen to merge with morphological data and build the total-evidence dataset (see also Table S1 of SI File 2). If more than one terminal per lineage was available, we selected either the one most closely related to that sampled for morphology, or the one with the highest occupancy.

**Figure S4:**
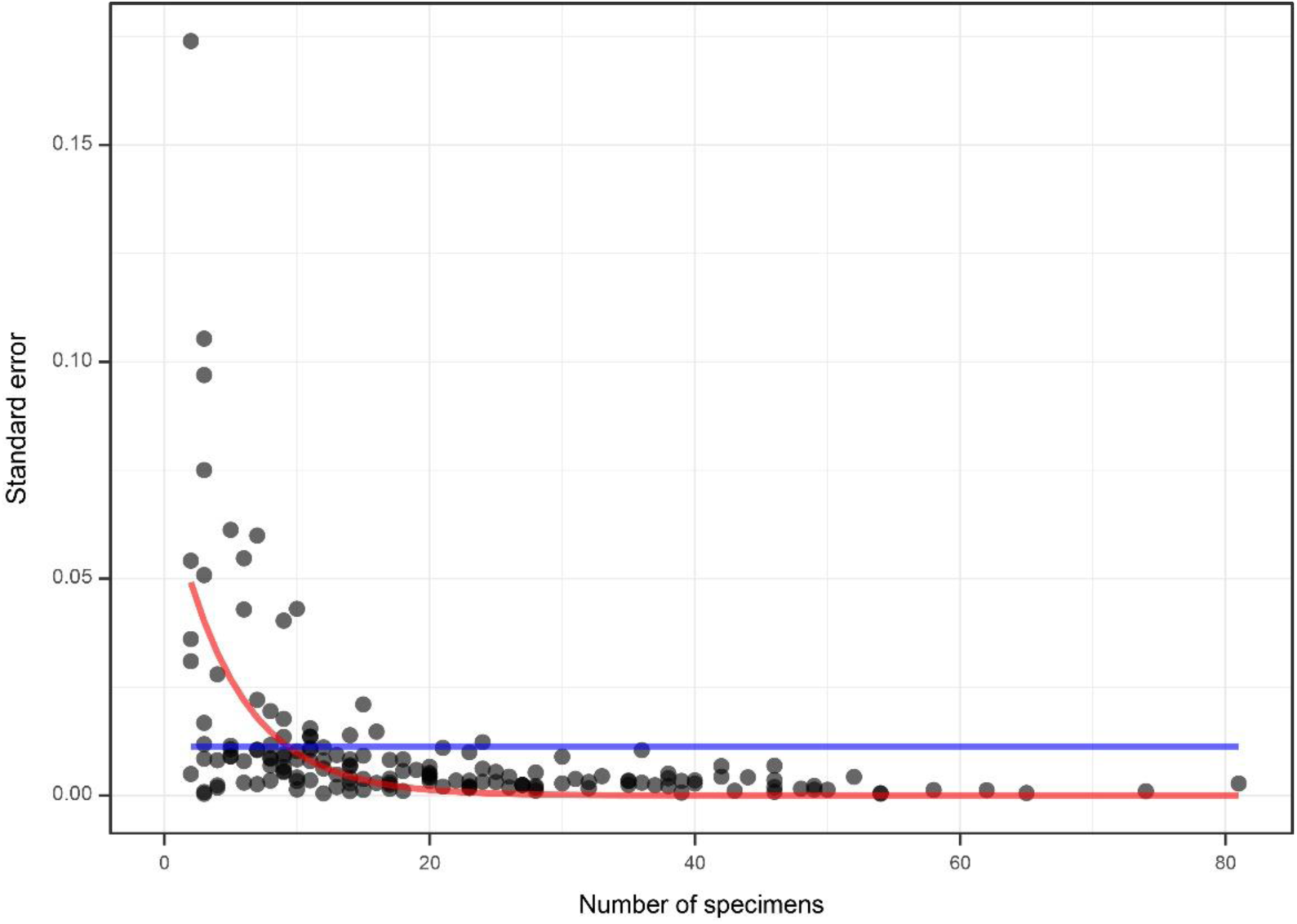
Approach to estimate body size measurement error. For 44 of the terminals sampled (27.8%), our dataset included observations of body sizes taken from the primary literature that averaged data across multiple individuals, sometimes even hundreds (see SI File 9 for further detail). Estimation of the standard errors of the mean for these species would require taking these observations as coming from one individual, inflating standard errors and increasing uncertainty downstream. Instead, an exponential decay function (red) was fitted to the standard error of species for which all observations come from individual specimens. This function was used to predict the standard error of the 44 taxa with individual observations derived from multiple individuals. A common alternative approach, replacing these by the mean standard error (blue), does not account for the reduced uncertainty obtained with increasing the number of measurements, and can therefore either underestimate or overestimate uncertainty. We also believe our approach provides a meaningful constraint on the expected error of terminals with a single observation, which cannot be estimated otherwise.

**Figure S5:**
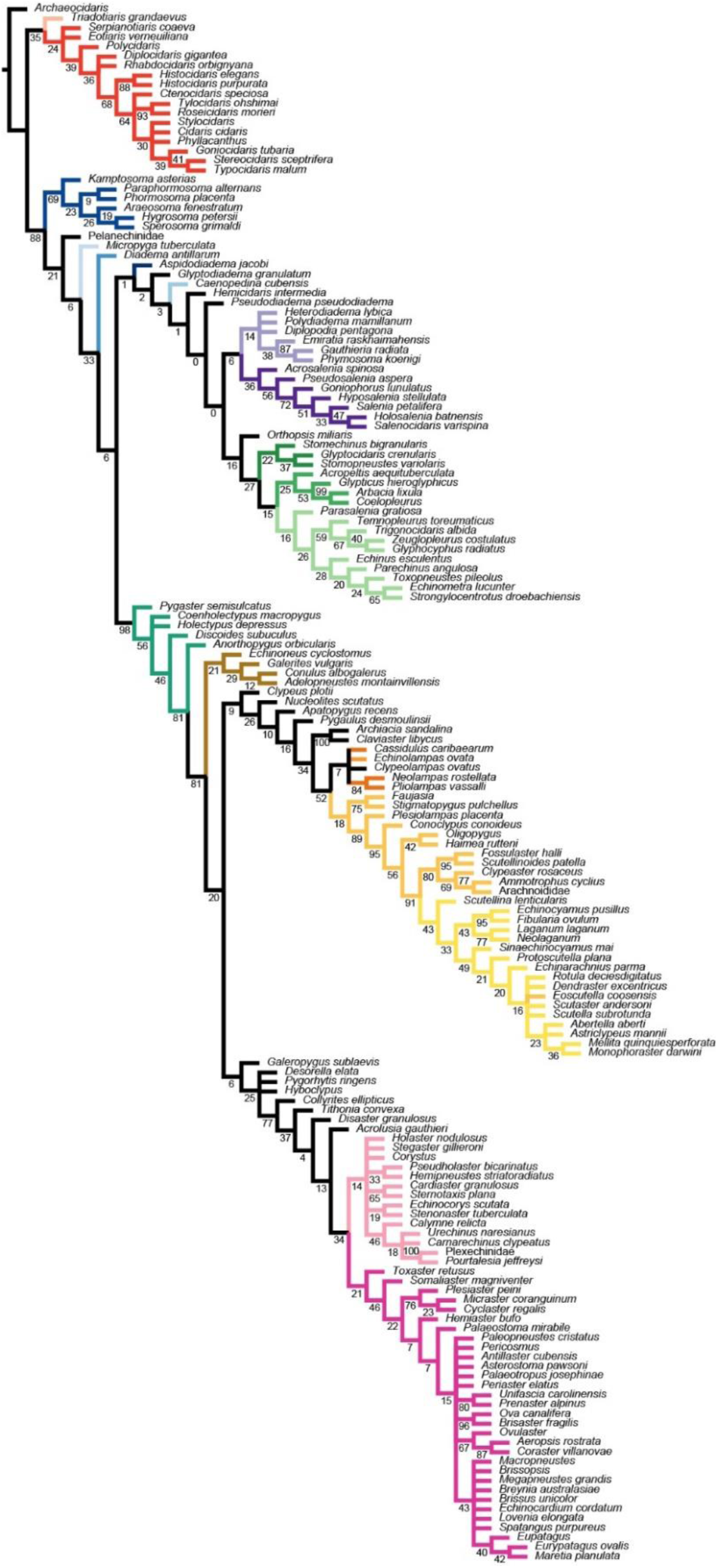
Strict consensus of the parsimony analysis of morphology under equal weights. Colors are as in Figure 2, numbers on nodes represent jackknife frequencies.

**Figure S6:**
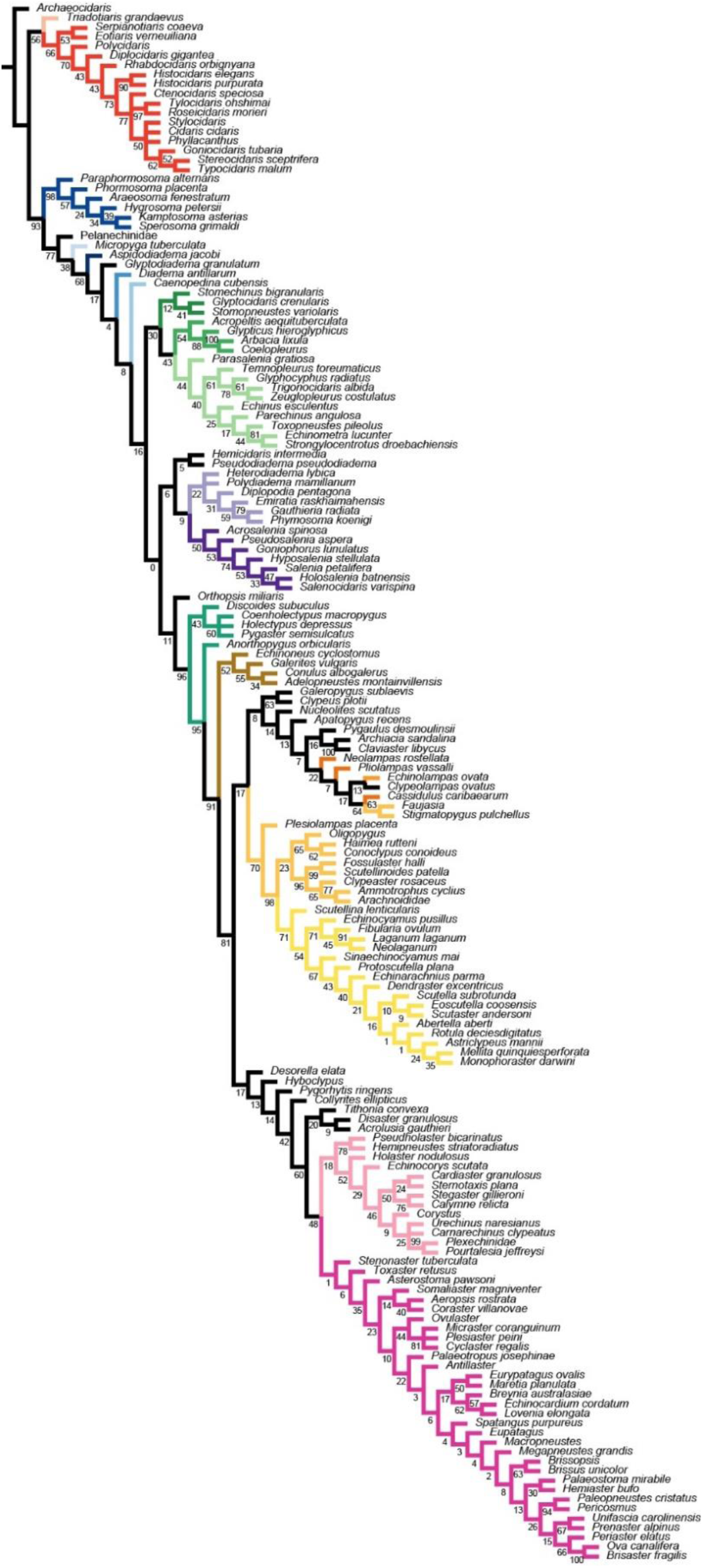
Strict consensus of the parsimony analysis of morphology under implied weighting (k = 6). Colors are as in Figure 2, numbers on nodes represent jackknife frequencies.

**Figure S7:**
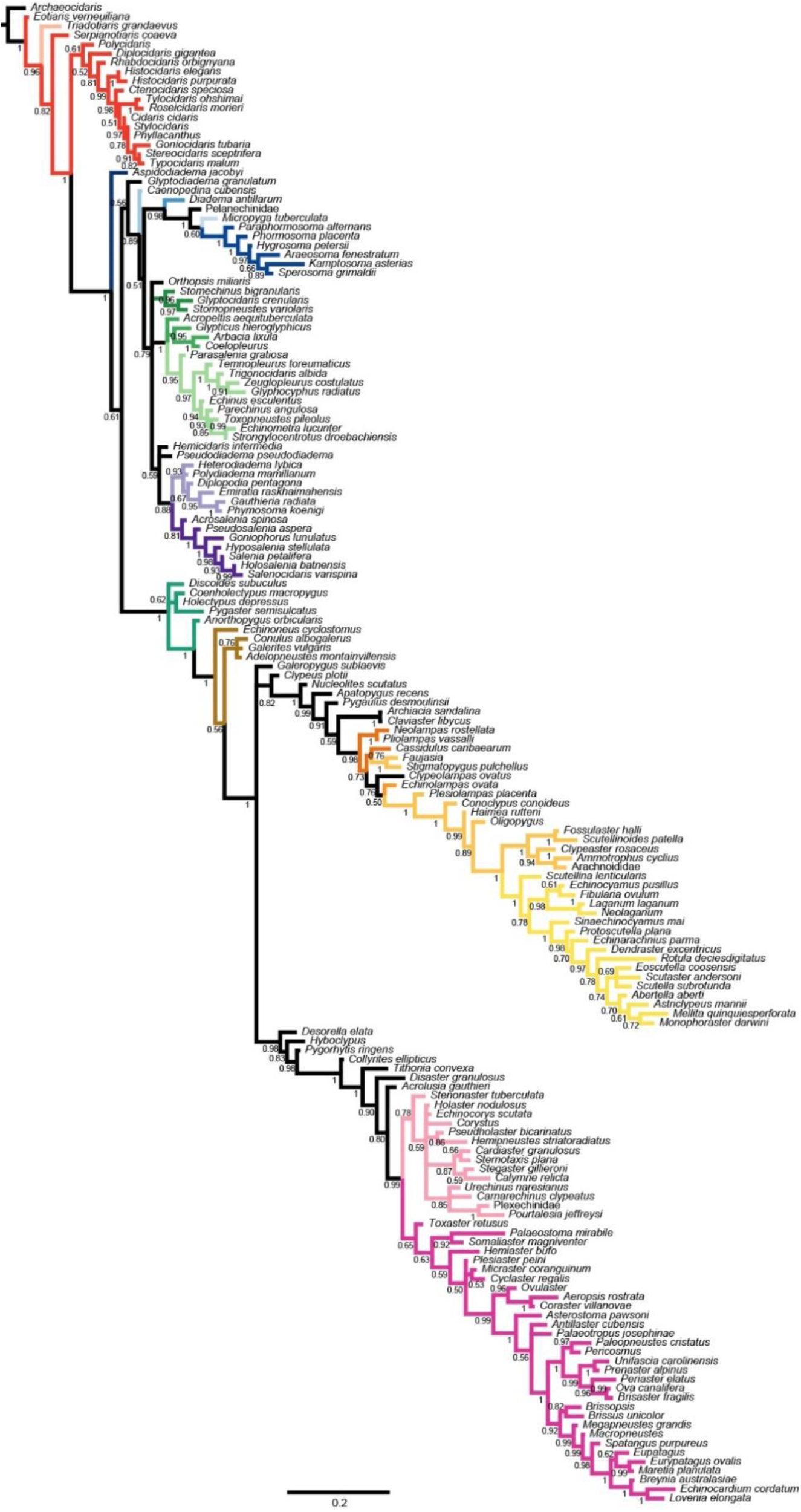
Majority-rule consensus of the uncalibrated Bayesian analysis of morphology. Colors are as in Figure 2, numbers on nodes represent posterior probabilities.

**Figure S8:**
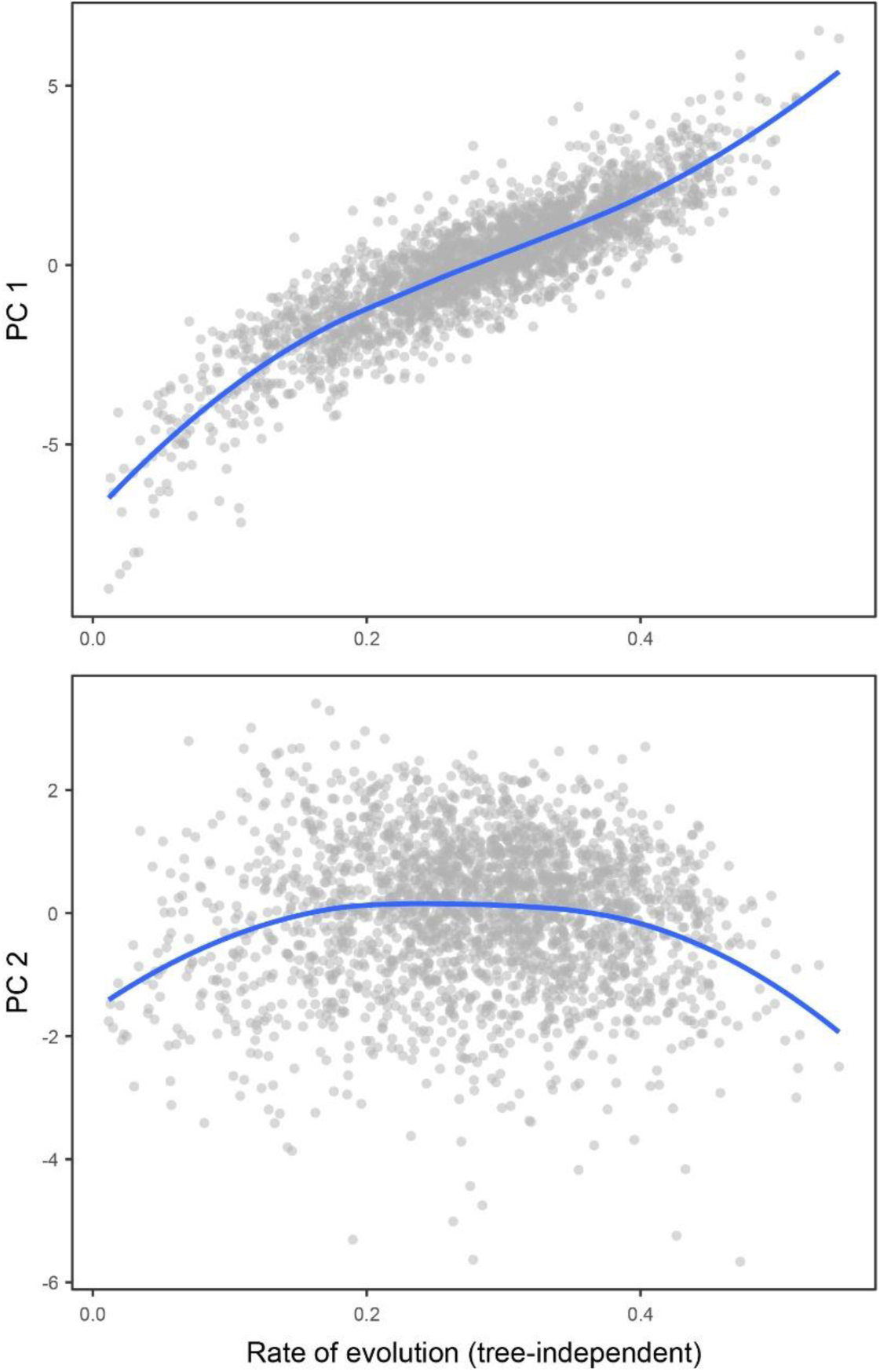
Correlation between rate of evolution and the first two axes obtained from the principal component analysis of the gene property dataset. The figure replicates the results shown in Figure 3c using a tree-independent method to estimate rate of evolution. Each dot represents a locus, and the blue lines correspond to loess regressions. Axis 1 shows a strong linear relationship with the rate of evolution (linear regression: R^2^ = 0.76, *P* < 10^-16^).

**Figure S9:**
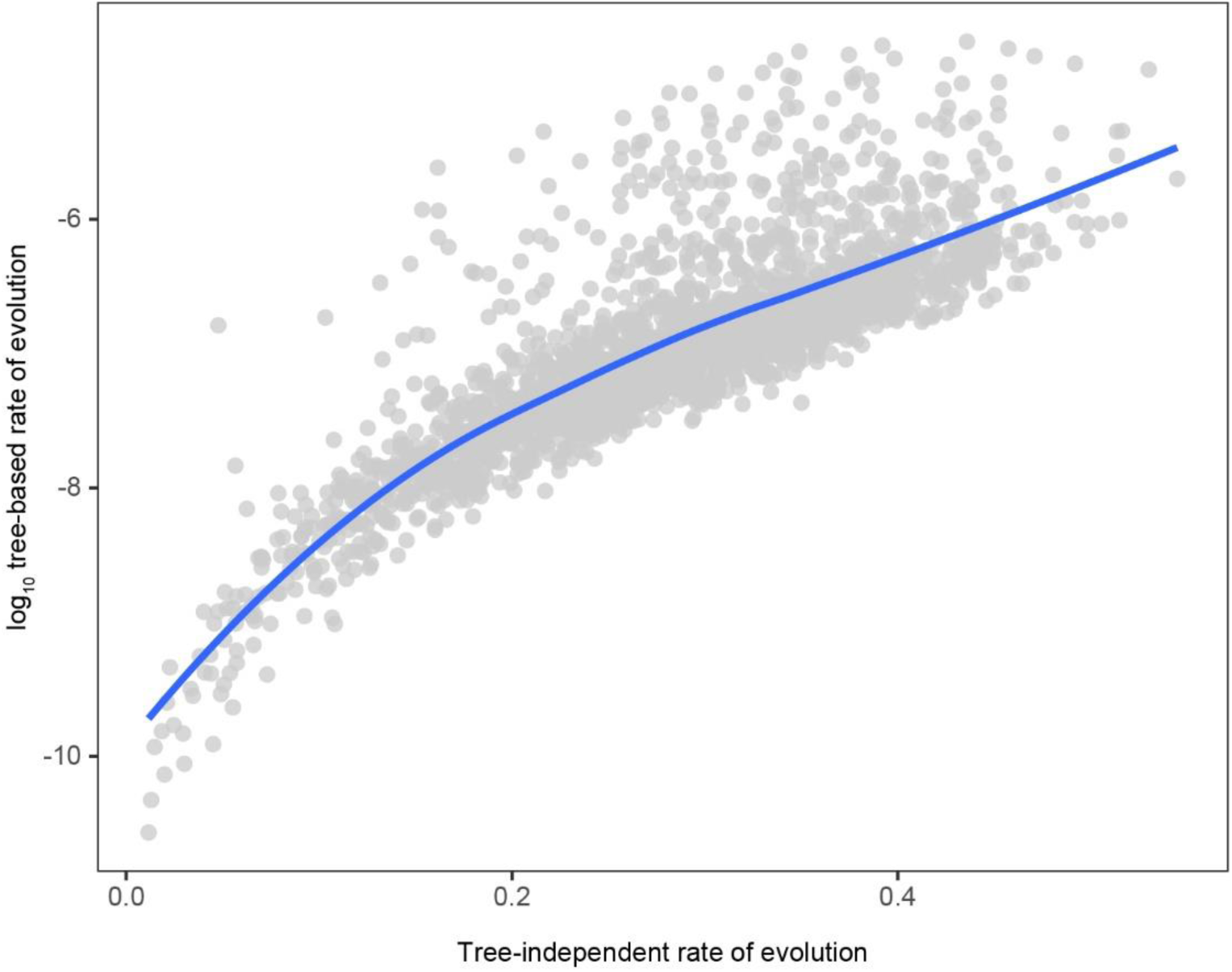
Correlation between tree-based and tree-independent estimates of evolutionary rate. Each dot represents a locus, and the blue lines correspond to loess regressions. There is a very strong log-linear relationship between both approaches to estimating evolutionary rates (Person’s ρ = 0.82, *P* < 10^-16^).

**Figure S10:**
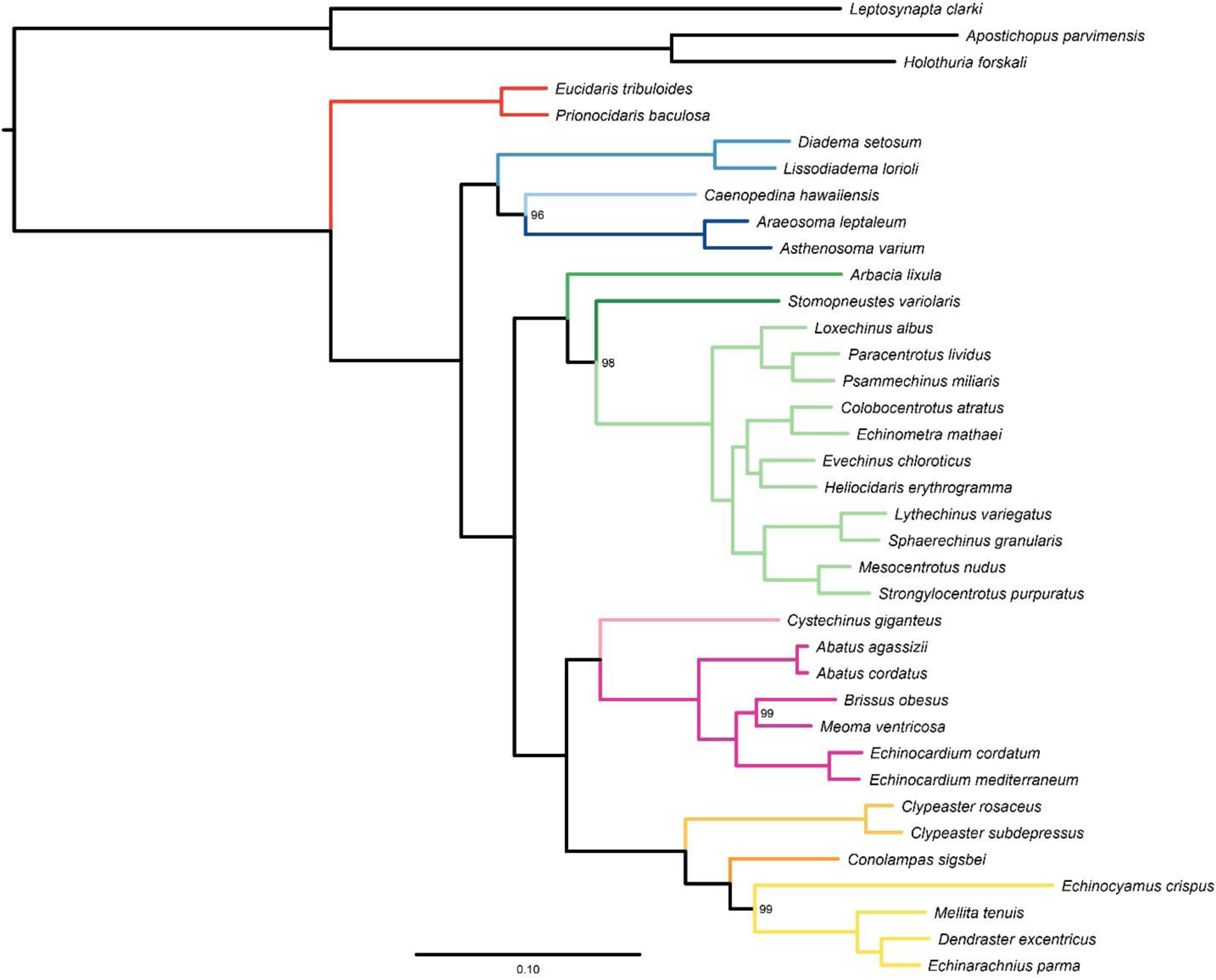
Partitioned maximum likelihood phylogeny of the subsampled 300-loci dataset. Colors are as in Figure 1. Numbers on nodes represent bootstrap frequencies, and are maximum unless shown. Topology is identical to that of Figure 1, even though the dataset was reduced to 12.7% of its original size.

**Figure S11:**
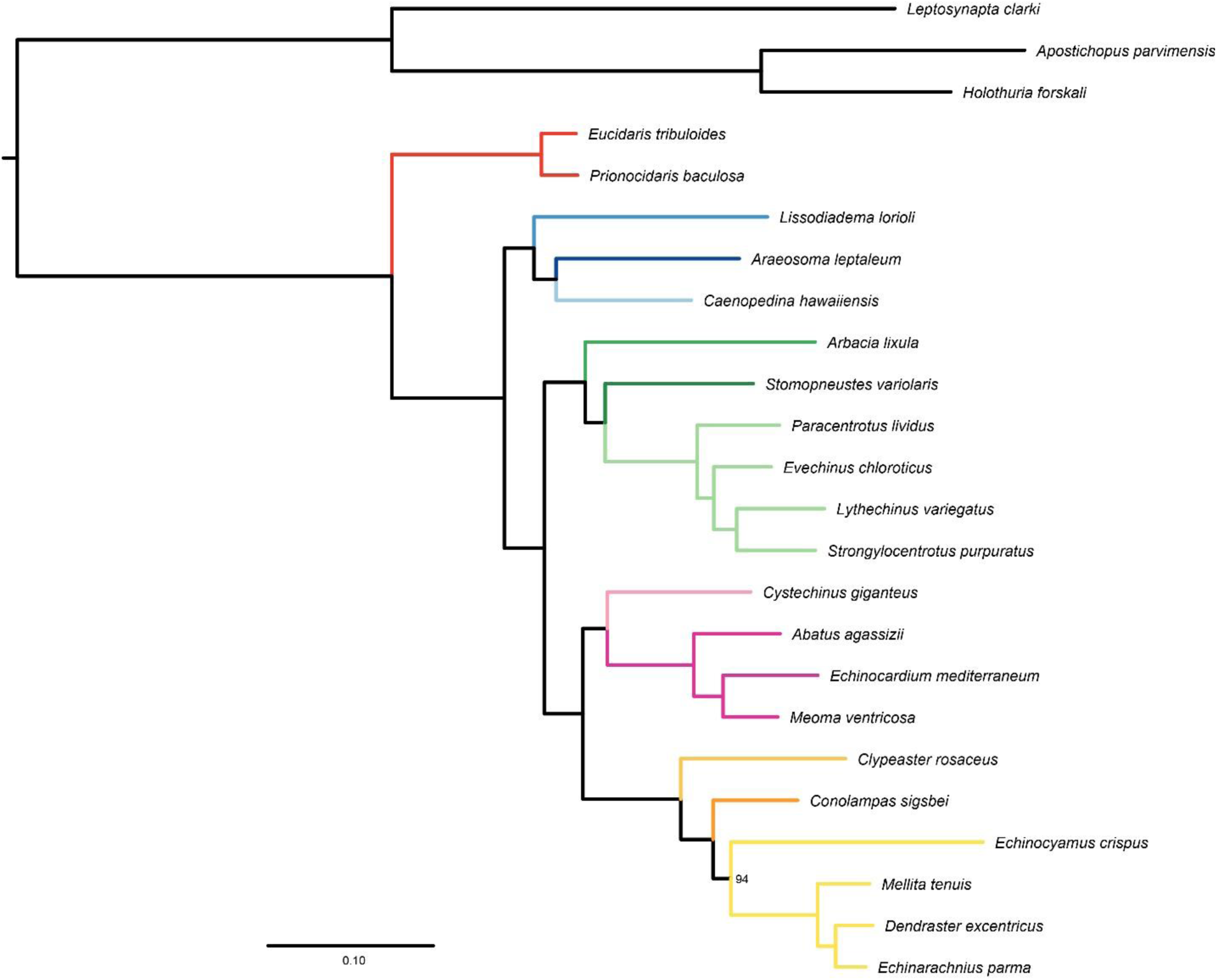
Partitioned maximum likelihood phylogeny of the subsampled 300-loci dataset after reducing taxon sampling to only one terminal per lineage represented in the morphological dataset (i.e., those highlighted in Fig. S3). Colors are as in Figure 1. Numbers on nodes represent bootstrap frequencies, and are maximum unless shown. Topology is again identical to that of Figure 1 and S10.

**Figure S12:**
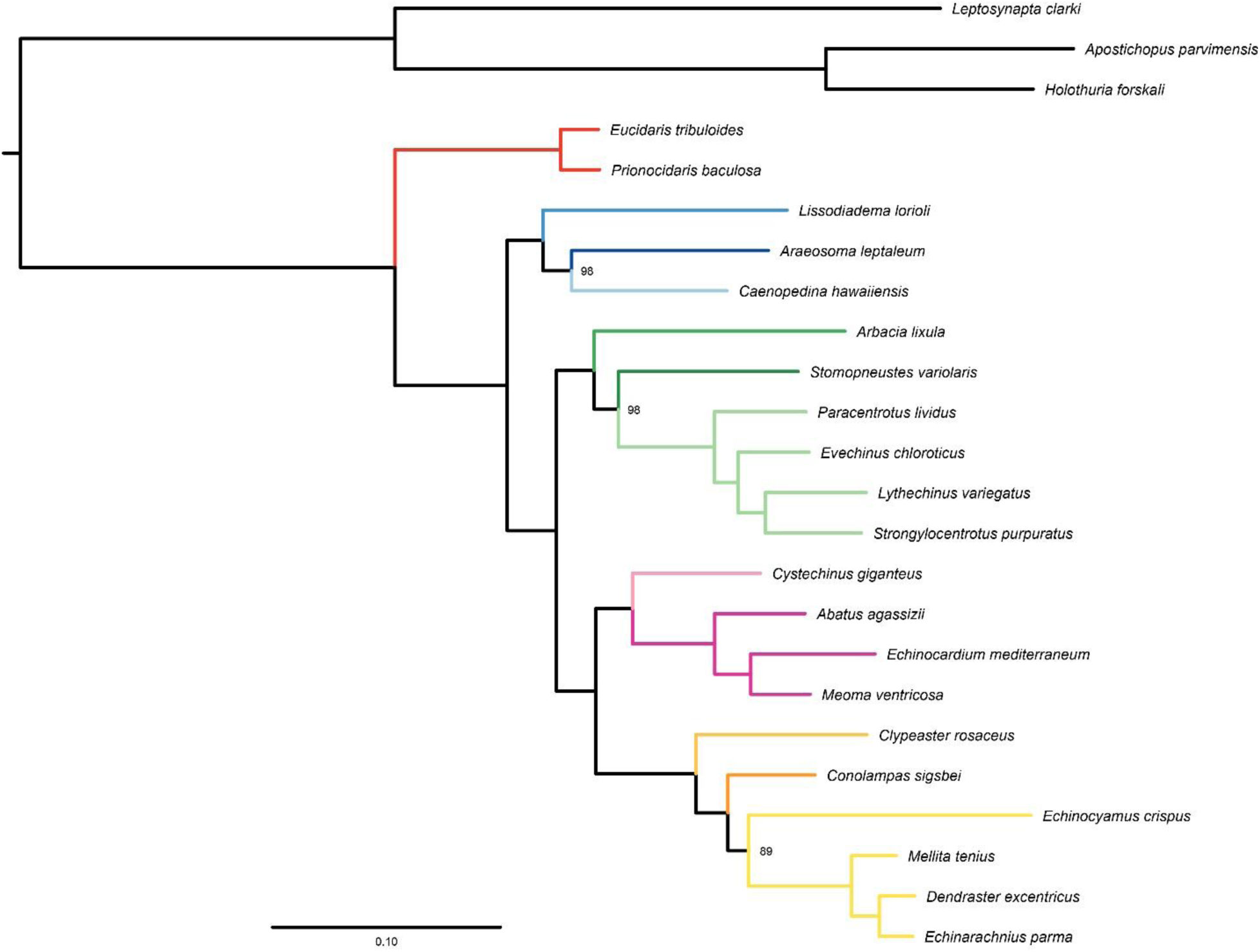
Partitioned maximum likelihood phylogeny of the subsampled 50-loci dataset after reducing taxon sampling to only one terminal per lineage represented in the morphological dataset (i.e., those highlighted in Fig. S3). Colors are as in Figure 1. Numbers on nodes represent bootstrap frequencies, and are maximum unless shown. Topology is again identical to that of Figure 1 and S10-S11, even though the dataset was reduced to 2.1% of its original size.

**Figure S13:**
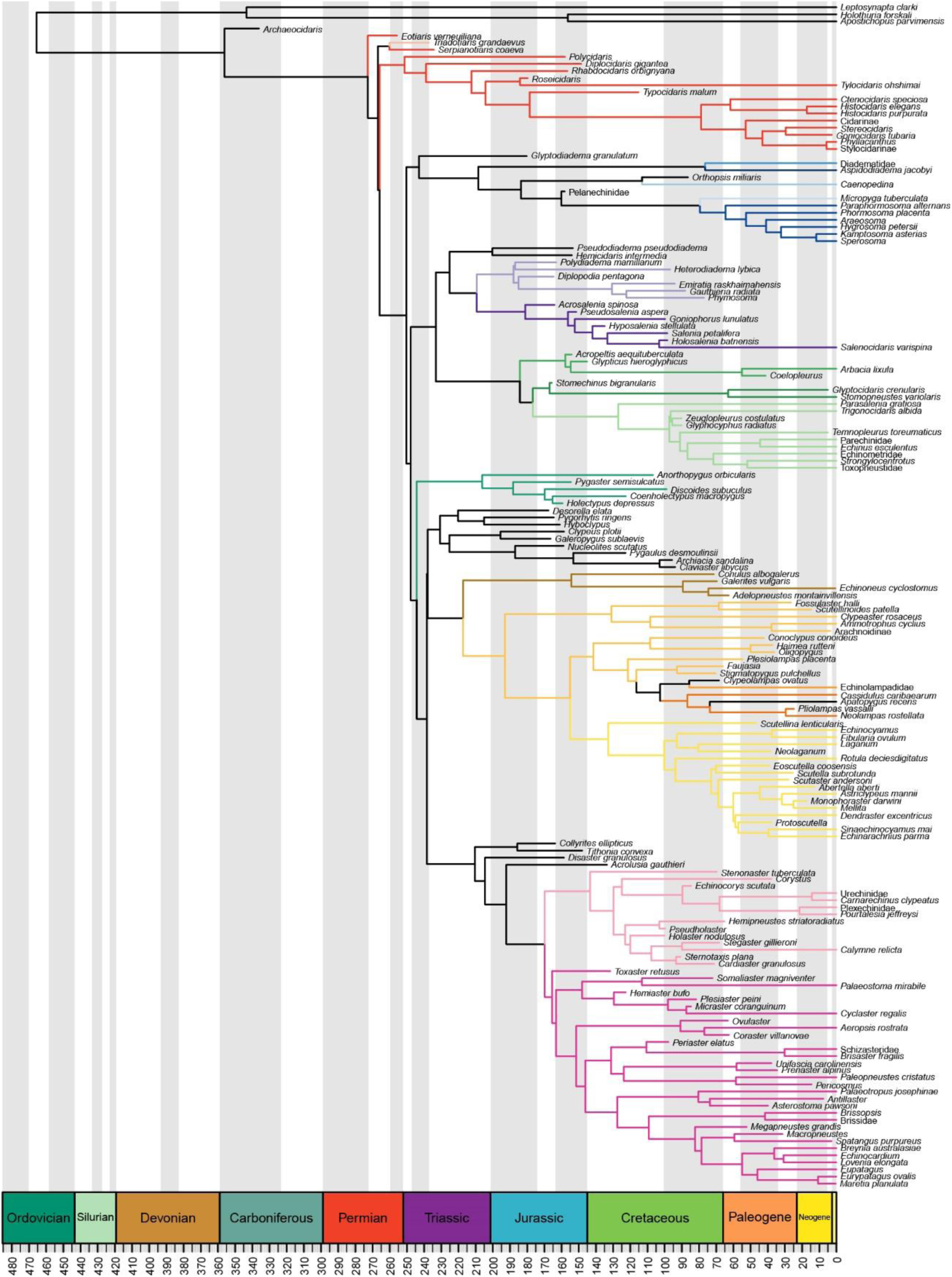
Maximum clade credibility topology of the total-evidence analysis (the majority-rule consensus tree is shown in Figure 5). Branches are colored as in Figure 2.

**Figure S14:**
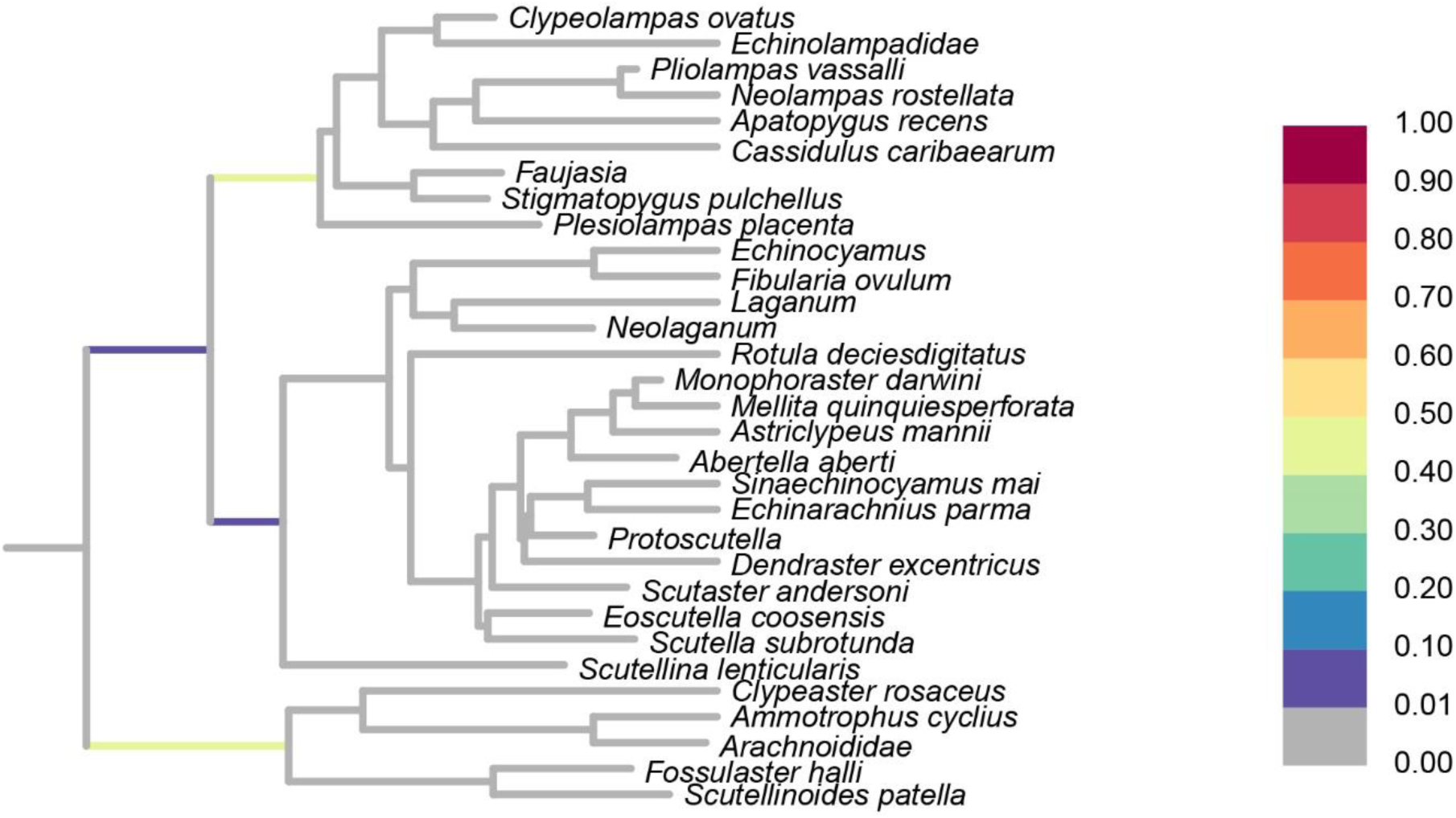
Phylogenetic affinities of Oligopygoida in the total-evidence dated analysis. The tree represents the topology of Neognathostomata in the total-evidence dated analysis after pruning oligopygoids. Branches are colored according to the posterior probability of the position of this clade (see heatmap). Almost equal posterior probabilities are recovered for a position as the sister group to Clypeasteroida (0.495) and Cassiduloida (0.465).

**Figure S15:**
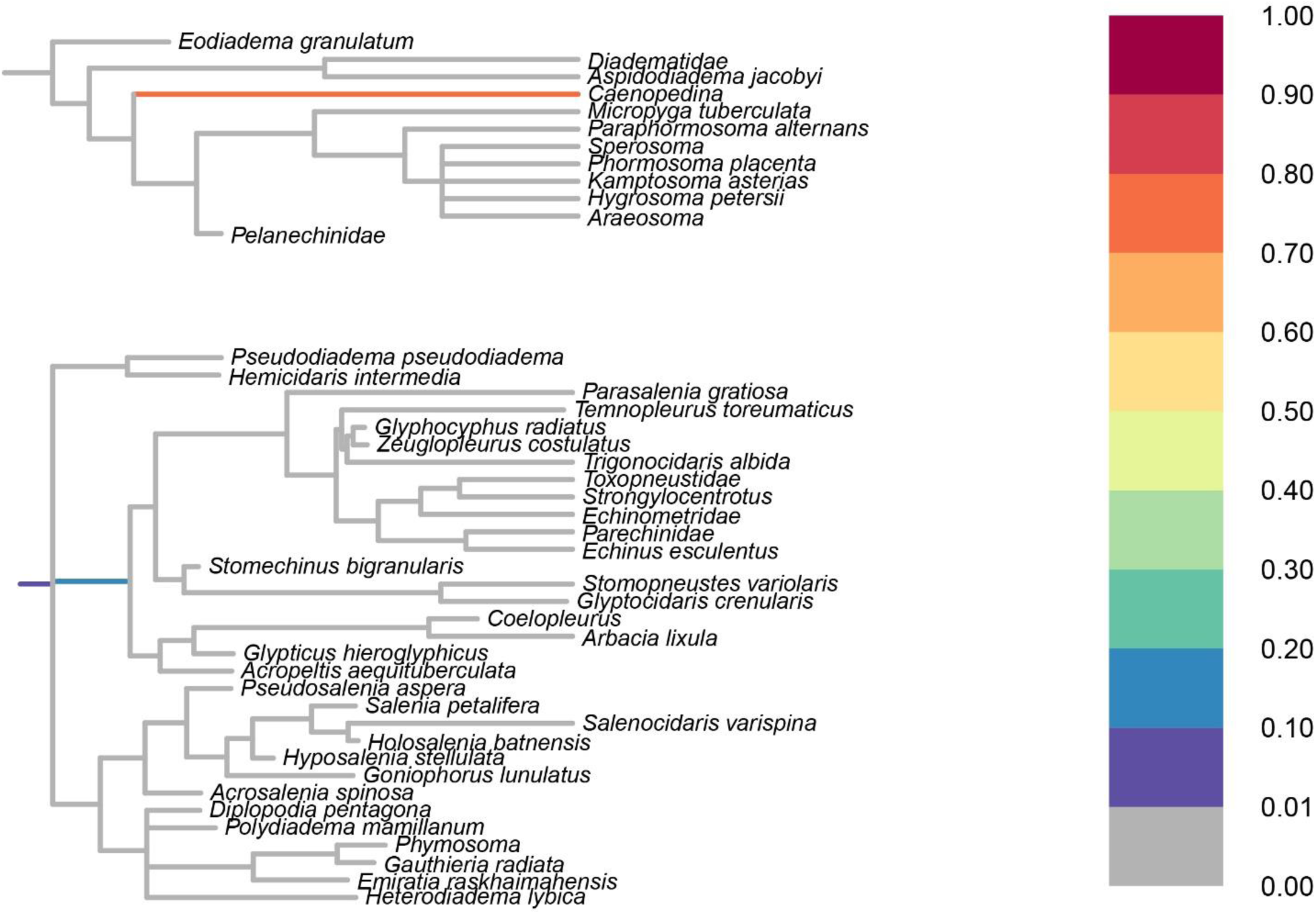
Phylogenetic affinities of *Orthospis miliaris* in the total-evidence dated analysis. The two trees represent different clades to which *Orthopsis miliaris* attaches in the total-evidence dated analysis. Branches are colored according to the posterior probability of the position of this terminal (see heatmap). A position within aulodonta (top, sister to Pedinoida) receives a much higher posterior probability than a relationship to echinaceans, calycineans and allies (bottom).

**Figure S16:**
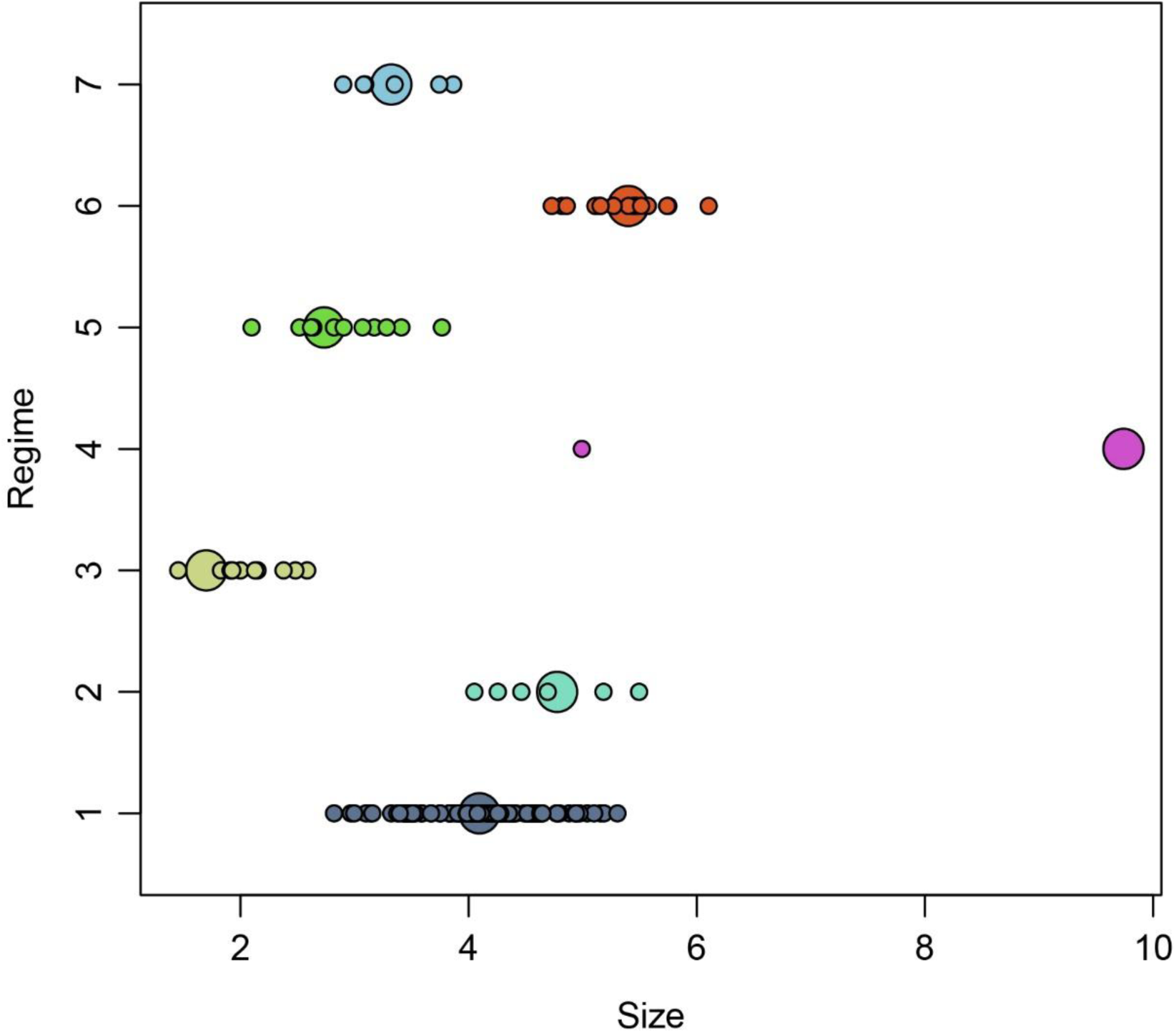
OUM model obtained using the SURFACE algorithm. The model includes 7 regimes, shown in different colors. Body sizes of terminals are shown with small dots, adaptive optima (θ) with large ones. One optimum is very unrealistic, implying attraction to body sizes more than 4 orders of magnitude larger than the largest sampled echinoid. The terminal in question has a very shallow divergence age with its sister clade, forcing the model to accommodate a rapid change with a fixed σ^2^ and α.

**Figure S17:**
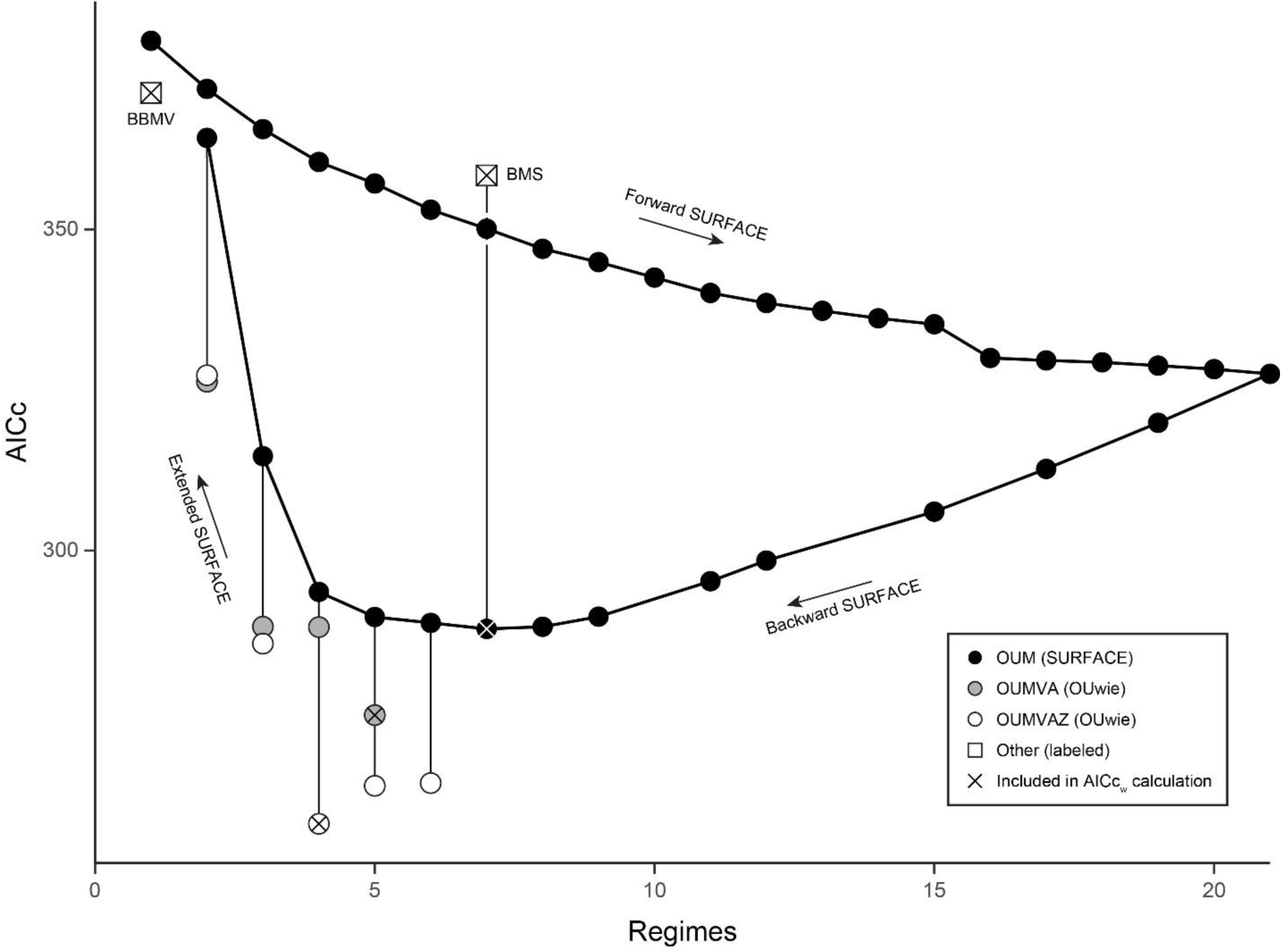
Relative fit (and method followed to obtain) the non-uniform models of body size macroevolution explored. Black dots show the path followed by the forward (regime addition), backwards (regime merging until optimal AICc found) and extended (regime merging beyond optimal AICc) phases of SURFACE. The optimal OUM model returned by SURFACE (with 7 regimes), as well as the suboptimal OUM models obtained through the extended phase (with between 2 and 6 regimes) were used as starting points for the optimization of OUMVA and OUMVAZ models with OUwie. If an AICc score is not shown, model fit was unreliable and the model was excluded (e.g., neither OUMVA nor OUMVAZ models with 7 regimes were successfully fit). Models incorporated into model selection (see Table 2) are identified with an X. The AICc value of the highest fitting non-uniform model (BBMV) is shown for comparison.

**Figure S18:**
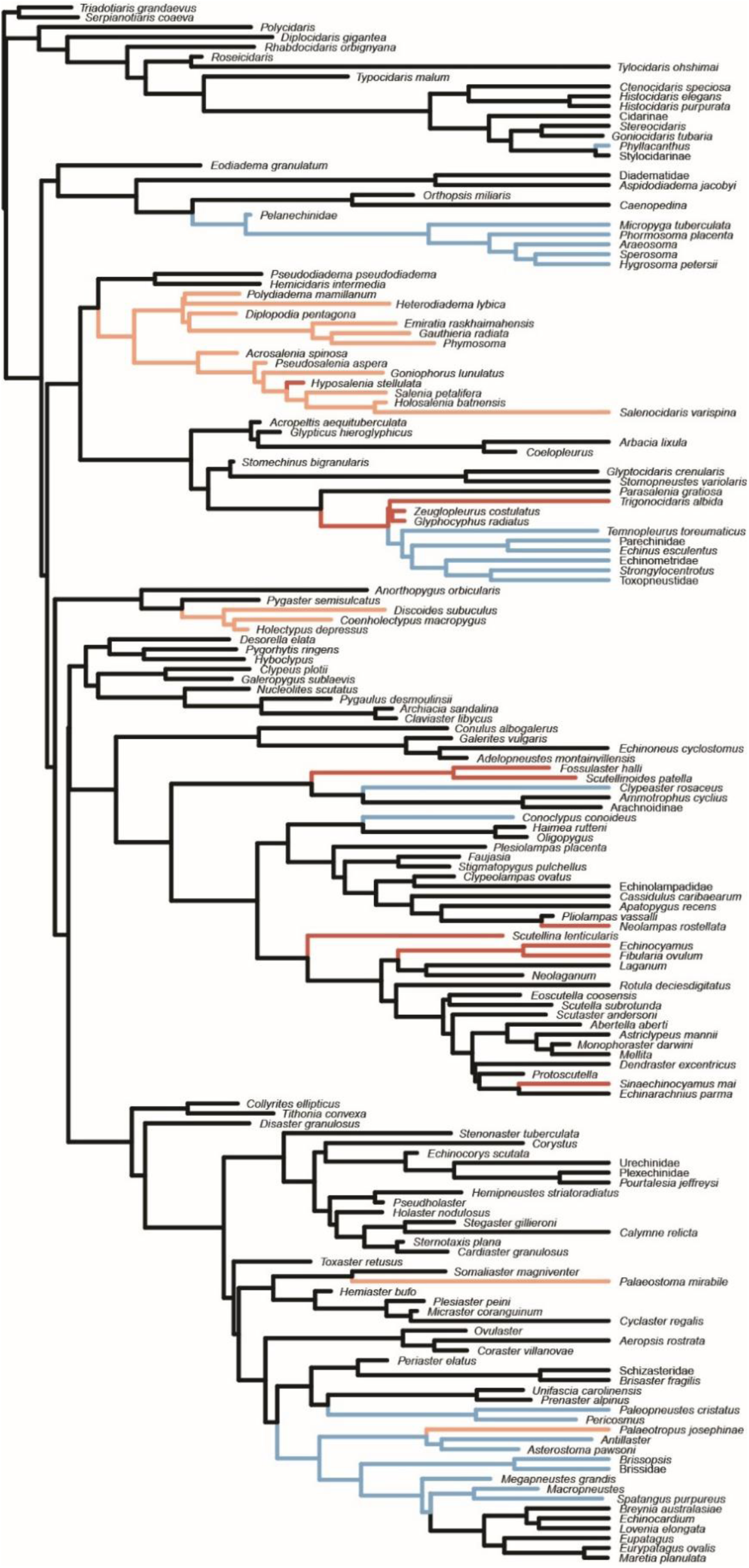
Optimal OUMVAZ model found in the process of model selection. The tree is identical to that of Figure 6 but shows the names of terminals occupying the different regimes.

**Fig. S19:**
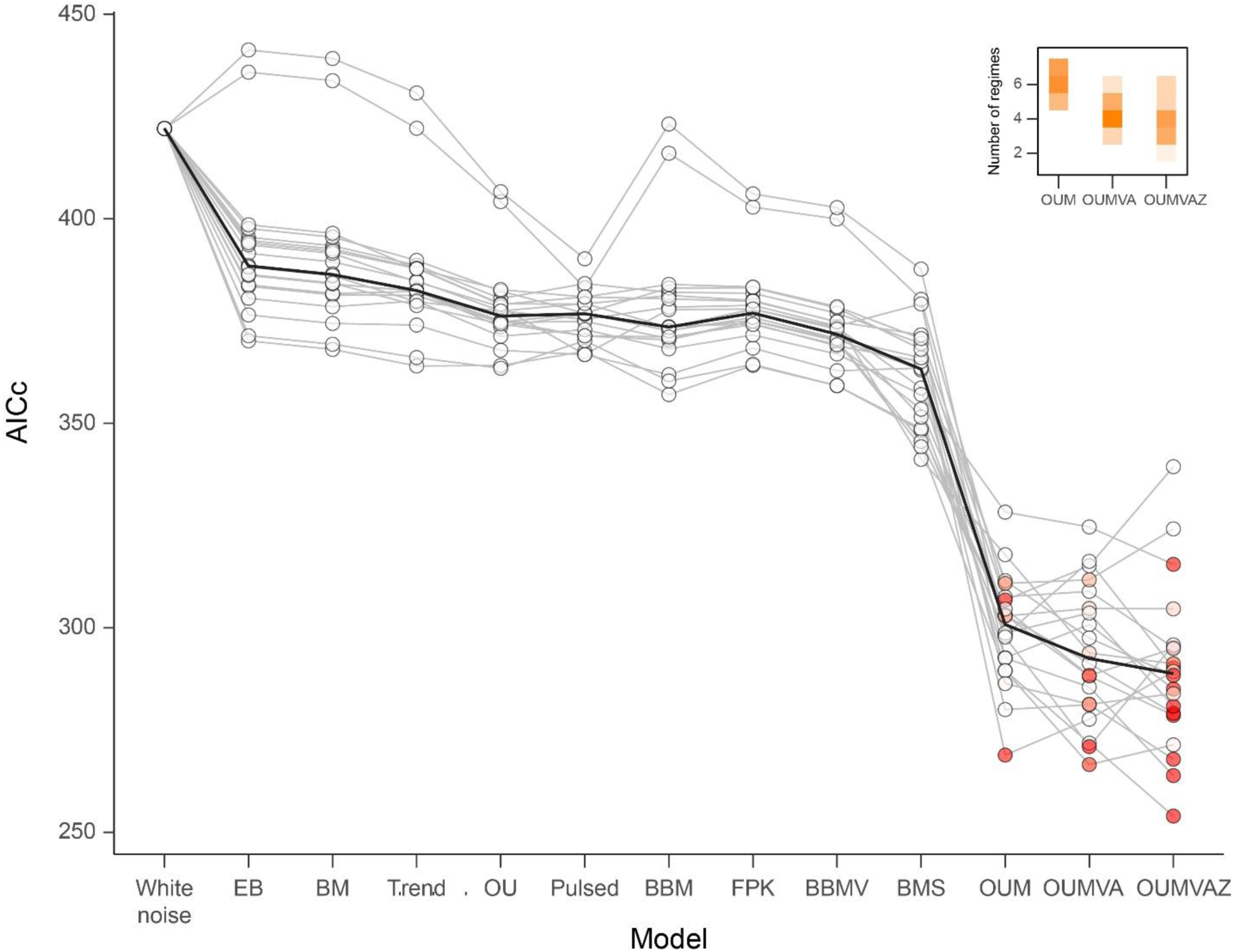
Summary of model fit across 20 topologies randomly sampled from the posterior distribution of the total-evidence dated analysis. Results for the same tree are connected by grey lines, median AICc values are connected by a black line. Dots are colored according to the AICc weight attained by each model for each topology. Across all trees, the added AICc weights of the OUM, OUMVA and OUMVAZ models is 1.00. OUMVA and OUMVAZ models obtained through the newly implemented extended phase of SURFACE, and optimized using OUwie, improved model fit with respect to the OUM model output by SURFACE in 80% of the sampled trees. These models were also systematically less complex (i.e., had less regimes), as shown in the inset.

**Figure S20:**
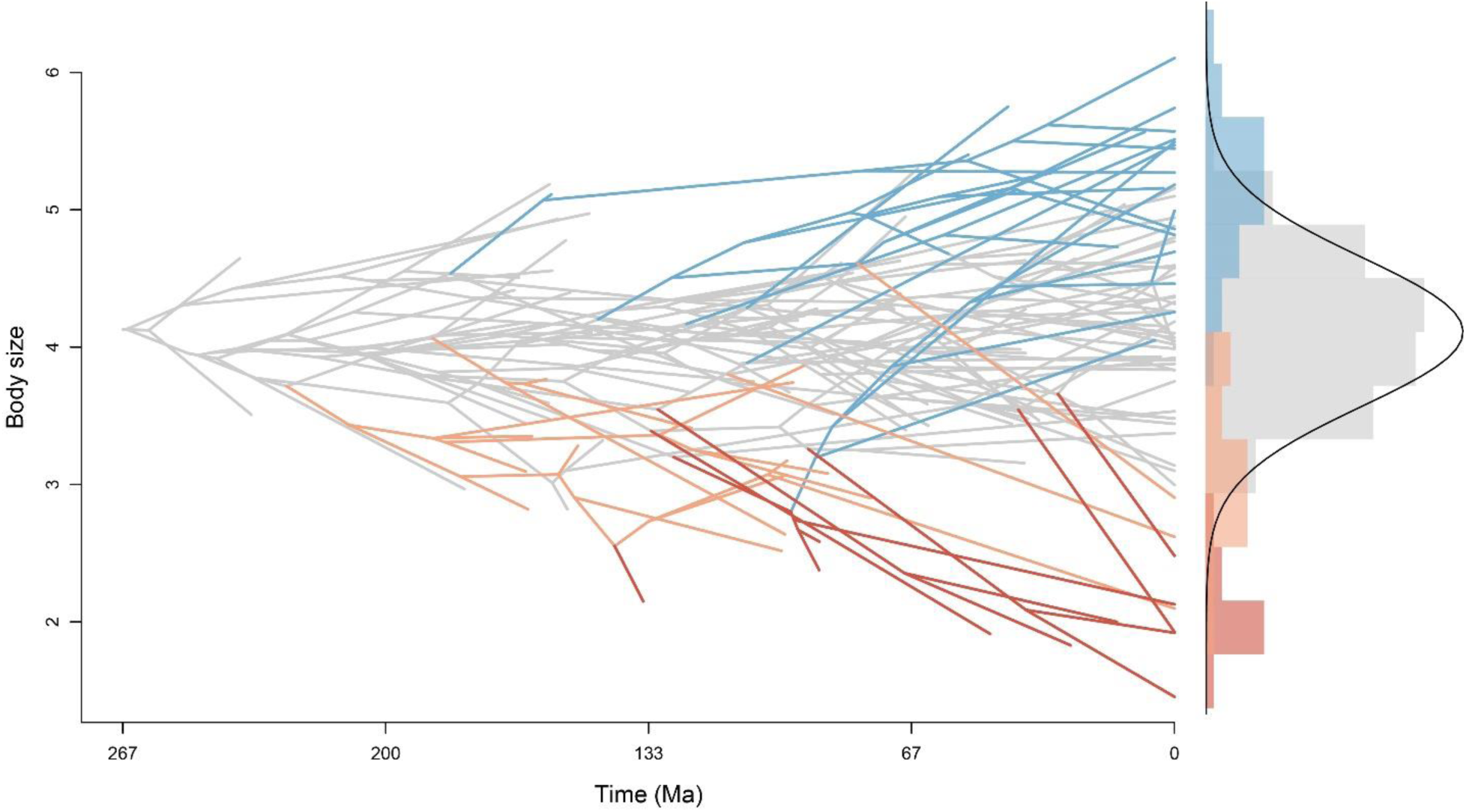
Phenogram showing the pattern of echinoid body size macroevolution. The topology is that of the mcc tree, and lineages are colored according to the adaptive regime they belong to (colored as in Fig. 6). The histogram on the right shows the morphological disparity explored by each regime, and the black line represents the expected disparity of the average size regime (grey) at equilibrium.

**Figure S21:**
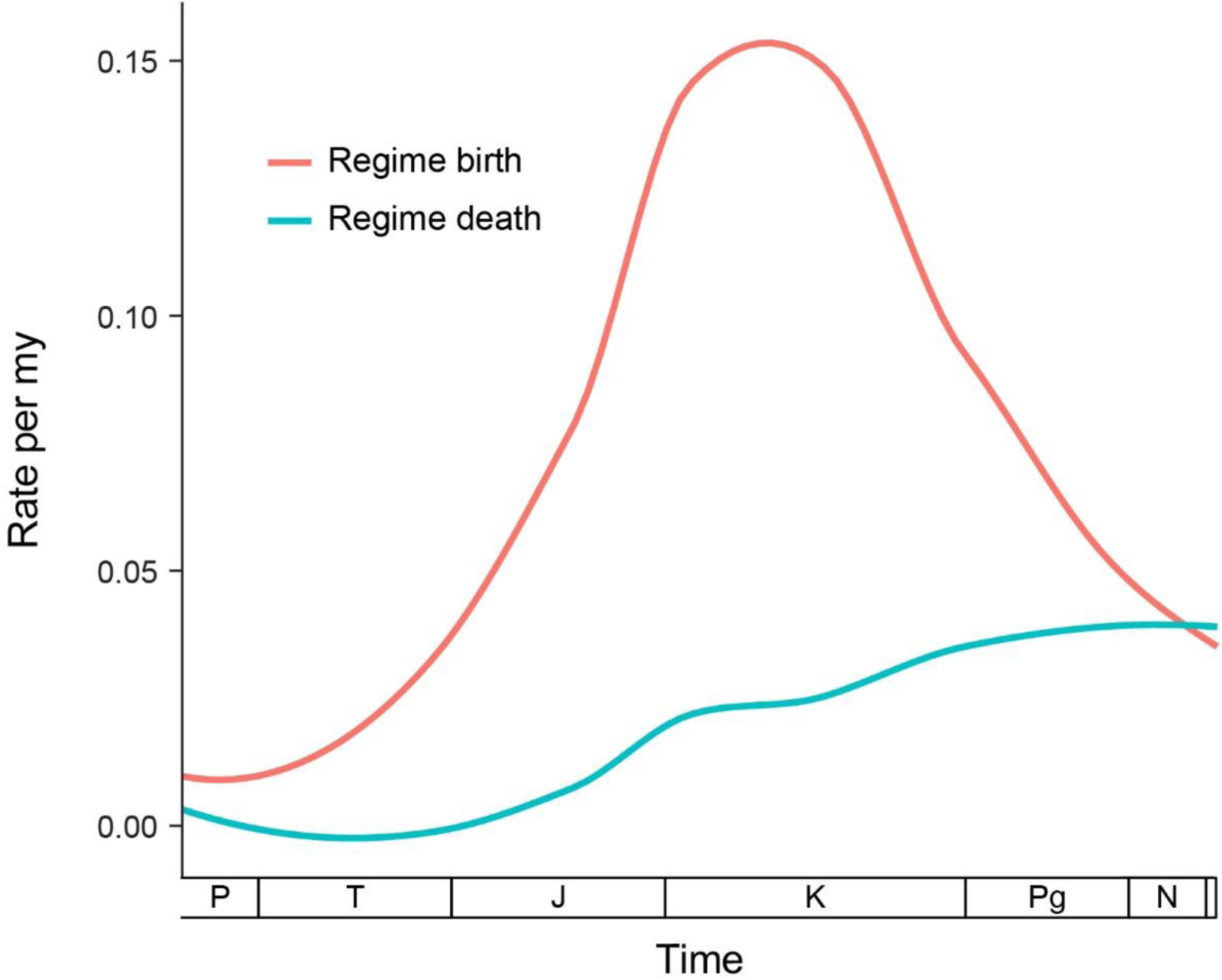
Rate of body size adaptive regime birth and death during the evolutionary history of echinoids. Rates were calculated by counting the number of origins and extinctions of regimes in a 20 my sliding window, taking 5 my steps. Rates are expressed as events per my. The Permian and Triassic have a relatively low birth rate, with a rapid increase in the Jurassic and a peak in the Lower Cretaceous. The birth rate then steadily decreases through the Cenozoic, until matching the death rate in the Recent.

**Figure S22:**
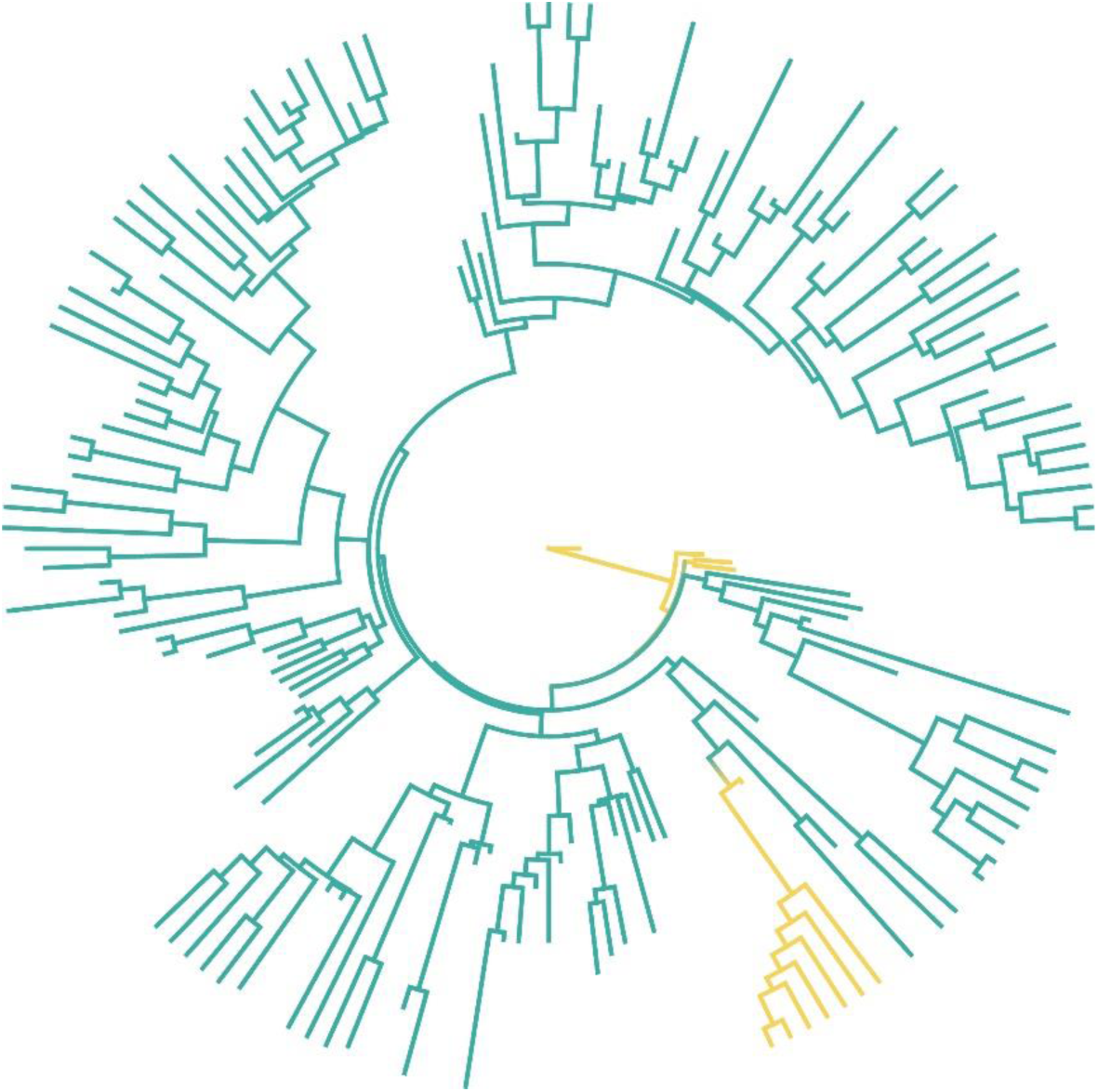
Ancestral state reconstruction of test flexibility. The topology corresponds to the mcc tree of the total-evidence dated analysis (Fig. S13). The character mapped is the condition of the ambulacral-interambulacral suture, which can be imbricate (yellow) or not (teal). Character scorings are those of Character A18 in the matrix of Kroh and Smith (2010). The loss of imbrication along the ambulacral-interambulacral suture is a synapomorphy of crown group Echinoidea as redefined by our analysis. A second origin of imbrication occurs within the clade, in the lineage that gives rise to pelanechinids, micropygoids and echinothurioids. Test flexibility among these is therefore not homologous with the condition that characterizes the echinoid stem group.

**Figure S23:**
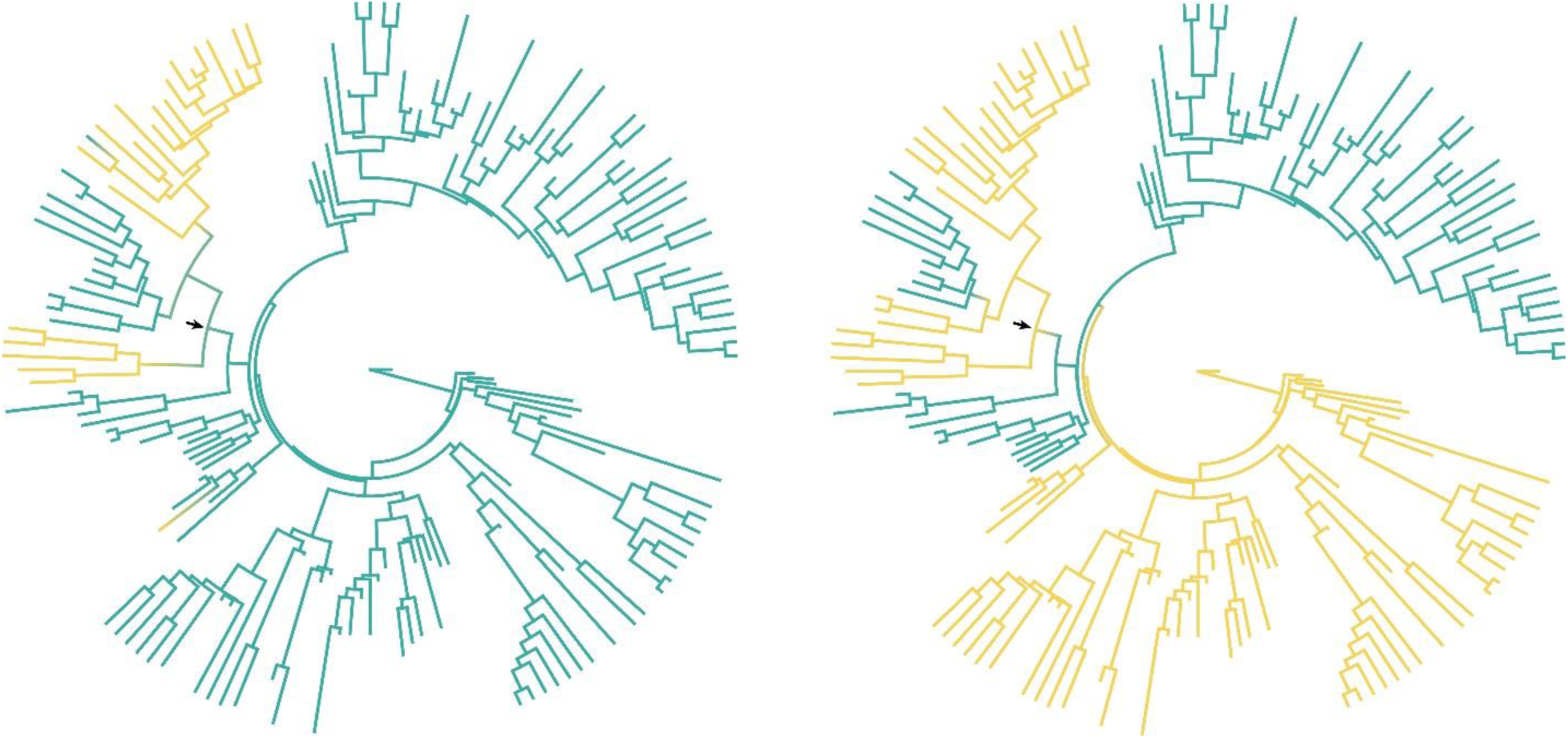
Ancestral state reconstruction of putative synapomorphies historically used to support a clade of Clypeasteroida + Scutelloida (last common ancestor of both shown with an arrow). The topology corresponds to the mcc tree of the total-evidence dated analysis (Fig. S13). Some traits, such as the presence of internal test reinforcements (left; presence = yellow), is inferred to be a case of convergent evolution between sand dollars and sea biscuits. Character scorings are those of Character A30 and A31 in the matrix of Kroh and Smith (2010). Others, such as the presence of Aristotle’s lantern in adults (right, presence = yellow) are a true innovation of the last common ancestor of these two clades, but have secondarily reversed in some of its descendants (Cassiduloida, as defined in this study). Character scorings are those of Character F1 in the matrix of Kroh and Smith (2010).

## References

Agassiz A. 1873. Revison of the Echini. Mem. Mus. Comp. Zool. Harv. Coll. 3:379–628.

Arcila D., Pyron R.A., Tyler J.C., Ortí G., Betancur-R R. 2015. An evaluation of fossil tip-dating versus node-age calibrations in tetraodontiform fishes (Teleostei: Percomorphaceae). Mol. Phylogenet. Evol. 82:131–145.

Barras C.G. 2007. Phylogeny of the Jurassic to Early Cretaceous ‘disasteroid’ echinoids (Echinoidea; Echinodermata) and the origins of spatangoids and holasteroids. J. Syst. Palaeontol. 5:133–161.

Bastide P., Ané C., Robin S., Mariadassou M. 2018. Inference of adaptive shifts for multivariate correlated traits. Syst. Biol. 67:662–680.

Beaulieu J.M., Jhwueng D.C., Boettiger C., O’Meara B.C. 2012. Modeling stabilizing selection: expanding the Ornstein–Uhlenbeck model of adaptive evolution. Evolution 66:2369–2383.

Beaulieu J.M., O’Meara B.C. 2019. OUwie: Analysis of Evolutionary Rates in an OU Framework. R package version 1.53. https://CRAN.R-project.org/package=OUwie.

Benson R.B., Hunt G., Carrano M.T., Campione N. 2018. Cope’s rule and the adaptive landscape of dinosaur body size evolution. Palaeontology 61:13–48.

Blomberg S.P., Garland T., Ives A.R. 2003. Testing for phylogenetic signal in comparative data: behavioral traits are more labile. Evolution 57:717–745.

Boivin S., Saucède T., Laffont R., Steimetz E., Neige P. 2018. Diversification rates indicate an early role of adaptive radiations at the origin of modern echinoid fauna. PLoS One, 13:e0194575.

Bolger A.M., Lohse M., Usadel B. 2014. Trimmomatic: a flexible trimmer for Illumina sequence data. Bioinformatics 30:2114–2120.

Bollback J.P. 2006. SIMMAP: stochastic character mapping of discrete traits on phylogenies. BMC Bioinformatics 7:88.

Boucher F.C. 2019. BBMV: an R package for the estimation of macroevolutionary landscapes. Ecography 42:558–564.

Boucher F.C., Démery V. 2016. Inferring bounded evolution in phenotypic characters from phylogenetic comparative data. Syst. Biol. 65:651–661.

Boucher F.C., Démery V., Conti E., Harmon L.J., Uyeda J. 2017. A general model for estimating macroevolutionary landscapes. Syst. Biol. 67:304–319.

Bronstein O., Kroh A. 2019. The first mitochondrial genome of the model echinoid Lytechinus variegatus and insights into Odontophoran phylogenetics. Genomics 111:710–718.

Brosseau O., Murienne J., Pichon D., Vidal N., Eléaume M., Ameziane N. 2012. Phylogeny of Cidaroida (Echinodermata: Echinoidea) based on mitochondrial and nuclear markers. Org. Divers. Evol. 12:155–165.

Butler M.A., King A.A. 2004. Phylogenetic Comparative Analysis: A Modeling Approach for Adaptive Evolution. Am. Nat. 164:683–695.

Calder W.A. 1996. Size, function, and life history. Harvard University Press, Cambridge.

Carpenter R.C. 1981. Grazing by Diadema antillarum (Phillipi) and its effects on the benthic algal community. J. Mar. Res. 39:749–765

Ceccarelli F.S., Mongiardino Koch N., Soto E.M., Barone M.L., Arnedo M.A., Ramírez M.J. 2019. The grass was greener: Repeated evolution of specialized morphologies and habitat shifts in ghost spiders following grassland expansion in South America. Syst. Biol. 68:63–77.

Chernomor O., von Haeseler A., Minh B.Q. 2016. Terrace aware data structure for phylogenomic inference from supermatrices. Syst. Biol. 65:997–1008.

Church S.H., Donoughe S., de Medeiros B.A., Extavour C.G. 2019. Insect egg size and shape evolve with ecology but not developmental rate. Nature 571:58–62.

Clark M.S., Suckling C.C., Cavallo A., Mackenzie C.L., Thorne M.A, Davies A.J., Peck L.S. 2019. Molecular mechanisms underpinning transgenerational plasticity in the green sea urchin Psammechinus miliaris. Sci. Rep. 9:952.

Colombo M., Damerau M., Hanel R., Salzburger W., Matschiner M. 2015. Diversity and disparity through time in the adaptive radiation of Antarctic notothenioid fishes. J. Evol. Biol. 28:376–394.

Cooper N., Purvis A. 2010. Body size evolution in mammals: complexity in tempo and mode. Am. Nat. 175:727–738.

Cummins C.A., McInerney J.O. 2011. A method for inferring the rate of evolution of homologous characters that can potentially improve phylogenetic inference, resolve deep divergence and correct systematic biases. Syst. Biol. 60:833–844.

Darriba D., Posada D., Kozlov A.M., Stamatakis A., Morel B., Flouri T. 2020. ModelTest-NG: a new and scalable tool for the selection of DNA and protein evolutionary models. Mol. Biol. Evol. 37:291–294.

Davis A.M., Betancur-R R. 2017. Widespread ecomorphological convergence in multiple fish families spanning the marine–freshwater interface. Proc. R. Soc. Lond. B Biol. Sci. 284:20170565.

Di Franco A., Poujol R., Baurain D., Philippe H. 2019. Evaluating the usefulness of alignment filtering methods to reduce the impact of errors on evolutionary inferences. BMC Evol. Biol. 19:21.

Dunn C.W., Howison M., Zapata F. 2013. Agalma: an automated phylogenomics workflow. BMC Bioinformatics 14:330.

Durham J.W. 1966a. Classification. In: Treatise on Invertebrate Paleontology, Part U. Moore RC, Ed. University of Kansas Press, Lawrence. pp. 270–296.

Durham J.W. 1966b. Clypeasteroids. Treatise on invertebrate paleontology, Part U. In: Treatise on Invertebrate Paleontology, Part U. Moore RC, Ed. University of Kansas Press, Lawrence. pp. 450–491.

Durham J.W., Melville R. 1957. A classification of echinoids. J. Paleontol. 31:242–272.

Eble G.J. 2000. Contrasting evolutionary flexibility in sister groups: disparity and diversity in Mesozoic atelostomate echinoids. Paleobiology 26:56–79.

Edmunds P.J., Carpenter R.C. 2001. Recovery of Diadema antillarum reduces macroalgal cover and increases abundance of juvenile corals on a Caribbean reef. Proc. Natl. Acad. Sci. 98:5067–5071.

Emlet R. 2002. Ecology of adult sea urchins. In: Sea Urchin: From Basic Biology to Aquaculture. Yokota Y, Matranga V, Smolenicka Z, Eds. AA Balkema, Rotterdam. pp. 111–114.

Erkenbrack E.M., Thompson J.R. 2019. Cell type phylogenetics informs the evolutionary origin of echinoderm larval skeletogenic cell identity. Commun. Biol. 2:160.

Erwin D.H. 1994. The Permo–Triassic extinction. Nature 367:231–236.

Fell H.B. 1966. Diadematacea. In: Treatise on Invertebrate Paleontology, Part U. Moore RC, Ed. University of Kansas Press, Lawrence. pp. 340–366.

Fell H.B, Pawson D.L. 1966. Echinacea. In: Treatise on Invertebrate Paleontology, Part U. Moore RC, Ed. University of Kansas Press, Lawrence. pp. 367-374.

Felsenstein J. 1985. Phylogenies and the comparative method. Am. Nat. 125:1–15.

Finnegan S., Droser M.L. 2008. Body size, energetics, and the Ordovician restructuring of marine ecosystems. Paleobiology 34:342–359.

Foote M. 1995. Morphological diversification of Paleozoic crinoids. Paleobiology 21:273–299

Freckleton R.P., Harvey P.H., Pagel M. 2002. Phylogenetic analysis and comparative data: a test and review of evidence. Am. Nat. 160:712–726.

Gaitán-Espitia J.D., Sánchez R., Bruning P., Cárdenas L. 2016. Functional insights into the testis transcriptome of the edible sea urchin Loxechinus albus. Sci. Rep. 6:36516.

Gavryushkina A., Heath T.A., Ksepka D.T., Stadler T., Welch D., Drummond A.J. 2017. Bayesian total-evidence dating reveals the recent crown radiation of penguins. Syst. Biol. 66:57–73.

Gladfelter W. 1978. General ecology of the cassiduloid urchin Cassidulus caribbearum. Mar. Biol. 47:149–160.

Godoy P.L., Benson R.B., Bronzati M., Butler R.J. 2019. The multi-peak adaptive landscape of crocodylomorph body size evolution. BMC Evol. Biol. 19:167.

Goloboff P.A. 1993. Estimating character weights during tree search. Cladistics 9:83–91.

Goloboff P.A., Catalano S.A. 2016. TNT version 1.5, including a full implementation of phylogenetic morphometrics. Cladistics 32:221–238.

Grabherr M.G., Haas B.J., Yassour M., Levin J.Z., Thompson D.A., Amit I., Adiconis X., Fan L., Raychowdhury R., Zeng Q. 2011. Full-length transcriptome assembly from RNA-Seq data without a reference genome. Nat. Biotechnol. 29:644–652.

Gregory J. 1897. On the Affinities of the Echinothuridæ; and on Pedinothuria and Helikodiadema, two new Genera of Echinoidea. Quart. J. Geol. Soc. 53:112–122.

Grun T.B., von Scheven M., Bischoff M., Nebelsick J.H. 2018. Structural stress response of segmented natural shells: a numerical case study on the clypeasteroid echinoid Echinocyamus pusillus. J. R. Soc. Interface 15:20180164.

Guang A., Howison M., Zapata F., Lawrence C.E., Dunn C. 2017. Revising transcriptome assemblies with phylogenetic information in Agalma 1.0. bioRxiv 202416.

Hansen T.F. 1997. Stabilizing selection and the comparative analysis of adaptation. Evolution 51:1341–1351.

Hansen T.F. 2012. Adaptive landscapes and macroevolutionary dynamics. In: The adaptive landscape in evolutionary biology. Svensson EI, Calsbeek R, Eds. Oxford University Press, Oxford. pp. 205:221.

Harmon L.J., Losos J.B., Jonathan Davies T., Gillespie R.G., Gittleman J.L., Bryan Jennings W., Kozak K.H., McPeek M.A., Moreno-Roark F., Near T.J. 2010. Early bursts of body size and shape evolution are rare in comparative data. Evolution 64:2385–2396.

Harmon L.J., Schulte J.A., Larson A., Losos J.B. 2003. Tempo and mode of evolutionary radiation in iguanian lizards. Science 301:961–964.

Harmon L.J., Weir J.T., Brock C.D., Glor R.E., Challenger W. 2008. GEIGER: investigating evolutionary radiations. Bioinformatics 24:129–131.

Harrold C., Pearse J.S. 1987. The ecological role of echinoderms in kelp forests. In: Echinoderm studies 2. Jangoux M, Laurence JM, Eds. AA Balkema, Rotterdam. pp. 137–233.

Hawkins H.L. 1920. Morphological Studies on the Echinoidea Holectypoida and their Allies: X. On Apatopygus gen. nov. and the affinities of some recent Nucleolitoida and Cassiduloida. Geol. Mag. 57:393–401.

Heath T.A., Huelsenbeck J.P., Stadler T. 2014. The fossilized birth–death process for coherent calibration of divergence-time estimates. Proc. Natl. Acad. Sci. 111:2957–2966.

Ho L.S.T., Ané C. 2014. Intrinsic inference difficulties for trait evolution with Ornstein-Uhlenbeck models. Methods Ecol. Evol. 5:1133–1146.

Hoang D.T., Chernomor O., Von Haeseler A., Minh B.Q., Vinh L.S. 2017. UFBoot2: improving the ultrafast bootstrap approximation. Mol. Biol. Evol. 35:518–522.

Hollertz K., Duchêne J.-C. 2001. Burrowing behaviour and sediment reworking in the heart urchin Brissopsis lyrifera Forbes (Spatangoida). Mar. Biol. 139:951–957.

Hopkins M.J., Smith A.B. 2015. Dynamic evolutionary change in post-Paleozoic echinoids and the importance of scale when interpreting changes in rates of evolution. Proc. Natl. Acad. Sci. 112:3758–3763.

Hughes M., Gerber S., Wills M.A. 2013. Clades reach highest morphological disparity early in their evolution. Proc. Natl. Acad. Sci. 110:13875–13879.

Hughes T.P. 1994. Catastrophes, phase shifts, and large-scale degradation of a Caribbean coral reef. Science 265:1547–1551.

Hunt G., Carrano M.T. 2010. Models and methods for analyzing phenotypic evolution in lineages and clades. Spec. Pap. Pal. Soc. 16:245–269.

Ingram T., Mahler D.L. 2013. SURFACE: detecting convergent evolution from comparative data by fitting Ornstein-Uhlenbeck models with stepwise Akaike Information Criterion. Methods Ecol. Evol. 4:416–425.

Jackson R.T. 1912. Phylogeny of the Echini, with a revision of Palaeozoic species. Mem. Boston Soc. Nat. Hist. 7:1–491.

Jensen M. 1981. Morphology and classification of Euechinoidea Bronn, 1860—a cladistic analysis. Vidensk. Meddel. Dansk Naturhist. Foren. Kjøbenhavn 143:7–99.

Jombart T., Dray S. 2010. adephylo: exploratory analyses for the phylogenetic comparative method. Bioinformatics 26:1–21.

Kalyaanamoorthy S., Minh B.Q., Wong T.K., von Haeseler A., Jermiin L.S. 2017. ModelFinder: fast model selection for accurate phylogenetic estimates. Nat. Methods 14:587–589.

Kealy S., Beck R. 2017. Total evidence phylogeny and evolutionary timescale for Australian faunivorous marsupials (Dasyuromorphia). BMC Evol. Biol. 17:240.

Khabbazian M., Kriebel R., Rohe K., Ané C. 2016. Fast and accurate detection of evolutionary shifts in Ornstein–Uhlenbeck models. Methods Ecol. Evol. 7:811–824.

Kidwell S.M., Baumiller T. 1990. Experimental disintegration of regular echinoids: roles of temperature, oxygen, and decay thresholds. Paleobiology 16:247–271.

Kier P.M. 1962. Revision of the cassiduloid echinoids. Smithson. Misc. Collect. 144:1–262.

Kier P.M. 1966. Cassiduloids. In: Treatise on Invertebrate Paleontology, Part U. Moore RC, Ed. University of Kansas Press, Lawrence. pp. 492–523.

Kier P.M. 1967. Revision of the oligopygoid echinoids. Smithson. Misc. Collect. 152:1–147.

Kier P.M. 1974. Evolutionary trends and their functional significance in the post-Paleozoic echinoids. J. Paleontol. 48:1–95.

Kier PM. 1977. Triassic echinoids. Smithson. Contrib. Paleobiol. 30:1–88.

King B., Beck R.M. 2019. Bayesian tip-dated phylogenetics: topological effects, stratigraphic fit and the early evolution of mammals. bioRxiv 533885.

King B., Qiao T., Lee M.S., Zhu M., Long J.A. 2017. Bayesian morphological clock methods resurrect placoderm monophyly and reveal rapid early evolution in jawed vertebrates. Syst. Biol. 66:499–516.

Klopfstein S., Spasojevic T. 2019. Illustrating phylogenetic placement of fossils using RoguePlots: An example from ichneumonid parasitoid wasps (Hymenoptera, Ichneumonidae) and an extensive morphological matrix. PLoS One 14:e0212942.

Kocot K.M., Struck T.H., Merkel J., Waits D.S., Todt C., Brannock P.M., Weese D.A., Cannon J.T., Moroz L.L., Lieb B., Halanych, K.M. 2017. Phylogenomics of Lophotrochozoa with Consideration of Systematic Error. Syst. Biol. 66:256–282.

Kozlov A.M., Darriba D., Flouri T., Morel B., Stamatakis A. 2019. RAxML-NG: a fast, scalable and user-friendly tool for maximum likelihood phylogenetic inference. Bioinformatics 35:4453–4455.

Kroh A. 2020. Phylogeny and classification of echinoids. In: Sea Urchins: Biology and Ecology, 4th edition. Lawrence JM, Ed. Academic Press, Cambridge. pp. 1–17.

Kroh A., Mooi R. 2019. World Echinoidea Database. Accessed at http://www.marinespecies.org/echinoidea on 2019-08-12.

Kroh A., Smith A.B. 2010. The phylogeny and classification of post-Palaeozoic echinoids. J Syst. Palaeont. 8:147–212.

Kück P., Struck T.H. 2014. BaCoCa–A heuristic software tool for the parallel assessment of sequence biases in hundreds of gene and taxon partitions. Mol. Phylogenet. Evol. 70:94–98.

Kudtarkar P., Cameron R.A. 2017. Echinobase: an expanding resource for echinoderm genomic information. Database 2017:bax074.

Landis M.J., Schraiber J.G. 2017. Pulsed evolution shaped modern vertebrate body sizes. Proc. Natl. Acad. Sci. 114:13224–13229.

Lanfear R., Frandsen P.B., Wright A.M., Senfeld T., Calcott B. 2017. PartitionFinder 2: new methods for selecting partitioned models of evolution for molecular and morphological phylogenetic analyses. Mol. Biol. Evol. 34:772–773.

Lawton JH, Jones CG. 1995. Linking species and ecosystems: organisms as ecosystem engineers. In: Linking species & ecosystems. Jones CG, Lawton JH, Eds. Springer, New York. pp. 141–150.

Lee M.S., Palci A. 2015. Morphological phylogenetics in the genomic age. Curr. Biol. 25:R922–R929.

Lessios H.A., Garrido M.J., Kessing B.D. 2001. Demographic history of Diadema antillarum, a keystone herbivore on Caribbean reefs. Proc. R. Soc. Lond. B Biol. Sci. 268:2347–2353.

Lewis P.O. 2001. A likelihood approach to estimating phylogeny from discrete morphological character data. Syst. Biol. 50:913–925.

Lin J.-P., Tsai M.-H., Kroh A., Trautman A., Machado D.J., Chang L.-Y., Reid R., Lin K.-T., Bronstein O., Lee S.-J, Janies D. 2019. The first complete mitochondrial genome of the sand dollar Sinaechinocyamus mai (Echinoidea: Clypeasteroida). Genomics doi:10.1016/j.ygeno.2019.10.007.

Ling S., Scheibling R., Rassweiler A., Johnson C., Shears N., Connell S., Salomon A., Norderhaug K., Pérez-Matus A., Hernández J. 2015. Global regime shift dynamics of catastrophic sea urchin overgrazing. Philos. Trans. R. Soc. Lond. B Biol. Sci. 370:20130269.

Littlewood D.T.J., Smith A.B. 1995. A combined morphological and molecular phylogeny for sea urchins (Echinoidea: Echinodermata). Philos. Trans. R. Soc. Lond. B Biol. Sci. 347:213–234.

Lohrer A.M., Thrush S.F., Gibbs M.M. 2004. Bioturbators enhance ecosystem function through complex biogeochemical interactions. Nature 431:1092–1095.

Mahler D.L., Ingram T., Revell L.J, Losos J.B. 2013. Exceptional convergence on the macroevolutionary landscape in island lizard radiations. Science 341:292–295.

Mai U., Mirarab S. 2018. TreeShrink: fast and accurate detection of outlier long branches in collections of phylogenetic trees. BMC Genomics 19:272.

Mayrose I., Graur D., Ben-Tal N., Pupko T. 2004. Comparison of site-specific rate-inference methods for protein sequences: empirical Bayesian methods are superior. Mol. Biol. Evol. 21:1781–1791.

McMahan C.D., Freeborn L.R., Wheeler W.C, Crother B.I. 2015. Forked tongues revisited: molecular apomorphies support morphological hypotheses of squamate evolution. Copeia 103:525–529.

Miller A.K., Kerr A.M., Paulay G., Reich M., Wilson N.G., Carvajal J.I., Rouse G.W. 2017. Molecular phylogeny of extant Holothuroidea (Echinodermata). Mol. Phylogenet. Evol. 111:110–131.

Molloy E.K., Warnow T. 2018. To include or not to include: the impact of gene filtering on species tree estimation methods. Syst. Biol. 67:285–303.

Mongiardino Koch N. 2019. The phylogenomic revolution and its conceptual innovations: a text mining approach. Org. Divers. Evol. 19:99–103.

Mongiardino Koch N., Ceccarelli F.S., Ojanguren-Affilastro A.A., Ramírez M.J. 2017. Discrete and morphometric traits reveal contrasting patterns and processes in the macroevolutionary history of a clade of scorpions. J. Evol. Biol. 30:814–825.

Mongiardino Koch N., Coppard S.E., Lessios H.A., Briggs D.E.G., Mooi R., Rouse G.W. 2018. A phylogenomic resolution of the sea urchin tree of life. BMC Evol. Biol. 18:189.

Mongiardino Koch N., Gauthier J.A. 2018. Noise and biases in genomic data may underlie radically different hypotheses for the position of Iguania within Squamata. PLoS One 13:e0202729.

Mongiardino Koch N., Parry L.A. 2019. Death is on Our Side: Paleontological Data Drastically Modify Phylogenetic Hypotheses. bioRxiv 723882.

Mooi R. 1990a. Living Cassiduloids (Echinodermata, Echinoidea) - A Key And Annotated List. Proc. Biol. Soc. Wash. 103:63–85.

Mooi R. 1990b. Paedomorphosis, Aristotle’s lantern, and the origin of the sand dollars (Echinodermata: Clypeasteroida). Paleobiology 16:25–48.

Mooi R. 1990c. Progenetic miniaturization in the sand dollar Sinaechinocyamus: implications for clypeasteroid phylogeny. In: Echinoderm Research. De Ridder C, Dubois P, Lahaye MC, Jangoux M, Eds. AA Balkema, Rotterdam. pp. 137–143.

Morel B., Kozlov A.M., Stamatakis A. 2018. ParGenes: a tool for massively parallel model selection and phylogenetic tree inference on thousands of genes. Bioinformatics 35:1771–1773.

Mortensen T. 1928. A Monograph of the Echinoidea. I. Cidaroidea. CA Reitzel, Copenhagen.

Mortensen T. 1935. A Monograph of the Echinoidea. II. Bothriocidaroida, Melonechinoida, Lepidocentroida, and Stirodonta. CA Reitzel, Copenhagen.

Mortensen T. 1940. A Monograph of the Echinoidea. III, 1. Aulodonta, with Additions to Vol. II (Lepidocentroida and Stirodonta). CA Reitzel, Copenhagen.

Mortensen T. 1943. A Monograph of the Echinoidea. III, 3. Camarodonta. II. Echinidæ, Strongylocentrotidæ, Parasaleniidæ, Echinometridae. CA Reitzel, Copenhagen.

Mortensen T. 1948a. A Monograph of the Echinoidea. IV, 1 Holectypoida, Cassiduloida. CA Reitzel, Copenhagen.

Mortensen T. 1948b. A Monograph of the Echinoidea. IV, 2. Clypeasteroida. Clypeasteridae, Arachnoidae, Fibulariidae, Laganidae and Scutellidae. CA Reitzel, Copenhagen.

Murrell D.J. 2018. A global envelope test to detect non-random bursts of trait evolution. Methods Ecol. Evol. 9:1739–1748.

Nebelsick J.H. 1996. Biodiversity of shallow-water Red Sea echinoids: implications for the fossil record. J. Mar. Biol. Assoc. UK, 76:185–194.

Nesnidal M.P., Helmkampf M., Bruchhaus I., Hausdorf B. 2010. Compositional heterogeneity and phylogenomic inference of metazoan relationships. Mol. Biol. Evol. 27:2095–2104.

Nguyen L.-T., Schmidt H.A., von Haeseler A., Minh B.Q. 2014. IQ-TREE: a fast and effective stochastic algorithm for estimating maximum-likelihood phylogenies. Mol. Biol. Evol. 32:268–274.

Nosenko T., Schreiber F., Adamska M., Adamski M., Eitel M., Hammel J., Maldonado M., Müller W.E., Nickel M., Schierwater B. 2013. Deep metazoan phylogeny: when different genes tell different stories. Mol. Phylogenet. Evol. 67:223–233.

Nowak M.D., Smith A.B., Simpson C., Zwickl D.J. 2013. A simple method for estimating informative node age priors for the fossil calibration of molecular divergence time analyses. PLoS One 8:e66245.

O’Reilly J.E., Donoghue P.C. 2016. Tips and nodes are complementary not competing approaches to the calibration of molecular clocks. Biol. Lett. 12:20150975.

Pagel M. 1999. Inferring the historical patterns of biological evolution. Nature 401:877–884.

Paradis E., Schliep K. 2018. ape 5.0: an environment for modern phylogenetics and evolutionary analyses in R. Bioinformatics 35:526–528.

Pérez-Portela R., Turon X., Riesgo A. 2016. Characterization of the transcriptome and gene expression of four different tissues in the ecologically relevant sea urchin Arbacia lixula using RNA-seq. Mol. Ecol. Resour. 16:794–808.

Peters R.H. 1986. The ecological implications of body size. Cambridge University Press, Cambridge.

Philip G. 1963. Two Australian Tertiary neolampadids, and the classification of cassiduloid echinoids. Palaeontology 6:718–726.

Philip G. 1965. Classification of echinoids. J. Paleontol. 39:45–62.

Price S.A., Hopkins S.S. 2015. The macroevolutionary relationship between diet and body mass across mammals. Biol. J. Linn. Soc. 115:173–184.

Püschel H.P., O’Reilly J.E., Pisani D., Donoghue P.C. 2020. The impact of fossil stratigraphic ranges on tip-calibration, and the accuracy and precision of divergence time estimates. Palaeontology 63:67–83.

Pyron R.A. 2015. Post-molecular systematics and the future of phylogenetics. Trends Ecol. Evol. 30:384–389.

Quental T.B., Marshall C.R. 2010. Diversity dynamics: molecular phylogenies need the fossil record. Trends Ecol. Evol. 25:434–441.

R Core Team. 2019. R: A Language and Environment for Statistical Computing. R Foundation for Statistical Computing, Vienna. https://www.R-project.org/

Rambaut A., Drummond A.J., Xie D., Baele G., Suchard M.A. 2018. Posterior summarization in Bayesian phylogenetics using Tracer 1.7. Syst. Biol. 67:901–904.

Reich A., Dunn C., Akasaka K., Wessel G. 2015. Phylogenomic analyses of Echinodermata support the sister groups of Asterozoa and Echinozoa. PLoS one 10:e0119627.

Reich M., Stegemann T.R., Hausmann I.M., Roden V.J., Nützel A. 2018. The youngest ophiocistioid: a first Palaeozoic-type echinoderm group representative from the Mesozoic. Palaeontology 61:803–811.

Revell L.J. 2012. phytools: an R package for phylogenetic comparative biology (and other things). Methods Ecol. Evol. 3:217–223.

Robinson D.F., Foulds L.R. 1981. Comparison of phylogenetic trees. Math. Biosci. 53:131–147.

Romiguier J., Gayral P., Ballenghien M., Bernard A., Cahais V., Chenuil A., Chiari Y., Dernat R., Duret L., Faivre N. 2014. Comparative population genomics in animals uncovers the determinants of genetic diversity. Nature 515:261–263.

Ronquist F., Teslenko M., Van Der Mark P., Ayres D.L., Darling A., Höhna S., Larget B., Liu L., Suchard M.A., Huelsenbeck J.P. 2012. MrBayes 3.2: efficient Bayesian phylogenetic inference and model choice across a large model space. Syst. Biol. 61:539–542.

Rose E.P.F. 1982. Holectypoid echinoids and their classification. In: International Echinoderm Conference, Tampa Bay. Lawrence JM, Ed. AA Balkema:Rotterdam. pp. 145–152.

Ryan J.F. 2014. Alien Index: identify potential non-animal transcripts or horizontally transferred genes in animal transcriptomes. doi:10.5281/zenodo.21029.

Salichos L., Rokas A. 2013. Inferring ancient divergences requires genes with strong phylogenetic signals. Nature 497:327–331.

Sanderson M.J. 2002. Estimating absolute rates of molecular evolution and divergence times: a penalized likelihood approach. Mol. Biol. Evol. 19:101–109.

Sayyari E., Mirarab S. 2016. Fast coalescent-based computation of local branch support from quartet frequencies. Mol. Biol. Evol. 33:1654–1668.

Schliep K.P. 2011. phangorn: phylogenetic analysis in R. Bioinformatics 27:592.

Schultz H.A. 2015. Echinoidea: with pentameral symmetry. Walter de Gruyter GmbH, Berlin.

Seilacher A. 1979. Constructional morphology of sand dollars. Paleobiology 5:191–221.

Sharma P.P., Kaluziak S.T., Pérez-Porro A.R., González V.L., Hormiga G., Wheeler W.C., Giribet G. 2014. Phylogenomic interrogation of Arachnida reveals systemic conflicts in phylogenetic signal. Mol. Biol. Evol. 31:2963–2984.

Simion P., Belkhir K., François C., Veyssier J., Rink J.C., Manuel M., Philippe H., Telford M.J. 2018. A software tool ‘CroCo’detects pervasive cross-species contamination in next generation sequencing data. BMC Biol. 16:28.

Simmons M.P., Gatesy J. 2016. Biases of tree-independent-character-subsampling methods. Mol. Phylogenet. Evol. 100:424–443.

Simpson G.G. 1944. Tempo and mode in evolution. Columbia University Press, New York.

Slater G.J, Harmon L.J. 2013. Unifying fossils and phylogenies for comparative analyses of diversification and trait evolution. Methods Ecol. Evol. 4:699–702.

Smith A.B. 1990. Echinoid evolution from the Triassic to the Lower Jurassic. Cah. Univ. Cath. Lyon Ser. Sci. 3:79–117.

Smith A.B., Hollingworth N. 1990. Tooth structure and phylogeny of the Upper Permian echinoid Miocidaris keyserlingi. Proc. Yorks. Geol. Soc. 48:47–60.

Smith A.B, Littlewood D.T.J., Wray G. 1995. Comparing patterns of evolution: larval and adult life history stages and ribosomal RNA of post-Palaeozoic echinoids. Phil. Trans. R. Soc. Lond. B, 349:11–18.

Smith A.B. 1980. Stereom microstructure of the echinoid test. Paleontology 25:1–81.

Smith A.B. 1981. Implications of lantern morphology for the phylogeny of post-Palaeozoic echinoids. Palaeontology 24:779–801.

Smith A.B. 1984. Echinoid palaeobiology. George Allen & Unwin, London.

Smith A.B. 1997. Echinoderm phylogeny: how congruent are morphological and molecular estimates? Paleontol. Soc. Pap. 3:337–355.

Smith A.B. 2001. Probing the cassiduloid origins of clypeasteroid echinoids using stratigraphically restricted parsimony analysis. Paleobiology 27:392–404.

Smith A.B. 2007. Intrinsic versus extrinsic biases in the fossil record: contrasting the fossil record of echinoids in the Triassic and early Jurassic using sampling data, phylogenetic analysis, and molecular clocks. Paleobiology 33:310–323.

Smith A.B. 2015. British Jurassic Regular Echinoids. Part 1: Introduction, Cidaroida, Echinothurioida, Aspidodiadematoida and Pedinoida. Palaeontographical Society, London.

Smith A.B., Jeffery C.H. 1998. Selectivity of extinction among sea urchins at the end of the Cretaceous period. Nature 392:69–71.

Smith AB, Kroh A. 2013. Phylogeny of sea urchins. In: Sea Urchins: Biology and Ecology, 3rd edition. Lawrence JM, Ed. Academic Press, Cambridge. pp. 1–14.

Smith A.B., Pisani D., Mackenzie-Dodds J.A., Stockley B., Webster B.L., Littlewood D.T.J. 2006. Testing the molecular clock: molecular and paleontological estimates of divergence times in the Echinoidea (Echinodermata). Mol. Biol. Evol. 23:1832–1851.

Smith F.A., Lyons S.K. 2013. Animal body size: linking pattern and process across space, time, and taxonomic group. University of Chicago Press, Chicago.

Smith S.A., Brown J.W., Walker J.F. 2018. So many genes, so little time: A practical approach to divergence-time estimation in the genomic era. PLoS One 13:e0197433.

Souto C., Mooi R., Martins L., Menegola C., Marshall C.R. 2019. Homoplasy and extinction: the phylogeny of cassidulid echinoids (Echinodermata). Zool. J. Linn. Soc. 187:622–660.

Steneck R.S. 2013. Sea urchins as drivers of shallow benthic marine community structure. In: Sea Urchins: Biology and Ecology, 3rd edition. Lawrence JM, Ed. Academic Press, Cambridge. pp. 195–212.

Steneck R.S., Graham M.H., Bourque B.J., Corbett D., Erlandson J.M., Estes J.A., Tegner M.J. 2002. Kelp forest ecosystems: biodiversity, stability, resilience and future. Environ. Conserv. 29:436–459.

Stockley B., Smith A.B., Littlewood D.T.J., Lessios H.A., Mackenzie-Dodds J.A. 2005. Phylogenetic relationships of spatangoid sea urchins (Echinoidea): taxon sampling density and congruence between morphological and molecular estimates. Zool. Scr. 34:447–468.

Suter S.J. 1988. The decline of the cassiduloids: merely bad luck. In: Echinoderm biology. Burke RD, Mladenov PV, Lambert P, Parsley RL, Eds. AA Balkema, Rotterdam. pp. 91–95.

Suter S.J. 1994. Cladistic analysis of cassiduloid echinoids: trying to see the phylogeny for the trees. Biol. J. Linn. Soc. 53:31–72.

Talavera G., Castresana J. 2007. Improvement of phylogenies after removing divergent and ambiguously aligned blocks from protein sequence alignments. Syst. Biol. 56:564–577.

Telford M., Harold A.S., Mooi R. 1983. Feeding structures, behavior, and microhabitat of Echinocyamus pusillus (Echinoidea: Clypeasteroida). Biol. Bull. 165:745–757.

Thayer C.W. 1983. Sediment-mediated biological disturbance and the evolution of marine benthos. In: Biotic interactions in recent and fossil benthic communities. Tevesz MJS, McCall PL, Eds. Springer, Boston. pp. 479–625.

Thompson J.R., Ausich W.I. 2016. Facies distribution and taphonomy of echinoids from the Fort Payne Formation (late Osagean, early Viséan, Mississippian) of Kentucky. J. Paleont. 90:239–249.

Thompson J.R., Erkenbrack E.M., Hinman V.F., McCauley B.S., Petsios E., Bottjer D.J. 2017. Paleogenomics of echinoids reveals an ancient origin for the double-negative specification of micromeres in sea urchins. Proc. Natl. Acad. Sci. 114:5870–5877.

Thompson J.R., Hu S.-x., Zhang Q.-Y., Petsios E., Cotton L.J., Huang J.-Y., Zhou C.-y., Wen W., Bottjer D.J. 2018. A new stem group echinoid from the Triassic of China leads to a revised macroevolutionary history of echinoids during the end-Permian mass extinction. R .Soc. Open Sci. 5:171548.

Thompson J.R., Mirantsev G.V., Petsios E., Bottjer D.J. 2019. Phylogenetic analysis of the Archaeocidaridae and Palaeozoic Miocidaridae (Echinodermata, Echinoidea) and the origin of crown group echinoids. Pap. Palaeontol. doi:10.1002/spp2.1280.

Thompson J.R., Petsios E., Davidson E.H., Erkenbrack E.M., Gao F., Bottjer D.J. 2015. Reorganization of sea urchin gene regulatory networks at least 268 million years ago as revealed by oldest fossil cidaroid echinoid. Sci. Rep. 5:15541.

Thuy B., Hagdorn H., Gale A.S. 2017. Paleozoic echinoderm hangovers: Waking up in the Triassic. Geology 45:531–534.

Twitchett R.J., Oji T. 2005. Early Triassic recovery of echinoderms. C. R. Palevol, 4:531–542.

Uthicke S., Ebert T., Liddy M., Johansson C., Fabricius K.E., Lamare M. 2016. Echinometra sea urchins acclimatized to elevated pCO 2 at volcanic vents outperform those under present-day pCO 2 conditions. Global Change Biol. 22:2451–2461.

Valentine J.F., Heck Jr K.L. 1991. The role of sea urchin grazing in regulating subtropical seagrass meadows: evidence from field manipulations in the northern Gulf of Mexico. J. Exp. Mar. Biol. Ecol. 154:215–230.

Wagner C., Durham J. 1966. Holectypoids. In: Treatise on Invertebrate Paleontology, Part U. Moore RC, Ed. University of Kansas Press, Lawrence. pp. 440–450.

Wang H.-C., Minh B.Q., Susko E., Roger A.J. 2017. Modeling site heterogeneity with posterior mean site frequency profiles accelerates accurate phylogenomic estimation. Syst. Biol. 67:216–235.

Warren D.L., Geneva A.J., Lanfear R. 2017. RWTY (R We There Yet): An R package for examining convergence of Bayesian phylogenetic analyses. Mol. Biol. Evol. 34:1016–1020.

Whelan N.V., Kocot K.M., Moroz L.L., Halanych K.M. 2015. Error, signal, and the placement of Ctenophora sister to all other animals. Proc. Natl. Acad. Sci. 112:5773–5778.

Wiens J.J., Kuczynski C.A., Townsend T., Reeder T.W., Mulcahy D.G., Sites Jr J.W. 2010. Combining phylogenomics and fossils in higher-level squamate reptile phylogeny: molecular data change the placement of fossil taxa. Syst. Biol., 59:674–688.

Woodward S. 1863. On Echinothuria floris, a new and anomalous echinoderm from the Chalk of Kent. The Geologist, 6:327–330.

Zhang C., Rabiee M., Sayyari E., Mirarab S. 2018. ASTRAL-III: polynomial time species tree reconstruction from partially resolved gene trees. BMC Bioinformatics 19:153.

Ziegler A., Lenihan J., Zachos L.G., Faber C., Mooi R. 2016. Comparative morphology and phylogenetic significance of Gregory’s diverticulum in sand dollars (Echinoidea: Clypeasteroida). Org. Divers. Evol. 16:141–166.

Ziegler A., Stock S.R., Menze B.H., Smith A.B. 2012. Macro-and microstructural diversity of sea urchin teeth revealed by large-scale mircro-computed tomography survey. In: Developments in X-ray Tomography VIII. Stock SR, Ed. International Society for Optics and Photonics, San Diego. pp. 85061G.

